# Cyclin D-CDK4/6 couple proliferation to membrane and mitochondrial protein supply through a DeSI1 phospho-switch

**DOI:** 10.64898/2026.09.16.752001

**Authors:** Sharon Kaisari, Ethan Lane, Raymond Wang, Jeffrey Estrada, Marta Collu, Daniele Simoneschi, Yeon-Tae Jeong, Mengxi Liu, Qingyue Zhang, Yuki Kito, Ifat Abramovich, Jason Yin, Tori Rodrick, Gyles Ward, Tyler Cropley, Adriana Heguy, Sang Yong Kim, Ran Brosh, Herman Wolosker, Namrata D. Udeshi, Steven A. Carr, Cynthia A. Loomis, Feng-Xia Liang, Beatrix Ueberheide, Drew Jones, Ning Zheng, Michele Pagano

**Affiliations:** Department of Biochemistry and Molecular Pharmacology, NYU Grossman School of Medicine, New York, NY 10016, USA; Howard Hughes Medical Institute, NYU Grossman School of Medicine, New York, NY 10016, USA; Laura and Isaac Perlmutter Metabolomics Center, The Ruth and Bruce Rappaport Faculty of Medicine, Technion–Israel Institute of Technology, Haifa 3525433, Israel; Microscopy Laboratory, Division of Advanced Research Technologies, NYU Grossman School of Medicine, New York, NY 10016, USA; Metabolomics Core Resource Laboratory, NYU Grossman School of Medicine, New York, NY 10016, USA; Experimental Pathology Research Laboratory, Division of Advanced Research Technologies, NYU Grossman School of Medicine, New York, NY 10016, USA; Proteomics Laboratory, Division of Advanced Research Technologies, NYU Grossman School of Medicine, New York, NY 10016, USA; Genome Technology Center, Division of Advanced Research Technologies, NYU Langone Health, New York, NY 10016, USA; Department of Pathology and Laboratory Medicine, Miller School of Medicine, University of Miami, Miami, FL 33136, USA; Advanced Rodent Transgenics Laboratory, Division of Advanced Research Technologies, NYU Grossman School of Medicine, New York, NY 10016, USA; Department of Biochemistry, The Ruth and Bruce Rappaport Faculty of Medicine, Technion–Israel Institute of Technology, Haifa 3525433, Israel; Broad Institute of MIT and Harvard, Cambridge, MA 02142, USA; Department of Pharmacology, University of Washington, Seattle, WA 98195, USA; Howard Hughes Medical Institute, University of Washington, Seattle, WA 98195, USA

## Abstract

Cyclin D-CDK4/6 complexes drive cell-cycle entry through an RB-E2F-dependent transcriptional program, but how they coordinate proliferation with the membrane and organelle protein supply required for growth is unclear. We identify DeSI1 as a cyclin D-CDK4/6 substrate whose phosphorylation at S25 converts a latent homodimer into an active monomeric deubiquitylase that recognizes hydrophobic proteins such as those bearing transmembrane domains and mitochondrial targeting sequences. Phosphorylated, active DeSI1 extends the lifetime of newly synthesized hydrophobic proteins that support membrane and mitochondrial capacity. Constitutive DeSI1 activation uncouples this proteostatic program from metabolic supply, creating a cytidine-nucleotide supply-demand imbalance associated with impaired CTP-dependent phospholipid homeostasis, cardiolipin depletion, and mitochondrial decompensation — defects that cytidine reverses. In mice, constitutive DeSI1 activation causes progressive cerebellar degeneration with membrane-protein accumulation, respiratory-chain loss, and phospholipid depletion. These findings define a post-translational mechanism coupling cell-cycle entry to membrane and mitochondrial capacity, and reveal the cost of uncoupling this program from its metabolic support.

**Graphical Abstract:** 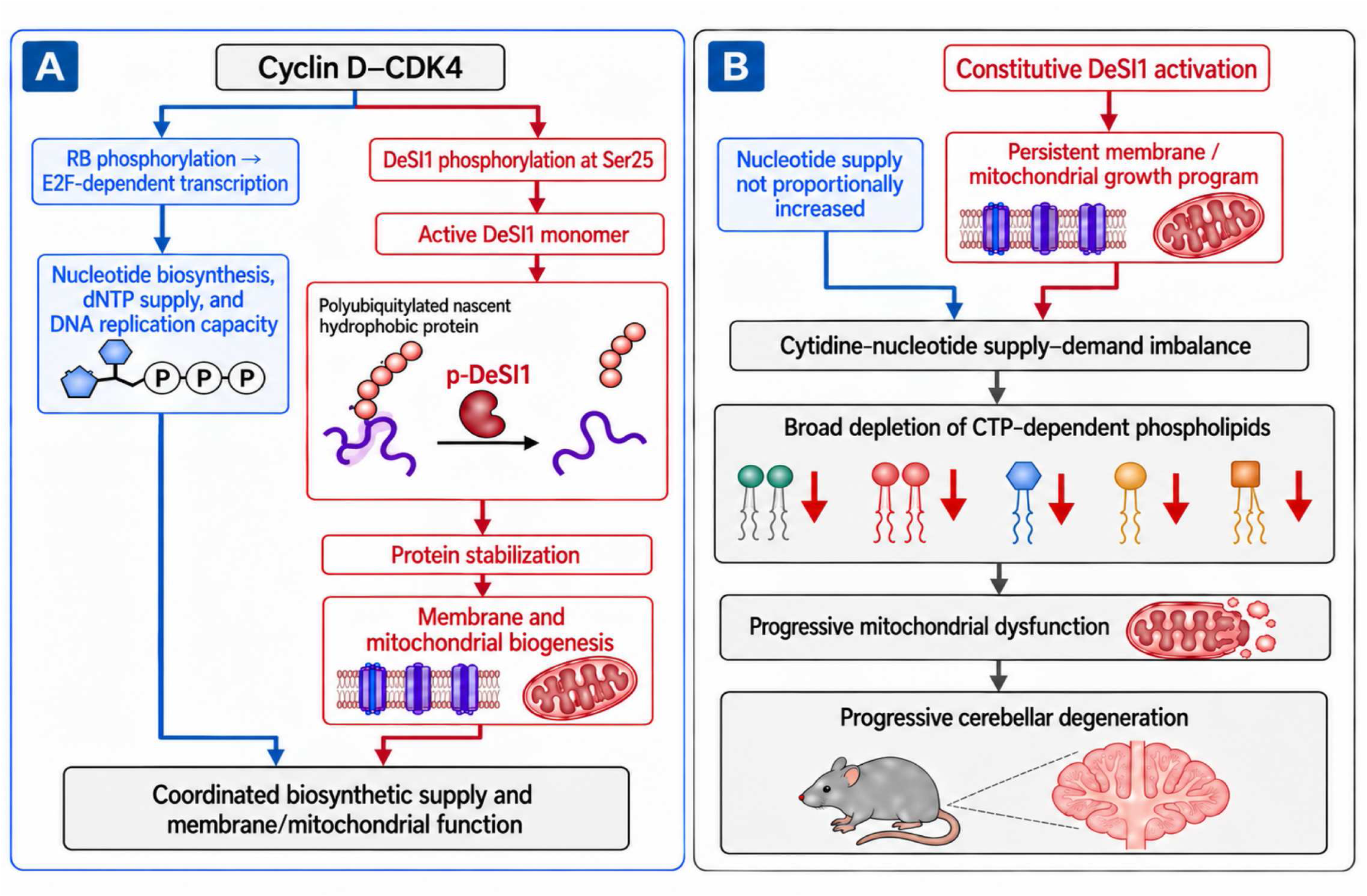

## Introduction

Mammalian D-type cyclins (D1, D2, and D3) assemble with CDK4 or CDK6 to form a family of cyclin D-CDK4/6 kinase complexes^1,2^. Although these complexes share core roles in promoting cell-cycle entry, their composition varies by cell type, developmental context, and upstream signaling environment, suggesting both overlapping and context-specific functions. Cyclin D-CDK4/6 complexes link mitogenic signals to progression through G1 primarily through phosphorylation of the retinoblastoma protein (RB) family^1,2^. Through phosphorylation and functional inactivation of RB and its paralogs RBL1 and RBL2, cyclin D-CDK4/6 enable E2F-dependent transcriptional programs that not only prime cells for S-phase entry but also increase the metabolic and biosynthetic capacity necessary to sustain proliferation^3,4,5^. Despite the importance of these kinases, few physiologically validated substrates outside the RB family have been linked to coherent cellular programs, even though functional evidence suggests that CDK4/6 has a broader repertoire^6,7^. This gap is significant because entry into the cell cycle entails more than preparing for DNA replication: cell growth must be coordinated with division, and proliferating cells must double their proteome, membranes, and organelles before they divide^8^. Whether cyclin D-CDK4/6 complexes help orchestrate these biosynthetic changes remains poorly understood.

Hydrophobic proteins destined for membranes, such as proteins containing transmembrane domains (TMDs) or amphipathic mitochondrial targeting sequences (MTSs), are intrinsically challenging for protein biogenesis and are subject to tight chaperone- and ubiquitin-dependent quality control^9,10^. The BAG6 complex, ubiquilins, and VCP/p97 are part of this surveillance system^11,12,13^. Importantly, proteolysis is not necessarily the endpoint of this quality-control process: delaying or antagonizing degradation can provide hydrophobic clients another opportunity for productive targeting to membranes^14^.

Mitogenic stimulation enhances protein synthesis and thereby places greater demands on protein quality-control systems. However, membrane and organelle expansion during growth increases the demand for membrane-targeted and mitochondrial proteins, creating a proteostatic pressure that must be buffered to allow these proteins to accumulate and be productively targeted rather than routed toward degradation.

Here, we identified DeSI1 as a previously unknown substrate of cyclin D-CDK4/6 and found that its phosphorylation activates its deubiquitylase activity and counteracts the proteostatic pressure for a defined class of hydrophobic and membrane-targeted proteins, thereby reducing their ubiquitin-dependent turnover and promoting their accumulation during G1. This pathway defines a second output of cyclin D-CDK4/6 that operates in parallel with E2F: whereas the RB-E2F arm increases nucleotide supply, the DeSI1 arm increases membrane-targeted and mitochondrial protein supply. The results of these studies are presented below.

## Results

### DeSI1 is a cyclin D1-CDK4 substrate

To identify physiologically relevant CDK4/6 substrates, we leveraged an HCT-116 cell system in which the mini-auxin-inducible degron (mAID) is fused to endogenous AMBRA1, a substrate receptor of a CUL4-RING E3 ubiquitin Ligase (CRL4) that targets all D-type cyclins for proteasomal degradation^15,16^. Treatment with auxin in three independent experiments induced rapid AMBRA1 depletion, which in turn resulted in the accumulation of endogenous cyclin D1 and increased phosphorylation of RB at serine 780, confirming CDK4/6 activation (Figure 1A). RB phosphorylation was inhibited by the selective CDK4/6 inhibitor palbociclib (Figure 1A). The three samples were also used for quantitative tandem mass tag (TMT) phosphoproteomics to capture phosphorylation events that occur upon acute accumulation of cyclin D1. Phosphoproteomic analysis revealed DeSI1 (AKA PPPDE2 and FAM152B), a small PPPDE-family cysteine protease with isopeptidase activity^17,18^, as the second most significantly enriched phosphoprotein following cyclin D1 accumulation, with phosphorylation at serine 25 (S25) showing even greater fold-change than canonical CDK4/6 substrates RB and RBL2 (Figure 1B and Supplementary Figure 1C). DeSI1 phosphorylation increased over 4-fold upon AMBRA1 depletion, an effect completely reversed by palbociclib treatment, confirming its CDK4/6 dependence. Global proteomic analysis verified that these phosphorylation changes occurred without alterations in its total protein abundance (Supplementary Figure 1A). These findings were independently validated in a second set of three independent experiments analyzed by a different proteomic facility where the top-ranking phospho-peptide corresponded to a DeSI1 phospho-S25 (p-S25) containing peptide (Figure 1C and Supplementary Figure 1B,C).

**Figure 1.**
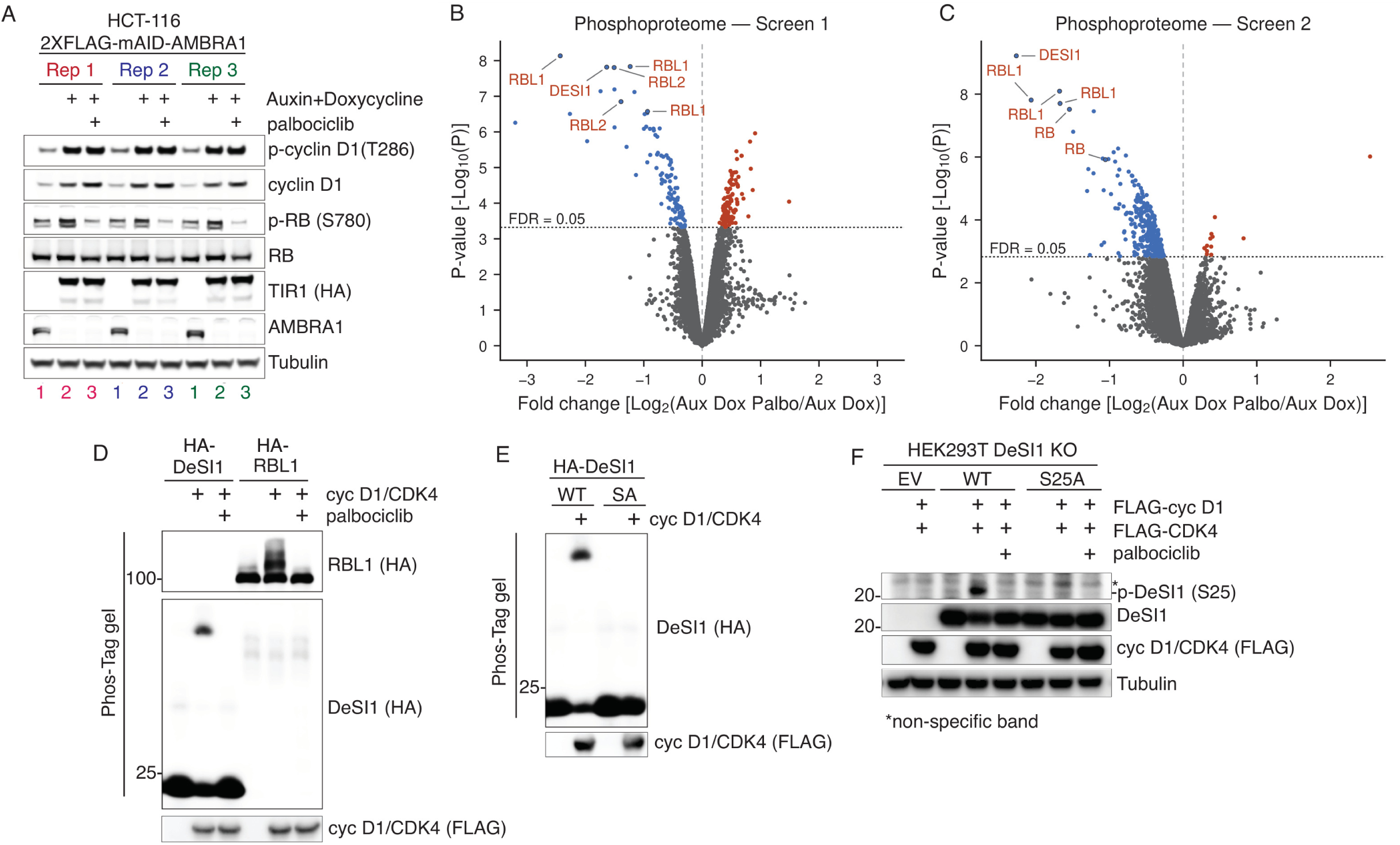
DeSI1 is a cyclin D1-CDK4 substrate. **(A)** Validation of the cyclin D accumulation system. HCT-116 2×FLAG-mAID-AMBRA1 cells were treated with auxin and doxycycline to degrade AMBRA1, ± palbociclib, in three biological replicates. Whole-cell extracts were immunoblotted for the indicated proteins. See Materials and Methods. **(B)** Phosphoproteomic Screen 1. Volcano plot showing phosphorylation changes following acute CDK4/6 inhibition in AMBRA1-depleted HCT-116 cells, expressed as log₂(Aux+Dox+palbociclib/Aux+Dox). Statistical significance was assessed using a two-sided two-sample moderated *t*-test with empirical-Bayes variance shrinkage (limma/Protigy). DeSI1 and representative RB-family phosphosites are indicated. Dashed line denotes Benjamini–Hochberg FDR = 0.05. **(C)** Independent phosphoproteomic Screen 2 performed using the same experimental comparison. Statistical significance was assessed using the two-sided two-sample *t*-test implemented in Perseus. DeSI1 and representative RB-family phosphopeptides are indicated. Dashed line denotes Benjamini–Hochberg FDR = 0.05. **(D)** Phos-tag validation of DeSI1 phosphorylation. HEK293T cells expressing HA-DeSI1 or HA-p107 (control), ± cyclin D1–CDK4 and ± palbociclib, resolved by Phos-tag SDS–PAGE. See Materials and Methods. **(E)** Phosphorylation-site mapping. HA-DeSI1 wild-type (WT) or S25A (SA) expressed with cyclin D1–CDK4; the S25A substitution abolishes the mobility shift. **(F)** Specificity of the pDeSI1 (S25) antibody. HEK293T DeSI1-knockout cells expressing empty vector (EV), DeSI1 WT or S25A together with FLAG-cyclin D1 and FLAG-CDK4, ± palbociclib. See Materials and Methods.

To validate DeSI1 as a cyclin D1-CDK4 substrate, we employed multiple biochemical approaches. Using Phos-tag gel electrophoresis, which enhances resolution of phosphorylated protein isoforms, we observed a mobility shift of HA-tagged DeSI1 upon co-expression with cyclin D1-CDK4 in HEK293T cells (Figure 1D and Supplementary Figure 1D). This phosphorylation-induced shift was comparable to that observed for RBL1 and was inhibited by palbociclib treatment (Figure 1D), demonstrating CDK4/6 dependency. Mutation of S25 to alanine (S25A) abolished the mobility shift, confirming S25 as the primary CDK4/6 phosphorylation site (Figure 1E).

We also generated an antibody against a phospho-peptide corresponding to the region of DeSI1 containing p-S25. When wild-type DeSI1, but not DeSI1(S25A), was co-expressed with cyclin D1 and CDK4, the antibody detected a band that was absent following treatment with palbociclib (Figure 1F). Antibody specificity was further validated in DeSI1-knockout HEK293T cells (Figure 1F). Finally, the phospho-specific antibody also recognized DeSI1(S25E) purified from either HEK293T cells or bacteria (Supplementary Figure 1E).

DeSI1 S25 phosphorylation was also observed when the protein was co-expressed with cyclin D2-CDK4 or cyclin D3-CDK4, but not with other cyclin-CDK complexes (Supplementary Figure 1F).

Finally, using purified recombinant proteins, we performed an in vitro phosphorylation assay with cyclin D1-CDK4 and confirmed that DeSI1 is directly phosphorylated by cyclin D1-CDK4, similarly to RB (Supplementary Figure 1G).

### S25 phosphorylation enables DeSI1 monomerization and deubiquitylase function

Having established DeSI1 as a cyclin D1-CDK4 substrate, we next asked how phosphorylation at S25 affects DeSI1 function. The published crystal structure of DeSI1^18^ revealed that the protein forms a homodimer, with its N-terminal amphipathic helix, which harbors S25, contributing directly to the dimer interface. The location of S25 therefore suggests that its phosphorylation might regulate DeSI1 oligomerization. To test this possibility, we purified recombinant wild-type DeSI1 and the phosphomimetic DeSI1(S25E) mutant and analyzed their oligomeric states by size-exclusion chromatography. Wild-type DeSI1 eluted at ∼36 kDa, consistent with a homodimer, whereas DeSI1(S25E) eluted at ∼18 kDa, consistent with a monomeric species (Figure 2A). Consistent with these findings, phosphorylation of DeSI1 by cyclin D1-CDK4 shifted the protein into monomer-sized fractions, with the p-DeSI1 signal confined to these later-eluting fractions (Figure 2B).

**Figure 2.**
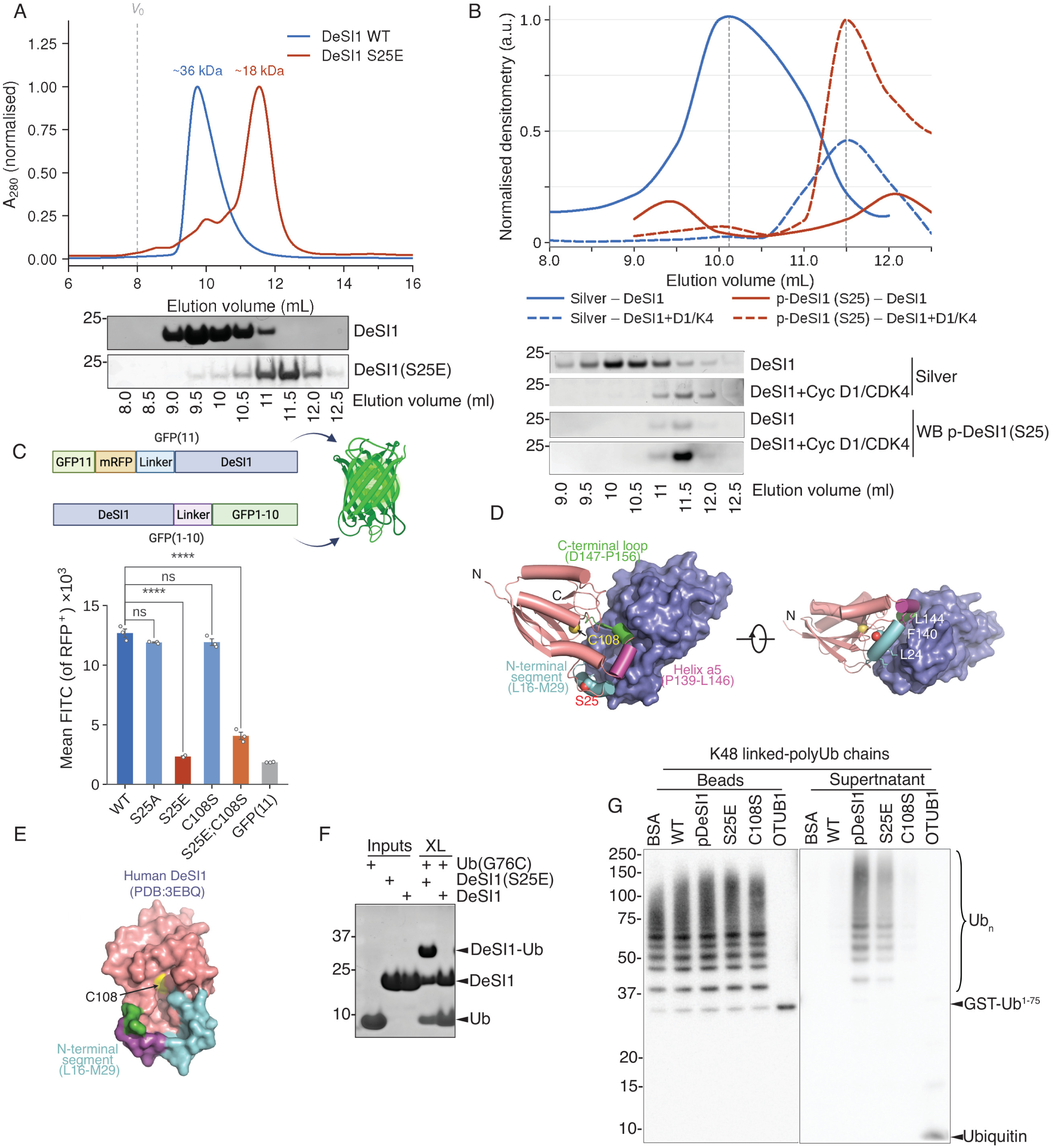
CDK4-mediated S25 phosphorylation monomerizes DeSI1 and activates a latent deubiquitylase function. **(A)** Size-exclusion chromatography of recombinant wild-type and S25E DeSI1 purified from E. coli (Superdex 75 Increase 10/300 GL). Absorbance at 280 nm is scaled to the apex of each trace; the dashed line marks the column void volume (V₀ = 8.0 mL). Wild-type elutes at 9.75 mL (∼36 kDa, dimer) and S25E at 11.54 mL (∼18 kDa, monomer), assigned from a globular protein standard curve. Below, immunoblots of the indicated fractions. **(B)** DeSI1 was immunopurified from HEK293F cells expressing DeSI1 alone or together with Cyclin D1–CDK4 (D1K4), eluted natively with Flag peptide and resolved by size-exclusion chromatography. Upper, densitometry of total DeSI1 (silver stain) and p-DeSI1 (S25) immunoreactivity across fractions; within each readout both preparations are scaled to the shared maximum, so traces are comparable within but not between readouts. Markers denote measured fractions; curves are shape-preserving interpolations. Dashed lines mark the dimer (∼36 kDa) and monomer (∼18 kDa) elution volumes from (A). Lower, corresponding silver stains and p-DeSI1 (S25) immunoblots. Monomer fractions from the Cyclin D1–CDK4 condition were pooled as the p-DeSI1 preparation used in (G). **(C)** Split-GFP complementation in U2OS DeSI1-knockout cells. Upper, construct design. Lower, mean FITC intensity of RFP-positive cells for the indicated DeSI1 variants; GFP(11) alone is the fragment-only negative control and was not included in the statistical comparisons. Mean ± SEM with individual replicates overlaid, n = 3 biological replicates. One-way ANOVA (F(5,12) = 548.9, P < 0.0001) with Bonferroni correction, each variant against wild-type; ****P < 0.0001, ns not significant. **(D)** Structure of the DeSI1 homodimer and organization of the dimer interface. One DeSI1 protomer is shown in salmon and the second protomer in blue. The three structural elements contributing to the dimer interface are highlighted on one protomer: the N-terminal segment containing S25 (L16–M29, cyan), helix α5 (P139–L146, magenta), and the C-terminal loop (D147–P156, green). The catalytic cysteine C108 and S25 are indicated. Right, a rotated view highlights representative interface residues L20, L24, F140, and L144. Residue-level detail is shown in Supplementary Figure 2. **(E)** Surface representation of the human DeSI1 crystal structure (PDB 3EBQ) in the monomeric state. The N-terminal interface segment (L16–M29) is highlighted, and the catalytic cysteine C108 is indicated within the exposed catalytic pocket. **(F)** Disulfide crosslinking of ubiquitin G76C to DeSI1 C58A variants, resolved under non-reducing conditions. Inputs and crosslinking reactions (XL) are shown; the DeSI1–ubiquitin adduct forms with S25E only. Size-exclusion analysis of these reactions in Supplementary Figure 2. **(G)** Deubiquitylase assay on GST-Ub^1–75^–anchored K48-linked polyubiquitin chains. Bead-bound and supernatant fractions were immunoblotted for ubiquitin after incubation with BSA, wild-type, p-DeSI1, S25E or C108S DeSI1, or OTUB1 as a K48-specific positive control. Ub_n_, polyubiquitin. The K63-linked counterpart is in Supplementary Figure 2.

A similar behavior was observed in cells. In a split-GFP complementation assay using U2OS *DeSI1*-/- cells, wild-type DeSI1, DeSI1(S25A), and the catalytically inactive DeSI1(C108S) mutant efficiently reconstituted GFP fluorescence, whereas DeSI1(S25E) and the double mutant DeSI1(S25E;C108S) failed to do so (Figure 2C). The fact that DeSI1(C108S) behaves as wild-type DeSI1 indicates that DeSI1 catalytic activity is dispensable for homodimerization.

The DeSI1 dimer interface is formed by three structural elements (PDB 2WP7)^18^: the N-terminal segment containing S25 (L16–M29), helix α5 (P139–L146), and the C-terminal loop (D147–P156) (Figure 2D and Supplementary Figure 2A). Mutations targeting the first two elements (L20E;L24E and L144R) yielded predominantly monomeric species by size-exclusion chromatography, whereas F140S produced a heterogeneous profile consistent with a dynamic monomer-dimer equilibrium (Supplementary Figure 2B). In contrast, deletion of residues 151-163 did not disrupt dimerization. Thus, dimer stability depends primarily on the N-terminal amphipathic helix and helix-turn-helix motif, placing S25 within a key dimerization element. Notably, the catalytic pocket containing C108 is buried within the dimer interface. In the monomeric structure (PDB 3EBQ), C108 is exposed (Figure 2E), suggesting that monomerization may promote an accessible, catalytically competent state.

We directly tested active-site accessibility by disulfide crosslinking. Under non-reducing conditions, ubiquitin(G76C), which places a cysteine at the scissile position, formed a 1:1 disulfide-linked adduct with DeSI1(S25E), but not wild-type DeSI1 (Figure 2F and Supplementary Figure 2H). Thus, the catalytic cysteine is accessible to ubiquitin in monomeric, but not dimeric, DeSI1.

Activity-based probe labeling further supported this model and established the modifier specificity of DeSI1. These probes consist of ubiquitin, SUMO, or NEDD8 conjugated to an electrophilic warhead (vinyl sulfone [VS] or vinyl methyl ester [VME]) that covalently traps the catalytic cysteine of active cysteine proteases. Only DeSI1(S25E) formed a covalent adduct with ubiquitin-VS, whereas wild-type DeSI1 and the other DeSI1 mutants were not labeled (Supplementary Figure 2C). None of the DeSI1 variants reacted with SUMO1-VS, SUMO2-VME (vinyl methyl ester), or NEDD8-VS, nor did they cleave SUMO2 polychains, despite robust labeling and cleavage by the respective positive controls (Supplementary Figure 2D-G). Thus, DeSI1 is a ubiquitin-directed enzyme and its catalytic reactivity is gated by S25 phosphorylation.

Finally, we assayed deubiquitylase activity using resin-anchored, K48-linked polyubiquitin chains assembled on GST-ubiquitin(1–75). Because GST-ubiquitin(1–75) cannot be cleaved, the release of ubiquitin chains or free ubiquitin into the supernatant directly reports deubiquitylase activity. Both phosphorylated DeSI1 and DeSI1(S25E) efficiently released intact polyubiquitin chains from substrates, whereas wild-type DeSI1 and the catalytically inactive DeSI1(C108S) mutant did not (Figure 2G and Supplementary Figure 2I). In contrast, K63-linked chains were refractory to all DeSI1 variants (Supplementary Figure 2J). Thus, S25 phosphorylation seems to convert DeSI1 into an active, K48-selective en bloc deubiquitylase that cleaves the substrate-ubiquitin linkage while largely preserving Ub-Ub linkages within the chain, suggesting that DeSI1 is specialized for substrate deubiquitylation and stabilization rather than ubiquitin-chain editing.

### Phosphorylated DeSI1 preferentially associates with and promotes the stabilization of hydrophobic and membrane-targeted proteins

Next, we asked which cellular proteins interact with DeSI1 and whether S25 phosphorylation regulates these interactions and possibly substrate deubiquitylation. We addressed these questions using two orthogonal approaches (Supplementary Figure 3A).

In the first approach, we expressed wild-type DeSI1 or DeSI1(S25E) in HEK293T cells and performed immunoprecipitation followed by mass spectrometry across three independent biological replicates. Wild-type DeSI1 associated primarily with the ubiquitin-proteasome and protein quality-control machinery. Prominent WT-enriched interactors included proteins of the BAG6 complex (*i.e.,* BAG6, UBL4A, GET4, and ASNA1), ubiquilins (UBQLN1, 2, and 4), VCP (AKA p97) and its cofactors NPL4 and UFD1, HSP70-family chaperones, ubiquitin (UBC and UBA52), and several proteasome subunits (Figure 3A, Supplementary Figure 3B, and Supplementary Table 1). In contrast, the DeSI1(S25E)-enriched interactors were dominated by membrane-associated and mitochondrial proteins. These included mitochondrial carriers, mitochondrial import and respiratory proteins, ER translocon and targeting components, plasma-membrane ion-transport proteins, and enzymes involved in lipid and sterol metabolism (Figure 3A, Supplementary Figure 3C, and Supplementary Table 1).

**Figure 3.**
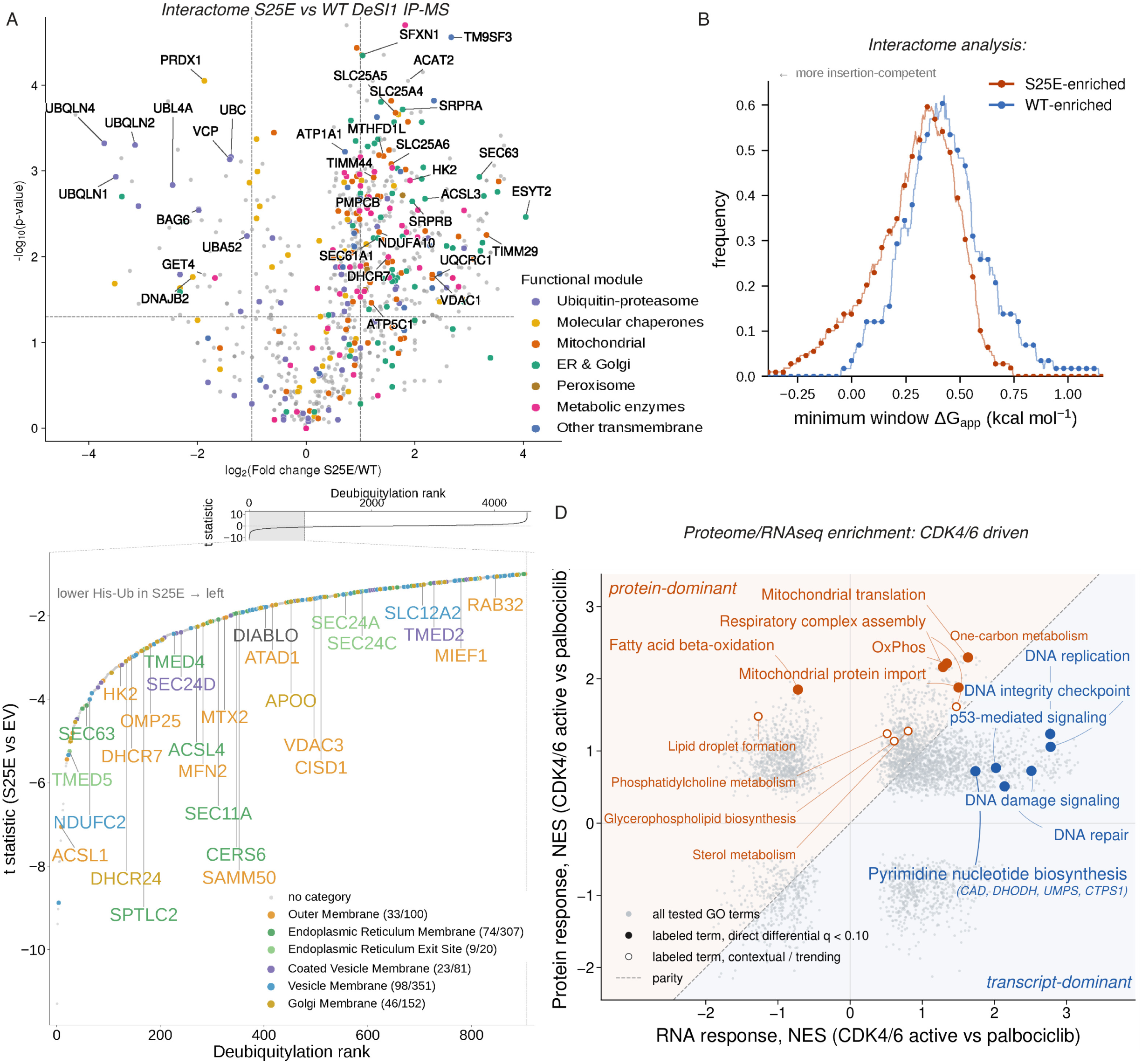
Phosphorylated DeSI1 preferentially interacts with and promotes the stabilization of hydrophobic and membrane-targeted proteins. **(A)** Phosphorylation-dependent DeSI1 interactome. Volcano plot of proteins recovered by FLAG–DeSI1 immunoprecipitation followed by mass spectrometry in HEK293T cells expressing DeSI1(S25E) versus wild-type DeSI1. The x axis shows log₂ fold change (S25E/WT) and the y axis −log₁₀(P). Points are colored according to the indicated functional modules; grey, unassigned proteins. Dashed lines indicate |log₂ fold change| = 1 and nominal P = 0.05; P values were derived from the regularized t statistic described in Methods. Representative WT-enriched and S25E-enriched interactors are labeled. n = 3 biological replicates. **(B)** S25E-enriched interactors are biased toward membrane-insertion-prone sequences. Moving-window frequency distributions of minimum-window ΔG_app for S25E-enriched interactors (n = 109) and WT-enriched interactors (n = 58). For each protein, the Hessa biological hydrophobicity scale was summed across overlapping 19-residue windows and the minimum value was retained; lower values indicate greater membrane-insertion propensity. S25E-enriched interactors were shifted toward lower minimum-window ΔG_app values than WT-enriched interactors (median 0.336 versus 0.420 kcal mol−1; one-sided Mann–Whitney P = 2.1 × 10−5; rank-biserial correlation = −0.385). Ten of 109 S25E-enriched proteins (9.2%) had minimum-window ΔG_app ≤ 0, compared with 0 of 58 WT-enriched proteins. Group definition and matched-permutation sensitivity analyses are described in Methods. **(C)** His-ubiquitin pulldown identifies a membrane-enriched S25E deubiquitylation tail. HEK293F cells expressing 6×His–ubiquitin together with empty vector (EV), DeSI1(S25E), or DeSI1(S25E/C108S) were analyzed by denaturing Ni-NTA pulldown and mass spectrometry. A common analysis universe of 4,531 proteins was ranked by the S25E-versus-EV His-Ub t statistic; the 907 proteins in the lowest 20% define the DeSI1(S25E) reduced-ubiquitylation tail. Upper, complete ranked distribution. Middle, magnified lowest-20% tail; colored points denote proteins assigned for display to the indicated membrane categories, and selected proteins are labeled. Lower, non-exclusive category-membership rugs for outer membrane, ER membrane, ER exit site, coated-vesicle membrane, vesicle membrane, and Golgi membrane proteins. Numbers in parentheses indicate tail members/background members for each category. Statistical enrichment of these categories is shown in Supplementary Figure 3. **(D)** CDK4 activation separates protein-dominant and transcript-dominant anabolic programs. HCT-116 mAID–AMBRA1 cells were compared under CDK4-active conditions and following acute CDK4/6 inhibition with palbociclib. Matched RNA-seq and quantitative-proteomic datasets were analyzed by gene-set enrichment analysis. Each point represents a pathway plotted by its normalized enrichment score (NES) at the RNA level (x axis) and protein level (y axis), with positive values indicating enrichment under CDK4-active conditions. The dashed diagonal indicates equal RNA and protein responses. Orange denotes protein-dominant pathways and blue transcript-dominant pathways; labeled filled points indicate direct differential enrichment and open labeled points denote contextual or trending pathways. Oxidative-phosphorylation and mitochondrial programs preferentially shift at the protein level, whereas de novo pyrimidine biosynthesis is predominantly transcriptional.

Because these DeSI1(S25E)-enriched proteins spanned diverse biochemical functions but shared membrane- and organelle-targeting annotations, we asked whether their convergence reflected a common biophysical feature rather than a shared biological function. To test this independently of annotation, we calculated for each sequence the minimum value obtained across overlapping 19-residue windows using the Hessa biological hydrophobicity scale, with lower minimum-window ΔG_app values indicating greater membrane-insertion propensity^19^. DeSI1(S25E)-enriched interactors were shifted toward lower minimum-window ΔG_app values than wild-type-enriched interactors (p=0.00002; Figure 3B). This directional difference was also evaluated by permutation analysis matched for protein length and abundance. Consistent with the overall distributional shift, segments predicted to insert spontaneously into membranes (ΔG_app<0) were present in 9.2% of the S25E-enriched interactors versus none of the WT-enriched interactors. Thus, DeSI1(S25E)-enriched interactors exhibit a biophysical bias toward proteins containing more membrane-insertion-favorable hydrophobic sequences.

In a second screen, we performed Ni-NTA pulldowns of His-ubiquitin conjugates from HEK293F cells expressing either an empty vector (EV), DeSI1(S25E) or DeSI1(S25E;C108S). The pulldowns were carried out under denaturing conditions, thereby capturing covalent ubiquitylation while disrupting non-covalent interactions (Supplementary Table 2). We first ranked ubiquitylated proteins according to the S25E-versus-EV t statistic. Proteins in the lowest 20% of this ranked list, which showed the strongest reductions in His-ubiquitin recovery in DeSI1(S25E)-expressing cells, defined a reduced-ubiquitylation tail. Within this tail, 193 unique proteins mapped to membrane compartments, all of which were significantly enriched relative to the quantified background (Figure 3C and Supplementary Figure 3D). Catalytic reversal was then evaluated independently across the protein tail using the S25E/C108S-versus-S25E t statistic. The catalytic reversal analysis identified 165 unique proteins distributed across mitochondrial matrix and membrane, endoplasmic-reticulum, vesicle-membrane, and organelle-bounding-membrane modules (Supplementary Figure 3E). Consistent with this membrane-associated signature, proteins less ubiquitylated in DeSI1(S25E)-expressing cells were shifted toward lower minimum-window ΔG_app values relative to the control population and this shift persisted after matching the groups for protein length (p = 0.00015), indicating greater membrane-insertion propensity (Supplementary Figure 3F). The interactome and ubiquitylome showed limited protein-level overlap (Supplementary Tables 1,2) but converged on the same membrane compartments and hydrophobic, membrane-insertion-prone protein features, with the ubiquitylome extending the interactome signature to additional members of the same membrane systems.

Cyclin D-CDK4/6 activity drives a transcriptional program necessary for the G1/S transition (see Introduction). Having defined a post-translational DeSI1 pathway downstream of cyclin D-CDK4/6, we asked whether the biophysical signature associated with this pathway was also evident in the broader cellular response to CDK4/6 activity. Specifically, we integrated matched transcriptomic and proteomic profiles from cells in which endogenous cyclin D-CDK4/6 activity was acutely increased by AMBRA1 depletion or inhibited with palbociclib in AMBRA1-depleted cells (Supplementary Figure 3A), similar to the experiments shown in Figure 1A-C. This analysis revealed distinct transcriptional and protein-level responses (Figure 3D). Certain signatures (*e.g.,* de novo pyrimidine metabolism and DNA replication) were induced predominantly at the transcriptional level, as expected. In contrast, other signatures (*e.g.,* oxidative phosphorylation and one-carbon metabolism) showed a strong protein-level response. Thus, acute CDK4/6 activation elicits separable transcriptional and protein-dominant responses, suggesting a coordination between biosynthetic resource supply and the abundance of membrane and mitochondrial proteins. Notably, proteins showing the strongest protein-dominant response to CDK4/6 activation were enriched for lower ΔG_app values relative to the remaining proteome, and this difference persisted after matching for protein length (Supplementary Figure 3G). These findings indicate that the membrane-insertion-prone protein class identified in the DeSI1 screens is also preferentially represented among the protein-level outputs of CDK4/6 activation.

### Monomerization exposes a hydrophobic substrate-binding surface

As an initial validation of the screening results, we expressed wild-type DeSI1 or DeSI1(S25E) in HEK293T cells, with or without co-expression of cyclin D1-CDK4. Following DeSI1 immunoprecipitation, we assessed its association with endogenous proteins by immunoblotting. Wild-type DeSI1 associated with VCP, ubiquilins 1 and 2, BAG6, and polyubiquitylated proteins (Figure 4A and Supplementary Figure 4A). The phospho-deficient DeSI1(S25A) mutant displayed the same wild-type-like interaction profile (Supplementary Figure 4A). In contrast, DeSI1(S25E), as well as wild-type DeSI1 phosphorylated upon cyclin D1-CDK4 co-expression, lost these interactions and instead associated with integral membrane proteins, such as COX4, TIM23, SLC25A5, and ATP1A1. Treatment with palbociclib largely restored the interaction profile of unphosphorylated DeSI1 (Figure 4A), demonstrating that CDK4 activity and the downstream phosphorylation of S25 in DeSi1 functions as a switch controlling this transition.

**Figure 4.**
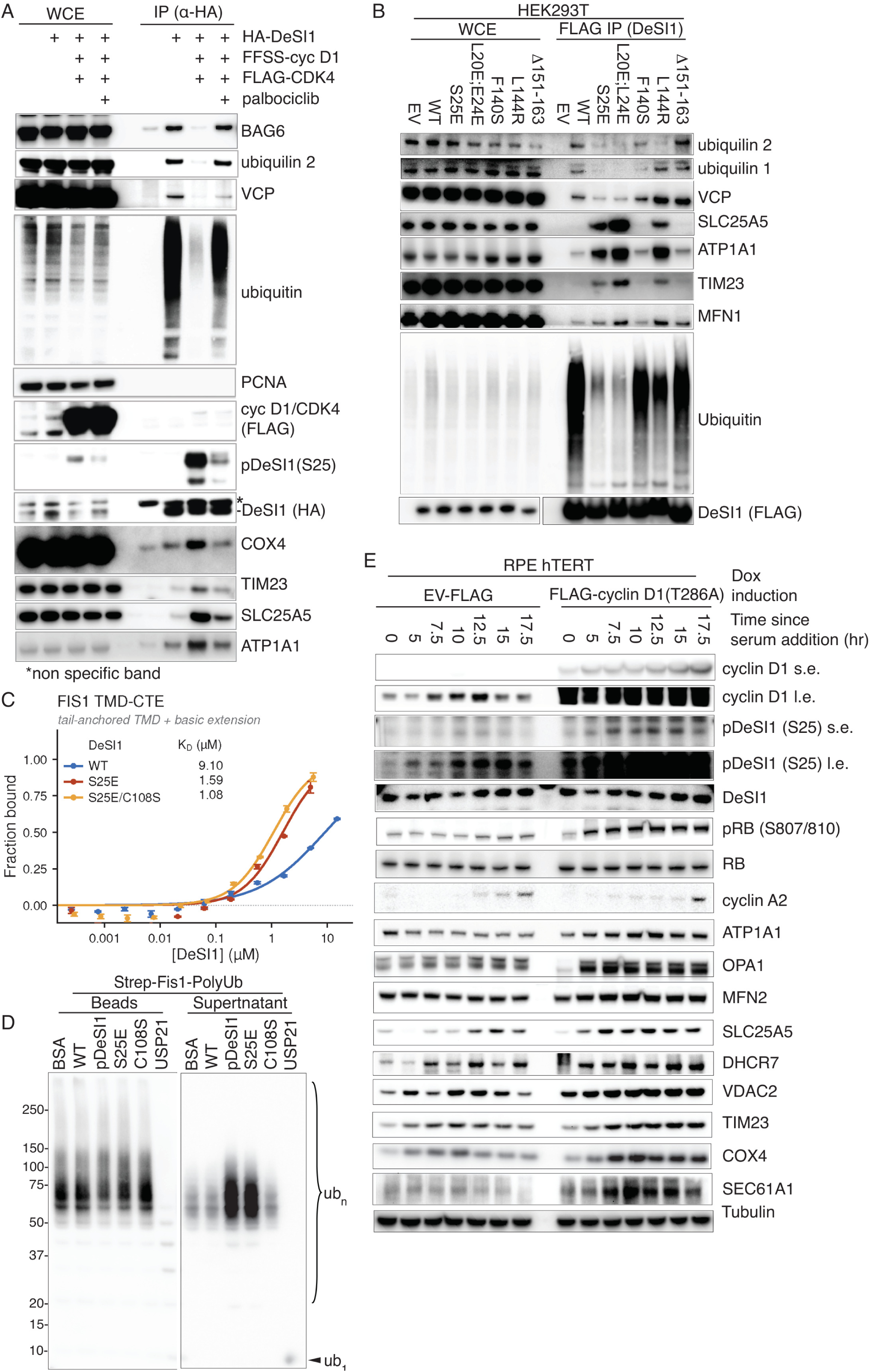
DeSI1 monomerization exposes a hydrophobic substrate-binding surface and promotes deubiquitylation of hydrophobic clients. **(A)** CDK4-dependent reprogramming of the DeSI1 interactome. HEK293T cells were transfected with pcDNA3-HA-DeSI1 alone or together with FFSS-cyclin D1 and Flag-CDK4, and treated with 0.5 μM palbociclib where indicated. Whole-cell extracts (WCE) and anti-HA immunoprecipitates were immunoblotted for the indicated proteins. pDeSI1 (S25) reports CDK4-dependent phosphorylation; PCNA serves as a negative control for non-specific recovery. Asterisk, non-specific band. **(B)** Interface mutants uncouple oligomeric state from phosphorylation. HEK293T cells expressing Flag-tagged EV, WT, S25E, L20E;L24E, F140S, L144R or Δ151–163 DeSI1 were subjected to anti-Flag immunoprecipitation. WCE and immunoprecipitates were immunoblotted for the indicated proteins. Polyubiquitylated species are detected as a high-molecular-weight smear. **(C)** Direct binding of recombinant DeSI1 to the FIS1 tail-anchored transmembrane domain with its basic C-terminal extension (FIS1 TMD-CTE), measured by fluorescence polarization. Fluorescently labelled FIS1 TMD-CTE peptide was titrated with recombinant DeSI1 WT, S25E or S25E/C108S. Points, mean ± s.d. of three biological replicates; curves, Hill fits; fitted Kᴅ values are inset. Peptide sequences, physicochemical parameters and 95% confidence intervals are given in Supplementary Table 3. **(D)** Deubiquitylation of FIS1-peptide-anchored polyubiquitin chains. Polyubiquitin chains were assembled on streptavidin-immobilized FIS1 TMD-CTE peptide and incubated with BSA, WT DeSI1, phosphorylated DeSI1 (pDeSI1), S25E, C108S or USP21. Bead-bound and supernatant fractions were resolved by SDS-PAGE and immunoblotted with anti-ubiquitin. **(E)** Sustained CDK4 activity drives substrate accumulation during G1. RPE-hTERT cells carrying doxycycline-inducible EV-Flag or Flag-cyclin D1(T286A) were serum-starved, induced with doxycycline, and released into G1 by serum addition for the indicated times. Whole-cell extracts were immunoblotted for the indicated proteins. Cyclin D1, cyclin A2 and RB phosphorylation (S807/810) monitor G1 progression; pDeSI1 (S25) reports activation of the switch; tubulin serves as loading control. s.e., short exposure; l.e., long exposure

We next tested the interface mutants characterized in Figure 2. The obligate monomeric DeSI1(L20E;L24E) closely phenocopied DeSI1(S25E), showing loss of binding to VCP, ubiquilins 1 and 2, and polyubiquitin chains, accompanied by increased association with TIM23, SLC25A5, ATP1A1, and MFN1 (Figure 4B). DeSI1(L144R) similarly gained interactions with hydrophobic proteins, although its loss of association with quality-control factors and polyubiquitin chains was less pronounced. In contrast, the dimeric DeSI1(Δ151-163) and the monomer-dimer equilibrium mutant DeSI1(F140S) largely retained a wild-type-like interaction profile (Figure 4B). These results indicate that the DeSI1 interactome switch is governed by its oligomeric state.

Inspection of the experimentally determined human DeSI1 structure revealed that the hydrophobic dimer-interface residues L20, L24, F140, and L144 are positioned adjacent to the acidic residues E34, E134, and the regulatory S25 site (Supplementary Figure 4B). Thus, monomerization exposes a surface with both hydrophobic and acidic features. Because S25 is unphosphorylated in the experimental structure, the additional negative charge introduced by S25 phosphorylation is not captured. This composite surface, combining acidic and hydrophobic features, could favor recognition of membrane-targeting sequences containing basic and hydrophobic residues.

To test this possibility, we selected two representative targeting sequences with distinct architectures: the FIS1 tail-anchor, comprising a hydrophobic transmembrane domain and basic C-terminal extension, and the amphipathic, basic N-terminal mitochondrial targeting sequence of COX4. We then measured their direct binding to DeSI1 by fluorescence polarization. Binding to the FIS1 tail-anchored TMD and C-terminal extension correlated with DeSI1 oligomeric state: wild-type DeSI1 bound with modest affinity (Kᴅ = 9.10 µM), whereas DeSI1(S25E) and DeSI1(S25E;C108S) bound substantially more tightly (1.59 and 1.08 µM, respectively) (Figure 4C and Supplementary Table 3). The COX4 presequence showed a similar pattern, with Kᴅ values of 7.45 µM (95% CI 5.74–9.67) for wild-type DeSI1, compared with 1.96 µM (0.81–4.73) and 0.88 µM (0.83–0.95) for DeSI1(S25E) and DeSI1(S25E;C108S), respectively (Supplementary Figure 4C, left, and Supplementary Table 3). These results suggest that both tail-anchored TMDs and N-terminal MTSs engage the same monomeric surface. This interaction depended on the hydrophobic element: a matched-length COX4 peptide in which the hydrophobic face was replaced by serine and asparagine (GRAVY −1.200 versus −0.036 for the cognate presequence) showed no saturable binding to any DeSI1 variant (Supplementary Figure 4C, middle, and Supplementary Table 3). Binding also exhibited the predicted electrostatic contribution, as increasing ionic strength progressively weakened DeSI1(S25E;C108S) binding to the COX4 presequence (Kᴅ = 0.88, 1.87, and 6.46 µM at 150, 225, and 300 mM NaCl, respectively; Supplementary Figure 4C, right).

Together, these results suggest that monomerization increases DeSI1 affinity for hydrophobic targeting elements such as TMDs and MTSs through a composite surface involving both hydrophobic and electrostatic interactions.

To test whether binding and deubiquitylase activity are coupled, we assembled polyubiquitin chains on streptavidin-immobilized FIS1 and COX4 peptides and monitored chain release. On the FIS1 substrate, both DeSI1(S25E) and phosphorylated DeSI1 released ubiquitylated chains into the supernatant, with a corresponding decrease in bead-associated material, whereas BSA, wild-type DeSI1, and DeSI1(C108S) produced only background release (Figure 4D). The same activity profile was observed with COX4-anchored polyubiquitin chains (Supplementary Figure 4D), and time-course experiments showed progressive release from both FIS1- and COX4-based substrates (Supplementary Figure 4E). In contrast, a hydrophilic COX4-derived peptide supported polyubiquitin assembly but showed no DeSI1-dependent release (Supplementary Figure 4F), demonstrating that cleavage requires a hydrophobic targeting element rather than the ubiquitin chain alone. Thus, monomeric DeSI1 is not a generic K48 deubiquitylase but preferentially acts on hydrophobic substrates bearing TMD or MTS elements.

Finally, we asked whether sustained cyclin D1-CDK4/6 signaling affects substrate candidates of DeSI1 in cells. RPE-hTERT cells were synchronized in G0 by 3 days of serum starvation and then released with or without doxycycline-induced expression of the stable cyclin D1(T286A) mutant^20^. In cells re-entering the cell cycle in the absence of doxycycline, DeSI1 is phosphorylated in G1 in parallel with the induction of endogenous cyclin D1, and this phosphorylation decreases at the G1/S transition (Figure 4E). By contrast, doxycycline-mediated cyclin D1(T286A) expression sustained DeSI1 S25 phosphorylation throughout G1 and into G1/S, accompanied by increased abundance of proteins bearing hydrophobic or amphipathic targeting elements, including COX4, TIM23, SLC25A5, and ATP1A1, MFN2, OPA1, DHCR7, VDAC2, and SEC61A1, without altering total DeSI1 levels (Figure 4E). To directly test the requirement for DeSI1 phosphorylation, we depleted endogenous DeSI1 and reconstituted RPE-hTERT cells with doxycycline-inducible DeSI1(S25A) or DeSI1(S25E). Compared with DeSI1(S25A), DeSI1(S25E) expression led to accumulation of hydrophobic proteins during serum release, despite comparable cell-cycle progression, as assessed by levels of cyclin E1 and cyclin A2 and RB phosphorylation (Supplementary Figure 4G). Thus, the DeSI1 dimer-to-monomer switch appears to operate within the physiological window of cyclin D1-CDK4/6 activity to regulate the abundance of its putative substrates.

### Phosphorylated DeSI1 stabilizes nascent hydrophobic proteins

Figures 3 and 4 indicated that phosphorylated DeSI1 preferentially engages hydrophobic proteins and can remove ubiquitin chains from hydrophobic peptides. Because membrane-destined hydrophobic proteins pose intrinsic challenges to biogenesis and are subject to ubiquitin-dependent quality control from the onset of synthesis, we asked whether DeSI1 activity extends the lifetime of nascent hydrophobic proteins in cells.

We used fluorescent Global Protein Stability (GPS) reporters^21^, in which EGFP fused to a TMD-containing sequence is expressed from a bicistronic transcript with an internal mCherry control, so that the GFP/RFP ratio reports degradation independently of expression (Supplementary Figure 5A). In U2OS cells, reporters containing the hydrophobic segments of SLC25A5, HMGCR, MFN2, and DHCR7 were stabilized both upon induction of cyclin D1(T286A) and upon reconstitution of *DeSI1*-knockout cells with DeSI1(S25E), whereas wild-type DeSI1 had no effect (Figure 5A, top and bottom panels). In contrast, a soluble peptide derived from CA2 and unfused GFP were unaffected. Consistent with this selectivity, cyclin D1(T286A) and DeSI1(S25E) did not stabilize hydrophilic peptides derived from SLC25A5 or DHCR7 (Figure 5B). Stabilization required ongoing protein synthesis, as it was abolished by inhibiting translation with cycloheximide (Figure 5A, middle panel, and Supplementary Figure 5B, bottom panel). Moreover, the stabilizing effect was lost upon mitochondrial or ER stress and during serum starvation, but was restored upon stress removal or serum re-addition, respectively (Supplementary Figure 5B,C). Thus, DeSI1-mediated stabilization is dictated by the hydrophobic determinant itself and appears to act specifically on newly synthesized proteins, when their hydrophobic features are exposed during translation.

**Figure 5.**
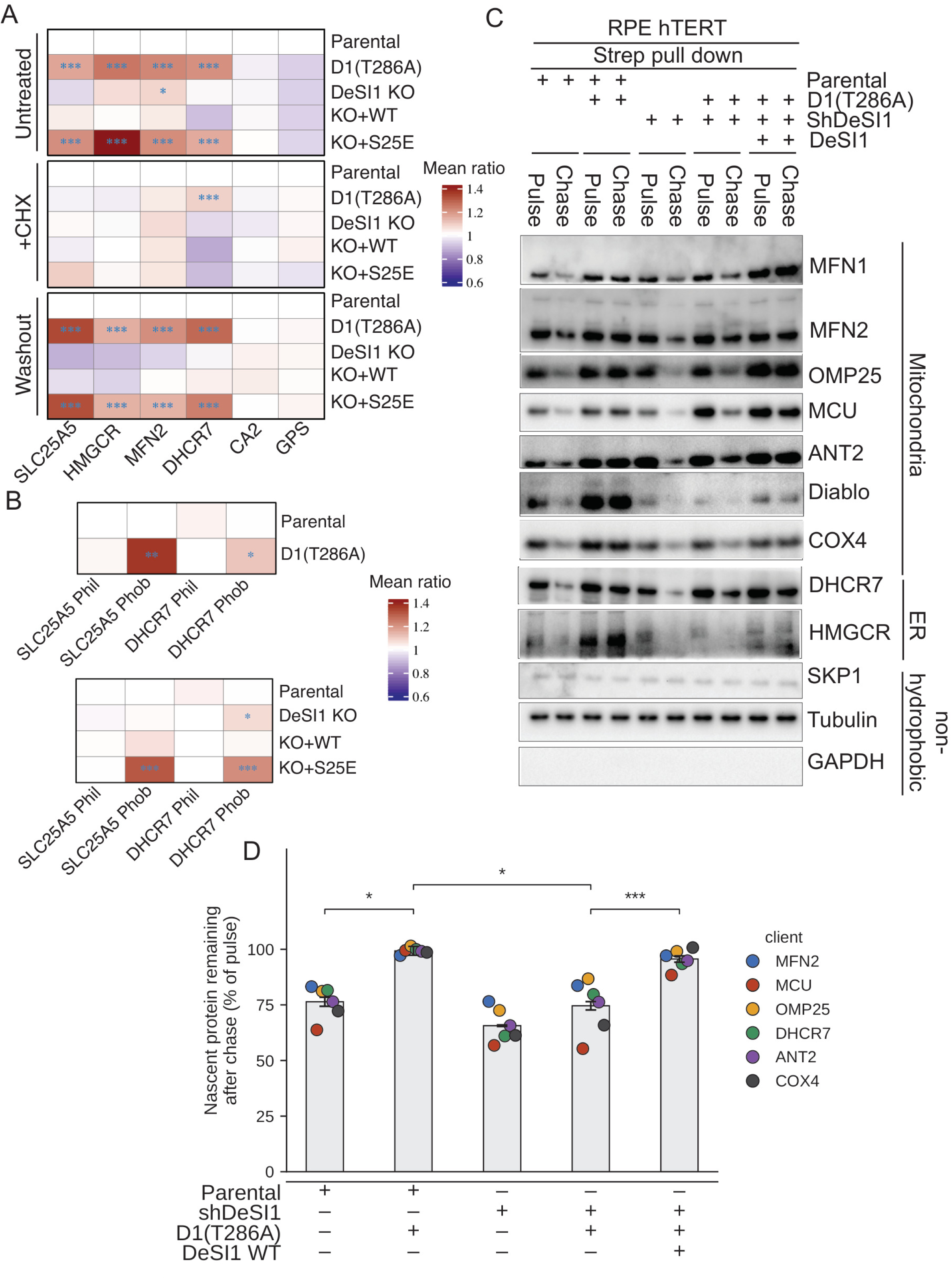
Cyclin D1-CDK4 phosphorylation of DeSI1 protects nascent hydrophobic proteins from degradation. **(A)** GPS reporter stability requires ongoing translation. U2OS cells stably expressing GFP/RFP reporters (SLC25A5, HMGCR, MFN2, DHCR7, and the soluble controls CA2 and unfused GFP) were analysed under basal conditions, after cycloheximide (CHX, 4 h), or after CHX washout (120 min). Cell lines: parental; parental expressing doxycycline-inducible cyclin D1 T286A (+Dox); DeSI1 KO; and DeSI1 KO reconstituted with pBabe HA-DeSI1 WT or S25E. Heatmaps show mean GFP/RFP ratios normalized to parental cells within each condition (parental = 1). Asterisks indicate significantly increased stability relative to parental within the same condition; pooled-residual two-sided t-tests with Holm correction (*P < 0.05, P < 0.01,* P < 0.001); n = 3 independent experiments. **(B)** Stabilization is specific to the hydrophobic degron. U2OS cells expressing paired reporters in which the hydrophobic (Phob) or hydrophilic (Phil) segment of SLC25A5 or DHCR7 is fused to GFP with an RFP internal control. Upper heatmap, parental versus cyclin D1 T286A; lower heatmap, parental, DeSI1 KO, and KO reconstituted with WT or S25E DeSI1. GFP/RFP ratios normalized to parental; statistics as in (A); n = 3 independent experiments. **(C)** CDK4-driven protection of nascent membrane proteins requires DeSI1. RPE-hTERT cells of five genotypes-parental; shDeSI1; parental + cyclin D1 T286A; shDeSI1 + cyclin D1 T286A; and shDeSI1 + cyclin D1 T286A + HA-DeSI1 WT-were released into G1, pulse-labelled with AHA and chased for 90 min. Biotinylated nascent proteins were captured on streptavidin resin. Streptavidin pull-downs were immunoblotted for ER clients (DHCR7, HMGCR), mitochondrial clients (MFN2, MFN1, OMP25, MCU, ANT2, DIABLO, COX4) and non-hydrophobic controls (SKP1, tubulin, GAPDH). Whole-cell AHA-labelling, capture-efficiency and pathway controls are shown in Supplementary Figure 5. cyclin D1, total DeSI1 and pDeSI1 (S25) report the state of the pathway; vinculin, loading control. Only re-expression of wild-type DeSI1 in the shDeSI1 + cyclin D1 T286A background recapitulates the protection seen in parental cells expressing cyclin D1 T286A. S.E., short exposure; L.E., long exposure. **(D)** Quantification of the AHA pulse-chase in (C). Band intensities for six nascent clients (MFN2, MCU, OMP25, DHCR7, ANT2, COX4) were measured in the streptavidin-captured fraction; retention is expressed as the chase signal as a percentage of the pulse signal within the same cell line (chase/pulse × 100), without normalization to a ceiling, so values near or slightly above 100% indicate no detectable loss during the chase. Bars show the mean of three independent experiments; error bars are SEM (n = 3 experiments). Each colored point is one client, averaged across the three experiments, and is shown to illustrate consistency of the effect across clients; clients were not treated as independent replicates. Condition composition is indicated by +/– below each bar. Conditions were compared by paired two-tailed t-test on experiment-level means (n = 3): Parental versus D1(T286A), p = 0.012; D1(T286A) versus shDeSI1 + D1(T286A), p = 0.011; shDeSI1 + D1(T286A) versus shDeSI1 + D1(T286A) + DeSI1 WT, p = 0.0006. *p < 0.05, ***p < 0.001.

To test whether DeSI1 acts on endogenous nascent proteins, we metabolically labelled G1-synchronized RPE-hTERT cells with the methionine analogue azidohomoalanine (AHA).Newly synthesized proteins were then biotinylated by click chemistry, enriched by streptavidin pull-down, and analyzed by immunoblotting for individual client proteins. Upon AHA chase, the levels of AHA-labelled ER proteins (DHCR7 and HMGCR) and mitochondrial proteins (MFN1, MFN2, OMP25, MCU, ANT2, COX4, and DIABLO) decreased, as expected (Figure 5C and Supplementary Figure 5D). Expression of cyclin D1(T286A) increased the fraction of these nascent proteins that remained after the chase, whereas the levels of the non-hydrophobic control proteins SKP1, tubulin, and GAPDH were unaffected (Figure 5C). This protective effect was lost upon DeSI1 depletion, indicating that DeSI1 acts downstream of cyclin D1-CDK4. Re-expression of DeSI1 in DeSI1-depleted cells restored the ability of cyclin D1(T286A) to protect the nascent protein pool (Figure 5C,D). Expression of HPV E7, which sequesters RB and consequently activates E2F, did not induce DeSI1 S25 phosphorylation and did not reproduce the effect of cyclin D1(T286A) on hydrophobic proteins (Supplementary Figure 5E). Thus, protection of nascent hydrophobic proteins is not a downstream consequence of E2F activation but instead reflects a distinct cyclin D-CDK4/6 output mediated through DeSI1.

### DeSI1 affects mitochondrial respiration

Proteins stabilized by phosphorylated DeSI1 converge functionally on membrane and mitochondrial functions. Thus, we asked whether DeSI1’s post-translational program regulates mitochondrial function. Induction of DeSI1(S25E) increased basal mitochondrial OCR, ATP-linked respiration and proton leak, while wild-type DeSI1 produced more modest increases in respiratory capacity (Figure 6A,B). Maximal respiration was also strongly increased by DeSI1(S25E), whereas coupling efficiency remained unchanged (Supplementary Figure 6A). Transmission electron microscopy revealed nominal differences in several mitochondrial morphometric parameters, including matrix density and lucency, cristae area fraction and aspect ratio (Supplementary Figure 6B,C). A similar increase in mitochondrial respiration was observed upon induction of cyclin D1(T286A) expression, but only in DeSI1 expressing cells (Supplementary Figure 6D). Together, these data show that acute DeSI1 activation increases mitochondrial respiratory capacity.

**Figure 6.**
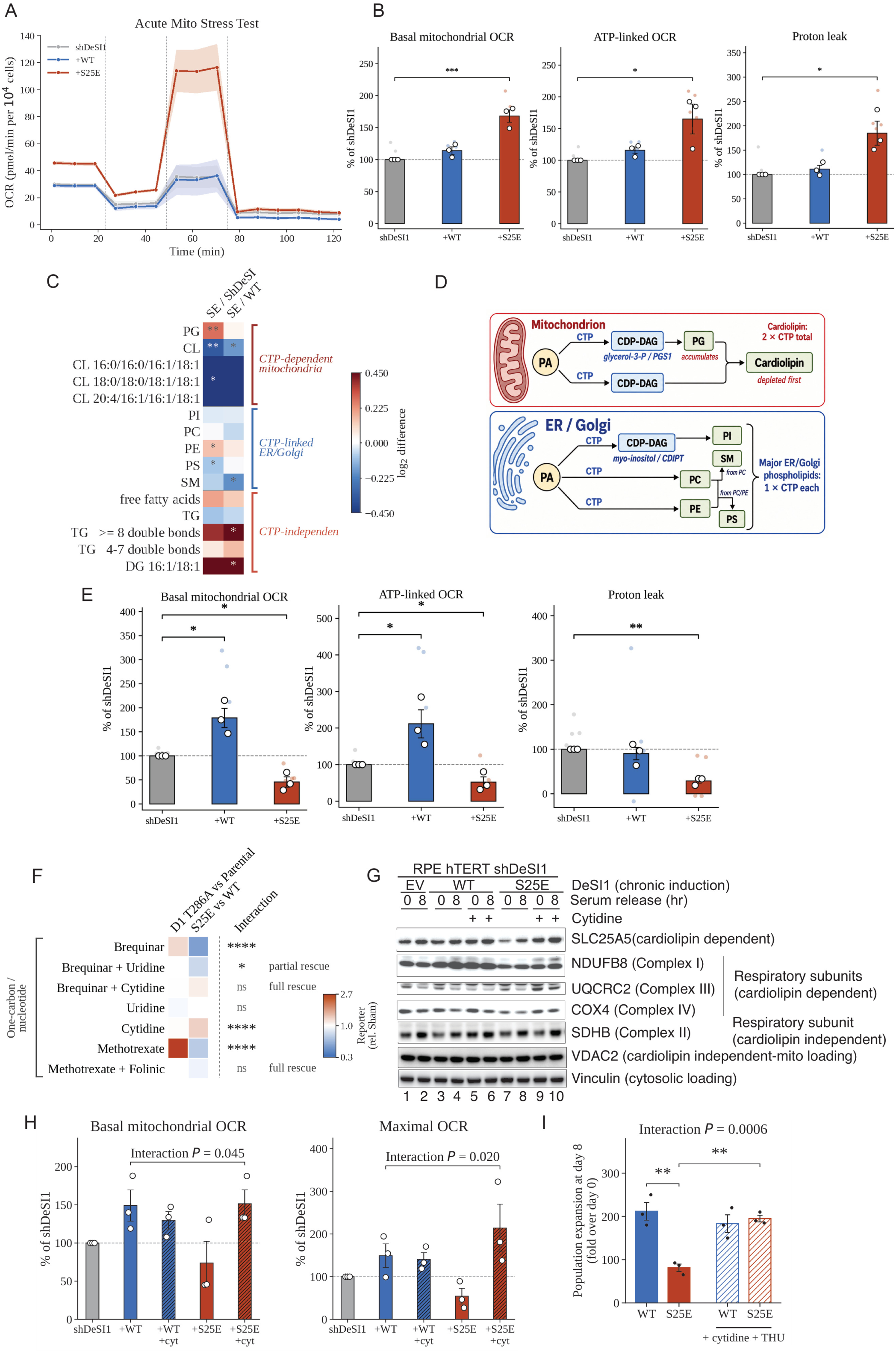
Phospho-DeSI1 acutely expands mitochondrial capacity, whereas sustained activation creates a cytidine-nucleotide constraint. **(A)** Seahorse Mito Stress Test after acute (24 h) doxycycline induction of shDeSI1 RPE-hTERT cells reconstituted with inducible DeSI1 WT or S25E. OCR was normalized in each well to post-assay cell number and expressed per 10,000 cells; technical wells were averaged within each independent experiment. Lines show mean ± SEM across n = 3 independent biological experiments; shading indicates SEM. Dashed lines indicate sequential injection of oligomycin, FCCP and rotenone+antimycin A. **(B)** Basal mitochondrial OCR, ATP-linked OCR and proton leak derived from the Seahorse Mito Stress Test. OCR was normalized in each well to post-assay cell number and endpoint values were expressed relative to the matched shDeSI1 condition within each experiment. Bars show mean ± SEM of n = 3 independent biological experiments; small shaded points indicate technical wells and open circles indicate biological-replicate means. Statistical analysis was performed on the underlying cell-normalized biological-replicate means using one-way repeated-measures/randomized-block ANOVA on log₂-transformed values followed by two-sided Dunnett comparisons versus shDeSI1. *P < 0.05, **P < 0.01, ***P < 0.001; stars denote Dunnett-adjusted P values. **(C)** Acute S25E produces phosphatidylglycerol accumulation and cardiolipin loss without generalized lipid depletion. RPE-hTERT cells depleted of endogenous DeSI1 (shDeSI1) or reconstituted with doxycycline-inducible wild-type DeSI1 (WT) or DeSI1 S25E (SE) were analyzed after 72 h serum starvation and after 8 h serum-stimulated G1 entry (n = 6 matched biological replicates per genotype). The heatmap shows the difference in the G1-versus-starved response between S25E and the indicated reference line (SE/ShDeSI and SE/WT). Lipid intensities were median-normalized within each sample. For lipid-class rows, normalized linear abundances of all detected species within each class were summed per sample and log₂-transformed; the G1−Starved response was then calculated within each matched biological replicate, and heatmap values represent the mean difference between genotypes. Individual cardiolipin species and TG subclasses were analyzed analogously. Rows are grouped as CTP-dependent mitochondrial lipids (PG and CL), CTP-linked ER/Golgi lipids (PI, PC, PE, PS and SM; PS and SM are indirectly linked through CTP-dependent precursor lipids), or CTP-independent lipid pools. Asterisks denote nominal two-sided one-sample t-tests of the matched difference-in-differences against zero (*P < 0.05, **P < 0.01, ***P < 0.001). For the nine predefined lipid-class endpoints, P values were additionally corrected within each contrast using the Benjamini–Hochberg procedure. In the SE/ShDeSI comparison, PG (q = 0.013) and CL (q = 0.024) remained significant after correction; PE was borderline (q = 0.050) and PS did not remain significant (q = 0.081). In the SE/WT comparison, the nominal decreases in CL and SM did not survive correction (both q = 0.137). **(D)** Schematic of CTP utilization in phospholipid synthesis. PI, PC and PE each require one CTP-dependent activation step. In mitochondria, one CTP-derived CDP-DAG equivalent is used to generate PG and a second CDP-DAG equivalent is required together with PG for cardiolipin synthesis. Thus, cardiolipin requires two CTP-derived activated lipid equivalents, providing a potential explanation for PG accumulation and preferential cardiolipin depletion under limiting cytidine-nucleotide supply. **(E)** Chronic phospho-DeSI1 reverses the acute respiratory gain. Basal mitochondrial OCR, ATP-linked OCR, proton leak and coupling efficiency were derived from cell-count-normalized Seahorse XF Mito Stress Tests after 7 days of induction of DeSI1 WT or S25E in shDeSI1 RPE-hTERT cells. OCR was normalized in each well to the post-assay cell count. For visualization, basal mitochondrial OCR, ATP-linked OCR and proton leak were expressed relative to the matched shDeSI1 condition within each biological experiment (shDeSI1 = 100%). Bars show mean ± SEM of n = 3 independent biological experiments. Small shaded points represent technical replicates and large open circles represent biological-replicate means. Statistical analysis was performed on the underlying cell-normalized biological-replicate means using a one-way ANOVA. Brackets denote two-sided comparisons versus shDeSI1; *P < 0.05, **P < 0.01. **(F)** Cytidine/THU and folinic acid rescue the one-carbon/nucleotide phenotypes. Focused view of reporter signal for the one-carbon and nucleotide arm of the screen, plotted on the color scale of Supplementary Figure 7; values are normalized within each setup to Sham (= 1.0), n = 3. Asterisks report the genotype × compound interaction, that is, whether a compound shifts competition differently in D1(T286A) vs. parental than in DeSI1(S25E) vs. wild-type DeSI1 expressing cells. Contrasts were tested against the residual variance pooled across the full screen (two-way ANOVA) and Holm–Šidák-adjusted across the seven compounds shown (*q < 0.05, ****q < 0.0001; ns, not significant). **(G)** Cytidine restores SLC25A5 and respiratory-chain subunits. RPE-hTERT shDeSI1 cells carrying doxycycline-inducible empty vector (EV), DeSI1 WT or DeSI1(S25E) were induced for 7 days (chronic), with or without cytidine throughout the induction window as indicated, then serum-released for 0 or 8 h. Whole-cell extracts were immunoblotted for the cardiolipin-sensitive carrier SLC25A5 and the cardiolipin-sensitive respiratory-chain subunits NDUFB8 (Complex I), UQCRC2 (Complex III), COX4 (Complex IV), together with the cardiolipin-independent respiratory subunit SDHB (Complex II). Vinculin, cytosolic loading control; VDAC2, cardiolipin-independent mitochondrial loading control. **(H)** Cytidine restores mitochondrial respiratory capacity in chronic S25E cells. Basal mitochondrial and maximal OCR were derived from cell-count-normalized Seahorse XF Mito Stress Tests after 7 days of induction of WT or S25E DeSI1 ± cytidine, with cytidine present throughout the induction period where indicated. Oxygen-consumption rate (OCR) was measured by Seahorse XF Mito Stress Test and normalized in each well to the corresponding post-assay cell count. Technical wells were averaged within each independent experiment before aggregation. The trace shows mean ± SEM across n = 3 independent biological experiments, with sequential injection of oligomycin, FCCP and rotenone+antimycin A. Basal mitochondrial OCR was calculated as the final pre-oligomycin OCR minus the minimum OCR following rotenone+antimycin A, and maximal OCR as the maximum FCCP-stimulated OCR minus the minimum OCR following rotenone+antimycin A. For visualization, endpoint values were expressed relative to the matched shDeSI1 condition from the same experiment (shDeSI1 = 100%). Bars show mean ± SEM and circles represent the three biological-replicate means. Statistical analysis of WT and S25E cells ± cytidine was performed by two-factor repeated-measures ANOVA, with genotype and cytidine as within-experiment factors and independent experiment as the matched factor. OCR parameters were analyzed on log2-transformed cell-count-normalized biological-replicate means. Interaction P values shown are the nominal genotype × cytidine interaction. **(I)** Cytidine rescues the chronic S25E growth defect when cytidine deamination is inhibited. RPE-hTERT shDeSI1 cells expressing doxycycline-inducible DeSI1 wild-type or DeSI1(S25E) were cultured for 8 days with or without cytidine (100 µM) together with tetrahydrouridine (THU; 100 µM), which inhibits cytidine deamination. Population expansion was calculated for each biological replicate as the day-8 cell count divided by the day-0 cell count and expressed as fold over day 0. Bars show mean ± SEM of n = 3 independent biological experiments, with individual replicates overlaid. Statistical analyses were performed on log2-transformed fold-expansion values. Indicated pairwise comparisons were assessed by two-sided Welch’s t-tests (**P < 0.01); the genotype × treatment interaction was assessed by two-way ANOVA (P = 0.0006).

Given the mitochondrial phenotype caused by DeSI1(S25E), we asked which metabolic resources are required to support this gain in mitochondrial function. We profiled polar metabolites and lipids in DeSI1-depleted cells reconstituted with wild-type DeSI1 or DeSI1(S25E) during G1 (see experimental design in Supplementary Figure 6E). The most prominent phenotype involved the mitochondrial arm of phospholipid biosynthesis which uses cytidine diphosphate-diacylglycerol (CDP-DAG) as an intermediate to generate cardiolipin, the signature phospholipid of the inner mitochondrial membrane (Figure 6C,D). In this pathway (see Figure 6D), phosphatidic acid (PA) is first activated with CTP to form diphosphate-diacylglycerol (CDP-DAG). One CDP-DAG is then used with glycerol-3-phosphate to generate phosphatidylglycerol (PG), which is subsequently combined with a second CDP-DAG molecule to produce cardiolipin. Relative to shDeSI1 cells, DeSI1(S25E)-expressing cells showed significant PG accumulation and cardiolipin depletion after multiple-testing correction, consistent with a constraint in the mitochondrial phospholipid pathway. By contrast, the major CTP-linked ER/Golgi lipids phosphatidylinositol (PI), phosphatidylcholine (PC), and phosphatidylethanolamine (PE), phosphatidylserine (PS), and sphingomyelin (SM) showed only minor changes (Figure 6C,D). CTP-independent lipid pools, including free fatty acids and total triacylglycerol, were likewise not reduced. Thus, the phenotype was not a generalized impairment of lipid synthesis but was preferentially associated with the mitochondrial phospholipid pathway.

Polar metabolomics provided a complementary view of pathway demand. Glycerol-3-phosphate and myo-inositol (the co-substrates used with CDP-DAG to generate PG and PI, respectively) were reduced, with glycerol-3-phosphate showing the strongest change, while purine nucleotides and the uridine pool were largely preserved (Supplementary Figure 6F). Notably, cardiolipin synthesis requires two CTP-derived CDP-DAG equivalents: one to generate PG and a second for the cardiolipin synthesis reaction itself (Figure 6D)^22^. Thus, cardiolipin biosynthesis places a greater demand on CTP-dependent CDP-DAG production than the corresponding ER/Golgi phospholipid pathways, which require only a single CDP-DAG equivalent. This stoichiometry provides a potential explanation for the selective lipid phenotype. Consistent with a biosynthetic constraint, less-unsaturated cardiolipin species were preferentially depleted, whereas more polyunsaturated species were relatively spared (Figure 6C). Together, these findings suggest that the DeSI1-dependent increase in membrane-targeted protein supply places a disproportionate demand on cytidine-nucleotide-dependent phospholipid synthesis, with cardiolipin production showing particular sensitivity to this metabolic constraint.

The experiments shown in Figures 3-5 and 6A-C were performed 24 hours after transient expression or DOX induction of either DeSI1(S25E) or cyclin D(T286A). Thus, acute DeSI1 activation enhanced mitochondrial respiration despite a concurrent lipid defect that could compromise mitochondrial function over time. We therefore asked whether this initially beneficial mitochondrial phenotype could be sustained during prolonged DeSI1 activation. We found that after 7 days of induction, the acute respiratory gain inverted: re-expression of wild-type DeSI1 increased basal mitochondrial OCR and ATP-linked respiration relative to shDeSI1 cells, consistent with a role for physiologically regulated DeSI1 in supporting mitochondrial respiratory capacity (Figure 6E). In contrast, chronic DeSI1(S25E) expression reduced basal OCR, ATP-linked respiration, and proton leak below shDeSI1 levels, revealing respiratory decompensation when the DeSI1 switch is constitutively engaged. Similarly, levels of the inner-membrane carrier SLC25A5 and subunits of the cardiolipin-sensitive respiratory complexes I, III and IV (e.g., NDUFB8, UQCRC2, and COX4) were reduced in chronically DeSI1(S25E) expressing cells (Figure 6G, compare lanes 7-8 to 1-4). By contrast, the cardiolipin-independent proteins VDAC2 and SDHB were unchanged. Finally, the G1 population of cells chronically expressing DeSI1(S25E) increased with a respective reduction of the population in S phase, a phenotype not observed after acute induction of DeSI1(S25E) or chronic expression of wild-type DeSI1 (Supplementary Figure 6G).

Thus, phosphorylation of DeSI1 initially increases mitochondrial capacity, whereas constitutive activation seems to progressively uncouple this response from the resources required to sustain it.

### Constitutive DeSI1 activation causes cerebellar degeneration and a cytidine-nucleotide supply-demand imbalance

Despite enhanced respiratory capacity upon acute activation of DeSI1, the cardiolipin deficit and altered cytidine-related metabolites, suggested a cytidine-nucleotide supply-demand imbalance when the DeSI1 switch is persistently locked in the active state. To investigate this mechanistically, we performed a competitive chemical-genetic screen^23^ in which DeSI1 knockout cells stably expressing either DeSI1(S25E) or wild-type DeSI1 were seeded together and challenged with 26 metabolic and signaling perturbations using relative competitive fitness as the readout (assay design shown in Supplementary Figure 7A). In parallel, we carried out the same assays with parental cells stably expressing cyclin D1(T286A), which engages both the RB-E2F transcriptional arm and the DeSI1 post-translational arm of cyclin D1 signaling. Most perturbations produced concordant responses in the two systems. In contrast, Brequinar and methotrexate produced the strongest discordant responses: cyclin D1(T286A) cells were relatively resistant, whereas DeSI1(S25E) cells were sensitive (Figure 6F and Supplementary Figure 7B). Brequinar inhibits dihydroorotate dehydrogenase (DHODH), a key enzyme in de novo pyrimidine synthesis^24^. Notably, the sensitivity of DeSI1(S25E) cells to brequinar was rescued by exogenous cytidine, and to a lesser extent by uridine, consistent with a pyrimidine supply limitation in these cells. Moreover, cytidine administered together with THU selectively increased the fitness of DeSI1(S25E) cells, whereas uridine had no detectable effect. Methotrexate perturbs folate-dependent nucleotide metabolism, and its effect is also consistent with the idea that DeSI1(S25E) cells are vulnerable to impaired nucleotide supply. Folinic acid rescued methotrexate sensitivity, supporting an on-target mechanism. Thus, the discordant responses were concentrated in one-carbon/nucleotide metabolism, which supplies pyrimidine precursors for nucleotide biosynthesis and folate-dependent one-carbon units. Consistent with the relevance of this axis, de novo pyrimidine biosynthesis was transcriptionally induced by CDK4/6 activity (Figure 3D).

Together, these results support a model in which constitutive DeSI1 activation increases membrane and mitochondrial protein demand that can become limiting when nucleotide and lipid biosynthetic capacity is insufficient. By contrast, cyclin D1(T286A)-expressing cells are comparatively protected, consistent with the broader biosynthetic response elicited by CDK4/6 activation, including induction of de novo pyrimidine biosynthesis. Cytidine also reversed multiple features of mitochondrial deficits. In cells expressing DeSI1(S25E) for 7 days, cytidine supplementation restored the levels of SLC25A5 and cardiolipin-sensitive respiratory-chain subunits (Figure 6G, compare lanes 7-8 to 9-10) without altering the corresponding proteins in wild-type cells (Figure 6G, lanes 3-6). Moreover, cytidine restored respiratory capacity in chronically induced DeSI1(S25E) cells: basal and maximal OCR showed significant interactions, indicating that the effect of cytidine depended on DeSI1 genotype, with DeSI1(S25E) cells showing the rescued phenotype (Figure 6H and Supplementary Figure 7C). Spare respiratory capacity also increased with cytidine in DeSI1(S25E)-expressing cells (Supplementary Figure 7C,D). Mitochondrial ultrastructure improved in parallel: chronic DeSI1(S25E) expression reduced matrix density, which was restored by cytidine, with accompanying cytidine-dependent changes in matrix lucency and circularity (Supplementary Figure 7E,F).

Finally, the metabolic rescue extended to cell proliferation. Sustained expression of DeSI1(S25E) reduced cell expansion over 8 days, whereas cytidine restored growth to wild-type levels (Supplementary Figure 7G). This rescue was retained when cytidine was administered together with tetrahydrouridine (THU) to inhibit its deamination to uridine (Figure 6I), supporting a specific requirement for cytidine rather than an indirect effect mediated through uridine.

Together, these results identify cytidine-nucleotide supply-demand imbalance imposed by sustained DeSI1 activity.

### Persistent DeSI1 activation causes cerebellar degeneration with a cytidine-nucleotide/phospholipid imbalance

Experiments in cell systems established that the consequences of DeSI1 activation depend on its duration: transient activation increases the abundance of membrane-targeted proteins and mitochondrial respiratory capacity, whereas sustained activation imposes a cytidine-nucleotide constraint and ultimately leads to mitochondrial decompensation. This raised a physiological question that cycling-cell models cannot fully address: what are the consequences of chronic DeSI1 activation in an intact organism, including in post-mitotic tissues? To address this question, we generated mice carrying constitutive knock-in alleles of either DeSI1(S25E) or DeSI1(S25A), alongside a DeSI1-null allele. All three alleles were expressed throughout the body, allowing the physiological consequences of altered DeSI1 activity – and tissue-specific vulnerability – to emerge without imposing a predefined tissue-specific perturbation.

Intercrosses of DeSI1^S25E/+^ mice yielded homozygous DeSI1^S25E/S25E^ animals at the expected Mendelian frequency (Figure 7A). In contrast, no homozygous DeSI1^S25A/S25A^ offspring were recovered from 94 pups, whereas homozygous DeSI1-null animals were significantly under-represented relative to the expected Mendelian frequency (Figure 7A). Thus, constitutive activation of the DeSI1 switch was compatible with development, whereas preventing S25 phosphorylation was incompatible with recovery of homozygous animals. Complete loss of DeSI1 produced a less severe, but still detectable, developmental phenotype. Consistent with this distinction, surviving DeSI1^−/-^ mice exhibited reduced body weight from weaning through early adulthood, whereas DeSI1^S25E/S25E^ mice were comparable to wild-type littermates (Supplementary Figure 8A).

**Figure 7.**
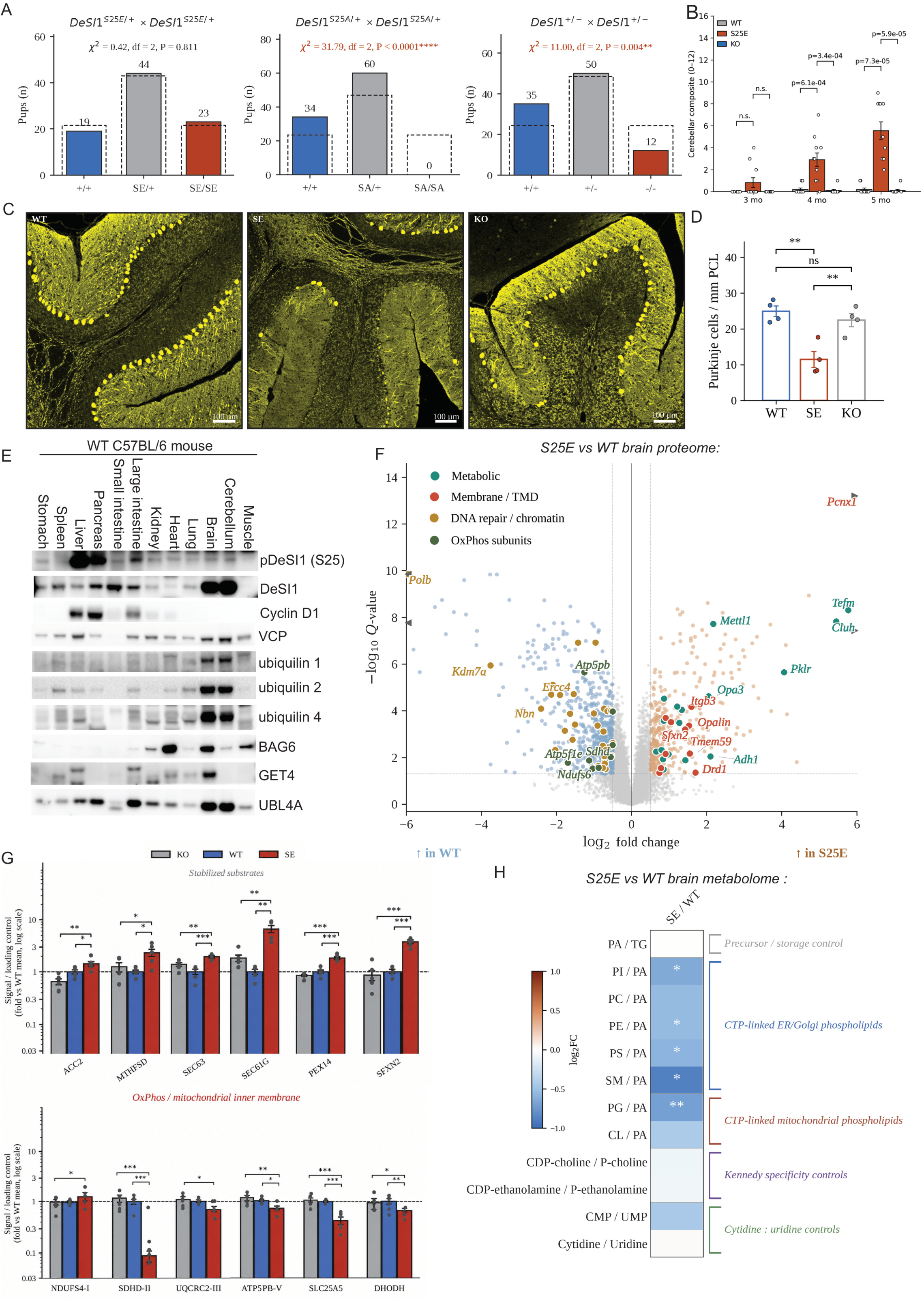
Constitutive DeSI1 phosphorylation causes progressive cerebellar degeneration with a cytidine-nucleotide supply–demand imbalance. **(A)** Mendelian transmission of DeSI1 S25E, S25A and null alleles. Left, *DeSI1*^S25E/+^ × *DeSI1*^S25E/+^ intercrosses (5 cages, 14 litters, n = 86 pups); middle, *DeSI1*^S25A/+^ × *DeSI1*^S25A/+^ intercrosses (5 cages, 14 litters, n = 94 pups); right, *DeSI1*^−/+^ × *DeSI1*^−/+^ intercrosses (6 cages, 17 litters, n = 97 pups). Bars show observed genotype counts and dashed outlines show the expected 1:2:1 Mendelian distribution. *DeSI1*^S25E/S25E^ mice were recovered at the expected frequency (χ² = 0.42, df = 2, P = 0.811), whereas no *DeSI1*^S25A/S25A^ mice were recovered (χ² = 31.79, df = 2, P < 0.0001) and DeSI1-null homozygotes were significantly under-recovered (χ² = 11.00, df = 2, P = 0.004). χ² goodness-of-fit tests were performed against the expected 1:2:1 distribution. **(B)** Progressive cerebellar phenotype. Cerebellar composite score (0–12) at 3, 4 and 5 months in wild-type, S25E and knock-out mice. Bars, mean ± SEM with individual animals shown; P values as indicated; n.s., not significant. Within-animal longitudinal progression in Supplementary Figure 8B. **(C)** Purkinje cell loss in S25E cerebellum. Representative multiplex immunofluorescence of mid-sagittal cerebellar sections from wild-type, S25E and knock-out mice stained for calbindin-D28k (yellow), imaged on the Vectra Polaris/PhenoImager platform. Scale bars, 100 µm. GFAP labelling of the same sections in Supplementary Figure 8E,F. **(D)** Purkinje linear density, scored in QuPath as the number of calbindin-positive somata per mm of traced Purkinje cell layer (PCL). Images were scored using coded identifiers and decoded after scoring; per-region values were collapsed to sections and then to animals, so that the replication unit is the animal. Bars, mean ± SEM with individual animals shown; pairwise Welch two-sided t-tests, uncorrected; **P < 0.01; ns, not significant. Molecular layer thickness from the same sections in Supplementary Figure 8C. **(E)** Tissue distribution of DeSI1 and its interaction machinery. Whole-cell lysates from the indicated tissues of adult wild-type C57BL/6 mice were immunoblotted for pDeSI1 (S25), total DeSI1, cyclin D1, VCP, UBQLN1/2/4, BAG6, GET4 and UBL4A. Ponceau staining in Supplementary Figure 9A. **(F)** Brain proteome. Volcano plot of S25E versus wild-type whole-brain DIA-MS (n = 6 mice per genotype); −log₁₀(q value) versus log₂ fold change. Dashed lines indicate q = 0.05 and |log₂ fold change| = 0.5. Statistical significance was determined by two-sided Welch’s t-tests on median-normalized log₂ protein quantities followed by Benjamini–Hochberg correction within the contrast. Proteins assigned to curated functional categories are shown as filled circles: metabolic, membrane/TMD, DNA repair/chromatin and OxPhos subunits. Totals of significantly changed proteins are inset. The corresponding S25E-versus-knockout contrast is shown in Supplementary Figure 9B. **(G)** Validation of the proteomic dissociation by immunoblot. Densitometric quantification of whole-brain immunoblots shown in Supplementary Figure 9D presented as two plots. Top, proteins increased in S25E brain: ACC2, MTHFSD, SEC63, SEC61G, PEX14 and SFXN2. Bottom, OxPhos/mitochondrial inner-membrane proteins: NDUFS4 (Complex I), SDHD (Complex II), UQCRC2 (Complex III), ATP5PB (Complex V), SLC25A5 and DHODH. Values are signal/loading-control ratios expressed as fold of the WT mean on a log scale; horizontal dashed line, WT mean. Bars, mean ± SEM; points, individual mice. Lower brackets compare S25E with WT and upper brackets compare S25E with KO; two-sided Welch’s t-tests. Only significant comparisons are shown: P < 0.05, P < 0.01, P < 0.001. n = 6 mice per genotype **(H)** Brain phospholipid pathway balance in DeSI1(S25E) mice. Heatmap shows the S25E-versus-WT difference in within-mouse log₂ pathway ratios (S25E n = 5, WT n = 5; WT3 excluded a priori based on cross-platform QC). Lipid-class abundance was calculated per mouse by summing median-normalized linear abundances of de-duplicated annotated species within each class. PA/TG serves as a CTP-independent precursor/storage control. ER/Golgi phospholipids (PI, PC, PE, PS and SM) and mitochondrial phospholipids (PG and CL) are shown relative to PA to assess membrane phospholipid output relative to the precursor pool. CDP-DAG output was additionally evaluated relative to the CTP-independent TG pool: mitochondrial CDP-DAG output/TG represents the geometric mean of PG and CL relative to TG, whereas total CDP-DAG output/TG represents the geometric mean of PI, PG and CL relative to TG. CDP-choline/P-choline and CDP-ethanolamine/P-ethanolamine provide Kennedy-pathway specificity controls, and CMP/UMP and cytidine/uridine assess cytidine-versus-uridine balance. Statistics were calculated using two-sided Welch t-tests on log₂ ratios. Asterisks indicate nominal significance (*P < 0.05, **P < 0.01, ***P < 0.001); Benjamini–Hochberg-adjusted q values across displayed rows and within prespecified pathway groups are reported in the source data.

DeSI1^S25E/S25E^ mice appeared phenotypically normal at 3 months of age but developed a progressive neurological phenotype beginning at 4 months, marked by increased composite neurological scores and declining performance on the balance-beam assay, with both phenotypes becoming more pronounced by 5 months (Figure 7B and Supplementary Figure 8B). In contrast, surviving DeSI1-null animals remained comparable to wild-type littermates throughout this period. Histological analysis of the cerebellum revealed progressive pathology in DeSI1^S25E/S25E^ mice, including disruption and loss of calbindin-positive Purkinje cells, shortened and disorganized dendritic arbors, and reduced molecular-layer thickness (Figure 7C,D and Supplementary Figure 8C). Silver staining at 6 months further confirmed ongoing neurodegeneration, whereas no comparable pathology was detected in wild-type or DeSI1-null littermates (Supplementary Figure 8D). Thus, chronic constitutive activation of DeSI1 causes a progressive, cerebellum-predominant neurodegenerative phenotype that is distinct from, and not recapitulated, by loss of DeSI1 function.

Interestingly, analysis of 12 tissues from wild-type mice revealed marked tissue-specific differences in DeSI1 abundance and phosphorylation. DeSI1 protein was particularly abundant in brain and cerebellum, yet phosphorylated DeSI1 was barely detectable in these tissues (Figure 7E and Supplementary Figure 9A). Conversely, DeSI1 was highly phosphorylated in liver and pancreas despite substantially lower overall protein abundance. Thus, the cerebellum contains abundant DeSI1 but appears to maintain the protein predominantly in its inactive, unphosphorylated state.

We next asked whether the tissue affected in DeSI1^S25E/S25E^ mice recapitulated the molecular signature of chronic DeSI1 activation observed in cultured cells. Proteomic analysis of whole brain, including the cerebellum, revealed a striking partitioning of the proteome: hydrophobic proteins, including membrane-associated proteins and metabolic enzymes, accumulated in DeSI1^S25E/S25E^ brain, whereas proteins involved in oxidative phosphorylation, as well as chromatin/DNA repair proteins were depleted (Figure 7F). The same pattern was observed when DeSI1^S25E/S25E^ brain was compared with DeSI1-null brain (Supplementary Figure 9B), further supporting a gain-of-function effect of the phospho-mimetic allele. Immunoblotting confirmed the accumulation of representative DeSI1-regulated proteins, including ACC2, MTHFSD, SEC63, SEC61G, PEX14, and SFXN2, together with reduced levels of SLC25A5 and multiple OxPhos/mitochondrial inner-membrane proteins (Figure 7G and Supplementary Figure 9D).

Brain lipidomics revealed altered phospholipid partitioning rather than generalized lipid loss. In DeSI1^S25E/S25E^ brain, phosphatidic acid (PA) remained proportional to the CTP-independent triacylglycerol (TG) pool, whereas ER/Golgi and mitochondrial membrane phospholipids showed a coordinated decrease relative to PA. This imbalance was strongest among CDP-DAG-derived phospholipids – phosphatidylinositol (PI) in the ER/Golgi and phosphatidylglycerol (PG) and cardiolipin (CL) in mitochondria – relative to TG (Figure 7H). Within CL, the deficit was greatest among the less unsaturated species, consistent with preferential vulnerability of the newly synthesized CL pool (Supplementary Figure 9E), recapitulating the pattern observed during DeSI1 activation in cells. Together, these findings indicate that CTP-dependent membrane phospholipid synthesis does not scale with expansion of the lipid precursor/storage pool, resulting in a coordinated phospholipid deficit across both ER/Golgi and mitochondrial membranes rather than a defect confined to a single organelle.

The brain proteome showed no coordinated increase in the capacity to replenish the cytidine nucleotide supply. Several CTP-consuming enzymes involved in phospholipid synthesis, including PCYT1A, were increased, whereas key enzymes supporting de novo pyrimidine synthesis, including DHODH, and CTP production, including CTPS2, were reduced (Supplementary Figure 9F). Thus, chronic DeSI1 activation increased demand for CTP-dependent phospholipid synthesis without a corresponding increase— and potentially with a decrease— in the capacity to replenish cytidine-nucleotide resources.

Taken together, the DeSI1^S25E/S25E^ mouse model recapitulates the central features of chronic DeSI1 activation established in cell systems: persistent accumulation of membrane-associated proteins, depletion of mitochondrial respiratory-chain proteins, and selective loss of CTP-dependent phospholipids. Thus, the molecular imbalance defined in cultured cells is recapitulated in the intact organism, with the cerebellum emerging as a site of selective vulnerability.

## Discussion

Our study identifies a previously unrecognized mechanism through which cyclin D-CDK4/6 complexes couple cell-cycle entry to the post-translational supply of membrane-targeted and mitochondrial proteins. We show that DeSI1 is a CDK4/6 substrate whose phosphorylation at S25 converts a latent homodimer into an active monomer, thereby coupling two functions within a single conformational switch: exposure of the catalytic site and creation of a hydrophobic, acidic surface that recognizes proteins bearing hydrophobic regions, such as TMDs and MTSs. This newly exposed surface defines a substrate-recognition surface in DeSI1. Phosphorylated DeSI1 promotes the persistence of newly synthesized proteins bearing these targeting elements, thereby increasing the abundance of proteins involved in membrane-associated processes, mitochondrial protein import, and respiratory-chain function. DeSI1 therefore provides a post-translational output of cyclin D-CDK4/6 signaling that increases the abundance of membrane-targeted and mitochondrial proteins, together with mitochondrial respiratory capacity. When this output is constitutively engaged, however, accumulation of membrane-associated proteins becomes uncoupled from the metabolic resources required to sustain membrane phospholipid synthesis and mitochondrial function, resulting in a cytidine-nucleotide constraint, phospholipid imbalance, and progressive mitochondrial decompensation. In mice, constitutive DeSI1 activation causes progressive cerebellar degeneration accompanied by the same fundamental dissociation between persistent accumulation of membrane-associated proteins and mitochondrial dysfunction. Together, these findings expand the function of cyclin D-CDK4/6 beyond transcriptional preparation for S phase and reveal a direct mechanism through which cell-cycle signaling controls the proteostatic output of membrane-targeted proteins.

Membrane-destined proteins expose hydrophobic elements that can compromise productive folding and targeting, thereby engaging chaperone systems and ubiquitin-dependent quality-control pathways that promote ubiquitylation and degradation^11,12^. Interestingly, we found that unphosphorylated, dimeric DeSI1 associates with components of the BAG6-ubiquilin-VCP machinery, a central component of this quality-control system. Upon phosphorylation, DeSI1 may act near the point at which hydrophobic clients are triaged between productive targeting and degradation, establishing a substrate-proximal ubiquitin cycle that can influence their fate. Specifically, cyclin D-CDK4/6-dependent phosphorylation changes the outcome of nascent-protein triage: phosphorylated DeSI1 reduces ubiquitylation and extends the lifetime of newly synthesized membrane-destined proteins needed for membrane and mitochondrial expansion. The broader implication is that cyclin D-CDK4/6 complexes coordinate distinct forms of biosynthetic control during cell-cycle entry. Through RB phosphorylation, they release E2F-dependent transcriptional programs that prepare cells for cell division and increase nucleotide and other anabolic capacities^4,5,24^. Separately, through phosphorylation of DeSI1, the same kinases post-translationally increase the abundance of membrane- and mitochondrial-targeted proteins. Consistent with these distinct outputs, CDK4/6 activity produces a strong transcriptional response in de novo pyrimidine metabolism, whereas oxidative-phosphorylation capacity is increased preferentially at the protein level. Although this global protein-dominant response may not be attributed entirely to DeSI1, the enrichment of membrane-insertion-prone proteins in both the DeSI1 datasets and the broader CDK4/6 protein response supports a substantial contribution of DeSI1 to this post-translational program. We therefore propose that the RB-E2F and DeSI1 pathways are complementary outputs of cyclin D-CDK4/6 activity, with one arm increasing biosynthetic resource supply and the other increasing the supply of membrane-targeted and mitochondrial proteins. The DeSI1(S25E) allele, by constitutively engaging the DeSI1 arm without equivalently activating the broader cyclin D-CDK4/6 program, exposes the importance of this coordination. Major phospholipid-synthetic pathways consume CTP through CDP-activated intermediates, creating a direct link between increased membrane-associated protein supply and cytidine-nucleotide demand. Several independent observations indicate that sustained DeSI1 activation produces a cytidine-nucleotide supply-demand imbalance. The cellular cytidine pool decreases, while phosphatidylglycerol accumulates as cardiolipin falls, consistent with a bottleneck in cardiolipin synthesis. Perturbation of de novo pyrimidine synthesis selectively compromises cells expressing the phospho-mimetic DeSI1 mutant, whereas cytidine supplementation restores mitochondrial function and proliferation. Notably, the proliferative rescue persists when cytidine deamination to uridine is inhibited, indicating that cytidine itself, rather than its conversion to uridine, supports the rescue.

Because CTP and UTP were not resolved by our metabolomic platform, these experiments do not establish direct depletion of CTP itself. Rather, they support a cytidine-nucleotide supply-demand imbalance that impairs CTP-dependent phospholipid synthesis. Cardiolipin appears particularly sensitive to this imbalance because its synthesis requires two CTP-derived CDP-DAG equivalents^22^. Its depletion is already evident while respiratory capacity remains increased, indicating that the lipid abnormality precedes the subsequent loss of respiratory-chain components and mitochondrial function. Given the established requirement for cardiolipin in respiratory-complex organization^25,26^, this temporal sequence is consistent with cardiolipin depletion contributing to mitochondrial decompensation, although the present experiments do not establish cardiolipin loss as the sole causal intermediate. With sustained activation *in vivo*, however, the lipid phenotype extends beyond cardiolipin: phospholipids across both ER/Golgi and mitochondrial branches that depend on CDP-DAG are depleted, whereas the upstream phosphatidic-acid and CTP-independent triacylglycerol pools remain comparatively preserved. Together, the acute and chronic phenotypes are consistent with a common underlying metabolic liability, with cardiolipin providing an early and particularly sensitive readout, while prolonged DeSI1 activation reveals a broader failure of membrane phospholipid synthesis to keep pace with the increased abundance of membrane-associated proteins.

The DeSI1(S25E) mouse extends this principle to the organismal level. Constitutive DeSI1 activation caused progressive cerebellar degeneration that was absent in DeSI1-null mice, indicating that this phenotype reflects a gain of DeSI1 activity. The affected cerebellum recapitulated the molecular signature of chronic DeSI1 activation seen in cells, including accumulation of membrane-associated proteins, loss of respiratory-chain components, and depletion of CTP-dependent phospholipids, including cardiolipin, thereby reproducing the association between phospholipid imbalance and mitochondrial dysfunction observed in cell systems. Together, these findings suggest that long-term cellular homeostasis depends on coordinating membrane-associated and mitochondrial protein supply with the metabolic resources required to maintain membrane phospholipid composition and mitochondrial function. Like for many other mouse models, the molecular and cellular basis for the tissue selectivity remains unresolved^27,28^. Brain and cerebellum contain abundant DeSI1 while maintaining relatively little phosphorylated DeSI1, suggesting that these tissues may be particularly dependent on restraining DeSI1 activation. The extensive membrane architecture, high mitochondrial density, and energetic demands of Purkinje cells, together with their documented vulnerability to respiratory-chain dysfunction^29,30^ may further increase their susceptibility to the chronic imbalance produced by constitutive DeSI1 activity.

The differential outcomes of the knock-in alleles suggest that DeSI1 function depends not simply on protein presence, but on access to the S25-regulated state transition. Indeed, the lethality of the DeSI1^S25A/S25A^ genotype, together with the reduced viability and body weight of DeSI1-null mice, indicates that both the phosphorylation-dependent regulation and the basal activity of DeSI1 are important for normal development. In contrast, the viability and largely normal growth of DeSI1^S25E/S25E^ mice suggest that constitutive activation of DeSI1 is compatible with development, although its physiological consequences may emerge later in life and in a tissue-specific manner.

Together, our study identifies a phosphorylation-gated mechanism that connects canonical G1/S kinases to the proteostatic control of membrane-targeted and mitochondrial proteins. By altering the fate of nascent hydrophobic proteins, cyclin D-CDK4/6 complexes can rapidly increase the abundance of protein machinery that supports membrane and organelle expansion. Constitutive engagement of this program reveals its metabolic cost: increased membrane and organelle protein demand must remain matched to cytidine-nucleotide-dependent phospholipid synthesis and the other biosynthetic resources required to sustain it. More broadly, our findings suggest that proliferative signaling couples cell-cycle progression to cell growth through coordinated control of resource provisioning and protein stability. When these outputs become uncoupled, an anabolic program that normally supports growth can instead become a metabolic liability.

### Limitations of the study

This study defines a phosphorylation-gated function for DeSI1, but several mechanistic questions remain. First, although DeSI1 was originally characterized as a deSUMOylase^31^, our data indicate that the phosphorylated enzyme instead acts as a ubiquitin-directed protease. The apparent discrepancy may reflect the latent dimeric state of DeSI1, in which the catalytic cysteine and substrate cleft are occluded until phosphorylation of S25. However, because we did not detect deSUMOylase activity even with the phospho-mimetic form of DeSI1, the basis for this discrepancy remains unresolved. Second, we showed that DeSI1 is phosphorylated selectively during the G1 interval of rising cyclin D-CDK4/6 activity and then dephosphorylated at the G1/S boundary, likely confining the pulse of membrane and mitochondrial protein accumulation to G1. Yet the mechanism that reverses S25 phosphorylation, including the relevant phosphatase, is still unknown. Third, although the proposed client-recognition surface is supported by structural modeling, peptide-binding measurements, and substrate-selective cleavage, this model awaits direct validation by an experimental structure of phosphorylated DeSI1 bound to a ubiquitylated client. Our GPS reporter assay also has inherent limits: it isolates hydrophobic-sequence-dependent effects on protein stability, but it does not fully capture the endogenous context of membrane-protein targeting and quality control. Finally, although the brain lipidomic and proteomic phenotypes recapitulate the biochemical signature of the cytidine-nucleotide supply-demand imbalance defined functionally in cells, nucleotide flux through CTP-dependent phospholipid synthesis was not directly measured in vivo. Together, these limitations highlight the remaining mechanistic gaps while leaving intact the central model supported by our data: CDK4/6-dependent phosphorylation activates DeSI1 as a ubiquitin-directed protease, thereby linking cell-cycle progression to proteostatic and metabolic homeostasis.

## MATERIALS AND METHODS

### Cell lines and cell culture

HEK293T (ATCC CRL-3216), U2OS (ATCC HTB-96), HCT-116 (ATCC CCL-247), and hTERT RPE-1 (RPE-hTERT; ATCC CRL-4000) cells were maintained at 37 °C in a humidified atmosphere containing 5% CO₂. HEK293T and RPE-hTERT cells were cultured in Dulbecco’s modified Eagle’s medium (DMEM), whereas U2OS and HCT-116 cells were cultured in McCoy’s 5A medium. Unless otherwise indicated, media were supplemented with 10% fetal bovine serum (FBS) and penicillin/streptomycin/L-glutamine. Cells carrying doxycycline-inducible constructs were maintained in medium containing Tet System Approved FBS (Takara/Clontech).

HEK293F suspension cells (Gibco R79007), used for denaturing His-ubiquitin pulldowns, were maintained in FreeStyle 293 Expression Medium on an orbital shaker at 37 °C and 8% CO₂. HCT-116 cells carrying endogenous 2×FLAG-mAID-AMBRA1 together with HA-OsTIR1 were described previously^16^. Cell cultures were routinely tested for mycoplasma contamination.

RPE-hTERT cells stably depleted of endogenous DeSI1 were generated using a PiggyBac-ased shRNA construct targeting the DESI1 3′ UTR (target sequence, 5′-CTAAGCGTCCTCCACACTGG-3′). Where indicated, a scrambled shRNA targeting sequence (5′-CCTAAGGTTAAGTCGCCCTCG-3′) was used as the control. DeSI1 depletion was verified by immunoblotting.

Doxycycline-inducible DeSI1 reconstitution lines were generated using the TetOne-RFP-FLAG system. Unless otherwise specified, inducible DeSI1 and cyclin D1(T286A) constructs were induced with doxycycline at 1 µg/mL.

### Acute and chronic induction regimens

For doxycycline-inducible DeSI1 and cyclin D1(T286A) experiments, acute induction was defined as 24 h of doxycycline treatment, whereas chronic induction was defined as 7 days of continuous doxycycline treatment. Where indicated, cytidine was present throughout the induction period. THU was added together with cytidine in experiments designed to inhibit cytidine deamination, and uridine was used in the indicated control conditions. Where indicated, cytidine was added at 200 µM. Tetrahydrouridine (THU; 100 µM) was co-administered with cytidine in experiments designed to inhibit cytidine deamination, and uridine was used at 100 µM in the indicated control conditions.

For metabolomic and lipidomic experiments, doxycycline treatment was timed such that the inducible constructs were examined under the acute, 24-h induction regimen. Chronic respiratory, immunoblotting, ultrastructural, and proliferation experiments used the 7-day induction regimen.

### Plasmids and site-directed mutagenesis

Human DeSI1 wild type and the S25A, S25E, C108S, S25E/C108S, C58A, S25E/C58A, L20E/L24E, F140S, L144R, and Δ151–163 variants were cloned into the indicated mammalian or bacterial expression vectors, including modified pcDNA3.1, pBABE-puro, pLenti-TetOne, and pET-21b backbones. Point mutations were introduced by site-directed mutagenesis, and constructs were verified by Sanger sequencing.

Cyclin D1, cyclin D1(T286A), CDK4, and CDK6 were expressed from modified pcDNA3.1 vectors carrying an N-terminal tandem 2×FLAG–2×Strep (FFSS) tag, as described previously^16,32^. Cyclin D1(T286A) was additionally cloned into a doxycycline-inducible lentiviral vector. HA-p107 was used as a positive control for Phos-tag analysis. The use of modified pcDNA3.1, pBABE and lentiviral vectors and the FFSS designation follows the established laboratory nomenclature.

Split-GFP constructs consisted of GFP11-mRFP-DeSI1 and DeSI1-GFP1–10 fusion proteins separated by flexible linkers. GPS reporters were based on a bicistronic EGFP-IRES-mCherry stability reporter and contained the indicated hydrophobic or matched hydrophilic segments derived from SLC25A5, MFN2, DHCR7, and HMGCR; CA2-derived sequence and unfused GFP served as soluble controls. His₆-HA-ubiquitin was used for denaturing ubiquitin-conjugate capture.

### Transient transfection and RNA interference

HEK293T and HEK293F cells were transiently transfected using linear 25-kDa polyethylenimine (PEI; Polysciences), whereas U2OS, HCT-116, and RPE-hTERT cells were transfected using Lipofectamine 3000 (Thermo Fisher Scientific) according to the manufacturer’s instructions.

siRNA transfections were performed using Lipofectamine RNAiMAX (Thermo Fisher Scientific) at a final siRNA concentration of 20 nM. ON-TARGETplus SMARTpool siRNAs targeting *DESI1* and a non-targeting control pool were used as indicated. Cells were analysed 48–72 h after transfection.

### Viral transduction and stable line generation

Lentiviral and retroviral particles were produced in HEK293T cells by co-transfection of the appropriate transfer and packaging plasmids. Virus-containing medium was collected 48– 72 h after transfection, cleared through a 0.45-µm filter, supplemented with 8 µg/mL polybrene, and applied to target cells. Stable populations were selected using puromycin, blasticidin, or hygromycin B, as appropriate for the construct, or isolated by fluorescence-activated cell sorting. Doxycycline-inducible RFP-positive populations were sorted where required to obtain uniform transgene-expressing populations.

*DESI1*-knockout HEK293T and U2OS cells were generated by CRISPR–Cas9 editing using a *DESI1*-targeting sgRNA with the spacer sequence 5′-TGTCCAAAGGCCTGGCCCGG-3′. Following editing and selection, single-cell clones were isolated and screened for loss of DeSI1 protein by immunoblotting. Editing of the targeted locus was verified by Sanger sequencing.

### Cell-cycle synchronization and G1 release

For synchronization in G0, RPE-hTERT or U2OS cells were cultured under serum-free conditions for 48 h for GPS reporter or AHA pulse-chase experiments and for 72 h for metabolomic, lipidomic, and extracellular-flux experiments. Cells were released into G1 by replacement with complete medium containing 10% FBS and collected at the indicated times. Where indicated, cycloheximide (100 µg/mL) or MG132 (10 µM) was added at serum re-addition. Cell-cycle progression was monitored by immunoblotting for cyclin D1, cyclin E1, cyclin A2, and RB phosphorylation and, where indicated, by flow cytometry.

### Immunoblotting

Cells were lysed in RIPA buffer supplemented with protease and phosphatase inhibitors (cOmplete ULTRA and PhosSTOP, Roche) and, where preservation of ubiquitin conjugates was required, N-ethylmaleimide (NEM). Protein concentrations were determined by BCA assay. Lysates were resolved by SDS-PAGE and transferred to PVDF membranes. Membranes were blocked in 5% non-fat dry milk in TBS-T; phospho-specific antibodies were incubated in BSA-containing blocking buffer. HRP-conjugated secondary antibodies were detected by enhanced chemiluminescence using a Bio-Rad ChemiDoc imaging system. Band intensities were quantified in Fiji/ImageJ and normalized to the loading control indicated in the corresponding figure legend.

For immunoblotting of mitochondrial respiratory-chain and other highly hydrophobic membrane proteins, samples were incubated at 37 °C rather than boiled before electrophoresis to minimize heat-induced aggregation.

### Phos-tag SDS-PAGE

Phosphorylation-dependent mobility shifts were analysed by Phos-tag SDS–PAGE essentially as described previously^33^. Proteins were resolved on 7.5% polyacrylamide Phos-tag gels containing 40 µM Zn²⁺-Phos-tag reagent (FUJIFILM Wako Chemicals) at 50 V according to the manufacturer’s instructions. Following electrophoresis, gels were processed for transfer according to the manufacturer’s protocol and proteins were transferred to PVDF membranes for immunoblotting. Parallel conventional SDS–PAGE was performed from the same lysates to assess total protein abundance.

### Generation and validation of the phospho-DeSI1 (S25) antibody

A phospho-specific rabbit polyclonal antibody against DeSI1 phosphorylated at Ser25 was generated by YenZym Antibodies (South San Francisco, CA) using an approach analogous to that described for the phospho-GTSE1 antibody^33^. A phosphopeptide encompassing the DeSI1 S25 phosphorylation site (C-Ahx-LARRL-pSer-PIMLG-NH₂) was synthesized and used for rabbit immunization following conjugation to KLH. Phospho-specific antibodies were subsequently purified using a phosphopeptide-conjugated affinity matrix, and antibody reactivity was assessed by ELISA before and after purification. Antibody specificity was validated experimentally using multiple independent controls. The antibody recognized wild-type DeSI1 following co-expression of cyclin D1–CDK4 but showed no corresponding signal with the phospho-deficient DeSI1(S25A) mutant; the cyclin D1–CDK4-dependent signal was suppressed by palbociclib. Specificity was further assessed in DeSI1-knockout HEK293T cells. The antibody also recognized the phosphomimetic DeSI1(S25E) mutant in cells and recombinant DeSI1(S25E) purified from bacteria

### Immunoprecipitation

For immunoprecipitation, cells were lysed in buffer containing 50 mM Tris-HCl (pH 7.5), 150 mM NaCl, 0.2% NP-40, 10% glycerol, 1 mM EDTA, 1 mM EGTA and 2 mM MgCl₂ supplemented with protease and phosphatase inhibitors. Where preservation of ubiquitylated species was required, N-ethylmaleimide was included. Lysates were clarified by centrifugation at 20,000 × *g* for 15 min at 4 °C and incubated with FLAG-M2 magnetic beads or anti-HA affinity resin, as appropriate, for 2 h at 4 °C with rotation. Immunoprecipitates were washed extensively with lysis buffer containing 150 mM NaCl and 0.2% NP-40. Bound proteins were eluted with 3×FLAG peptide for native purification or mass-spectrometry applications, or directly in Laemmli sample buffer for immunoblot analysis.

### In vitro kinase assays

Kinase reactions were performed in 25 mM Tris-HCl (pH 7.5), 10 mM MgCl₂, 5 mM β-glycerophosphate, 2 mM DTT and 0.1 mM Na₃VO₄. Recombinant human cyclin D1–CDK4 (Abcam, ab55695) was activated before use by incubation at 2 µM with 200 nM GST-tagged CDK-activating kinase (CAK; CDK7–cyclin H–MAT1; Sigma-Aldrich, 14-476-M) in kinase buffer containing 1 mM ATP for 1 h at 30 °C. GST-CAK was subsequently removed by incubation with glutathione-Sepharose for 1 h at 4 °C, and the supernatant containing activated cyclin D1–CDK4 was recovered for kinase reactions.

Kinase reactions contained 1 µg purified FLAG-DeSI1 or GST-RB1 (residues 773–928) and activated cyclin D1–CDK4 at the indicated concentration (40 or 120 nM) in kinase buffer supplemented with 2 mM ATP, 20 mM creatine phosphate and 0.1 mg/mL creatine phosphokinase, in a final volume of 50 µL. Reactions were incubated for 12 h at 30 °C. Where indicated, palbociclib (1 µM) was pre-incubated with the kinase for 10 min on ice before addition of substrate. Reactions were terminated by addition of Laemmli sample buffer, heated at 95 °C for 5 min, resolved by SDS-PAGE and analysed by immunoblotting.

### Quantitative TMT phosphoproteomics

Two independent quantitative TMT phosphoproteomics screens were performed using the same HCT-116 mAID-AMBRA1 system at the Broad Institute Proteomics Platform (Screen 1; RRID:SCR_007073) and the NYU Langone Proteomics Laboratory (Screen 2; RRID:SCR_017926). HCT-116 cells carrying endogenous 2×FLAG-mAID-AMBRA1 and doxycycline-inducible *Oryza sativa* TIR1 were generated as described previously^16^. In this system, doxycycline induction of osTIR1 followed by auxin treatment results in rapid degradation of endogenous AMBRA1 and accumulation of D-type cyclins. Simoneschi et al.^16^ describe doxycycline treatment for 12 h followed by auxin for 4 h in this cell line.

For the phosphoproteomic experiments, osTIR1 expression was induced with doxycycline (0.4 µg/mL) for 12 h, followed by treatment with auxin (indole-3-acetic acid; 0.1 mM) for 4 h. Where indicated, palbociclib (1 µM) was added during the final 2 h of auxin treatment. Cells were harvested at the end of the 4-h auxin treatment. Three independent biological experiments were performed. Screen 1 additionally contained an untreated control arm; the analyses reported in Figure 1 compare auxin+doxycycline+palbociclib with auxin+doxycycline. The Broad quantitative output contains three samples for each of the untreated, Aux+Dox, and Aux+Dox+palbociclib groups.

Cell pellets were processed for TMT-based global proteomic and phosphoproteomic analysis by the respective proteomics facilities. Proteins were denatured, reduced, alkylated and proteolytically digested, and phosphopeptides were enriched before TMT labelling, fractionation and LC-MS/MS analysis. Because the two screens were performed independently at different facilities, each dataset was processed using the facility-specific analytical pipeline.

For Screen 1, peptide and protein identification and quantification were performed using the Spectrum Mill-based workflow of the Broad Institute Proteomics Platform, and differential analysis was performed using Protigy. Differential analysis for Screen 1 used a two-sample moderated t-test implemented in limma/Protigy. Screen 2 was processed using Proteome Discoverer 2.5 and differential analysis was performed in Perseus.

### Differential analysis of the phosphoproteomic screens

For both phosphoproteomic screens, the principal contrast was auxin+doxycycline+palbociclib versus auxin+doxycycline. Log₂ fold change was calculated as the difference in mean log₂ abundance between the palbociclib-treated and untreated arms; therefore, negative values indicate decreased phosphorylation or protein abundance following CDK4/6 inhibition. Features with complete quantitative measurements in the samples contributing to each comparison were retained for analysis, yielding 22,717 phosphosites and 9,455 protein groups in Screen 1 and 11,061 phosphopeptides and 8,036 protein groups in Screen 2.

Screen 1 was analysed using a two-sample moderated *t*-test with empirical-Bayes variance shrinkage implemented in the limma-based Protigy workflow. Screen 2 was analysed using the two-sample *t*-test implemented in Perseus. All reported *P* values are two-sided nominal values. False-discovery rates (*q* values) were calculated independently within each dataset using the Benjamini–Hochberg procedure, and *q* < 0.05 was used as the significance threshold.

Volcano plots display log₂ fold change versus −log₁₀(*P*), with features meeting *q* < 0.05 classified by the direction of change. For Supplementary Figure 1C, phosphoproteomic features were ranked by nominal *P* value and the 15 highest-ranking features from each screen were displayed together with the corresponding global-proteome measurements, where available.

Because the two screens were generated and processed independently, statistical significance was evaluated separately within each dataset and P values were not quantitatively compared or combined across screens. The two screens were instead considered independent experiments providing convergent evidence for CDK4/6-regulated phosphorylation.

Custom analyses and figure generation were performed in Python using pandas, NumPy, SciPy and Matplotlib. Analysis scripts and per-feature source-data tables are provided with the accompanying source data.

### Interactome analysis by immunoprecipitation–mass spectrometry

HEK293T cells expressing FLAG-tagged DeSI1 WT or S25E were harvested 24 h after transfection and subjected to anti-FLAG immunoprecipitation as described above. Samples were processed for LC-MS/MS, and three independent biological replicates were analyzed per bait. PSM measurements were depth-normalized using median-of-ratios size factors calculated from eligible proteins detected in all six runs and centered to a geometric mean of 1. Differential abundance was expressed as log2[(mean normalized S25E PSM + 3)/(mean normalized WT PSM + 3)]. For statistical testing, normalized PSM values were transformed as log2(PSM + 1) and analyzed using a SAM/Perseus-style regularized statistic, with the regularization parameter defined from the median pooled standard deviation across eligible quantified proteins. Two-sided reference P values were calculated using four degrees of freedom. Functional-module colors in the volcano plot were assigned from the curated functional classifications provided in the accompanying source data.

### Denaturing His-ubiquitin pulldown (ubiquitylome)

HEK293F suspension cells were co-transfected with 6×His–HA–ubiquitin together with empty vector (EV), DeSI1(S25E), or DeSI1(S25E/C108S) and harvested 24 h later. Ubiquitin conjugates were isolated by Ni-NTA affinity capture under denaturing conditions. Cells were lysed in 8 M urea, 100 mM NaH₂PO₄, 10 mM Tris-HCl pH 8.0, 10 mM imidazole and 10 mM N-ethylmaleimide, sonicated and cleared by centrifugation. His-tagged conjugates were captured on Ni-NTA resin, washed under denaturing conditions, eluted and analyzed by LC-MS/MS. Three independent biological replicates were analyzed per condition. The primary Figure 3 analysis used samples not treated with MG132.

To ensure that comparisons with the catalytically inactive S25E/C108S mutant were performed on a common quantified protein universe, proteins were retained only when quantified in at least two of three biological replicates in each of EV, S25E and S25E/C108S. Protein groups were resolved to gene-level entries before ranking, yielding 4,531 proteins. Proteins were ranked by the S25E-versus-EV t statistic, with increasingly negative values indicating reduced His-Ub recovery in S25E cells. The proteins in the lowest 20% of this ranking were defined as the S25E-associated reduced-ubiquitylation tail used in Figure 3C. Targeted membrane enrichment within this tail was assessed for six prespecified Gene Ontology cellular-component categories: outer membrane, endoplasmic-reticulum membrane, endoplasmic-reticulum exit site, coated-vesicle membrane, vesicle membrane and Golgi membrane. Category membership was defined from the GO Cellular Component 2026.1 gene sets. Enrichment of lowest-20% membership relative to the complete 4,531-protein universe was tested using one-sided Fisher exact tests, with Benjamini–Hochberg correction across the six prespecified hypotheses. The union of these categories contained 193 unique proteins in the lowest-20% tail.

Catalytic reversal was evaluated independently across the complete 907-protein S25E-associated reduced-ubiquitylation tail using the S25E/C108S-versus-S25E t statistic. A positive t statistic was used as a permissive directional-reversal criterion. Proteins satisfying this criterion were organized according to the functional modules independently identified in the S25E interactome, yielding 165 unique proteins across mitochondrial matrix, mitochondrial membrane, endoplasmic-reticulum, vesicle-membrane and organelle-bounding-membrane modules. Because proteins can belong to more than one module, module assignments were non-exclusive. As a more stringent sensitivity analysis, catalytic reversal was also defined as t(S25E/C108S versus S25E) ≥ 1.5, yielding 74 unique proteins.

#### Membrane-insertion propensity analysis

Membrane-insertion propensity was quantified for sequence-resolved interactome proteins using the Hessa biological hydrophobicity scale^19^. Each protein sequence was scanned with overlapping 19-residue windows, and the minimum window value was retained as the minimum-window ΔG_app score; lower values indicate a more membrane-insertion-favorable sequence.

Proteins with log_2_(fold change)> 1 and *P* < 0.05 were classified as S25E-enriched, whereas proteins with log_2_(fold change)< −1 and *P* < 0.05 were classified as WT-enriched. Within the sequence-resolved analysis set used for Figure 3B, these criteria yielded 109 S25E-enriched and 58 WT-enriched proteins.

For visualization, minimum-window ΔG_app distributions were represented as moving-window relative-frequency curves. The x axis comprised 400 evenly spaced positions spanning the observed minimum-window ΔG_app range with 4% padding at each end. At each position, frequency was defined as the fraction of proteins whose minimum-window ΔG_app fell within ±0.1242 kcal mol^−1^ of that value; this half-width corresponds to 8% of the plotted x-axis range. The final Figure 3B panel displays only the S25E-enriched and WT-enriched distributions.

The directional shift toward lower minimum-window ΔG_app values in S25E-enriched interactors was assessed using a one-sided Mann–Whitney *U*test, with rank-biserial correlation reported as the effect-size measure. Robustness to differences in protein length and abundance was evaluated separately by matched permutation analysis of the combined S25E- and WT-enriched sets using 20,000 permutations, eight protein-length strata and three abundance strata. As a sensitivity analysis, enrichment of proteins below a series of minimum-window ΔG_app cutoffs was additionally examined across the prespecified ΔG_app sweep; this sweep was used to assess robustness and was not used to select the Figure 3B display parameters.

For the ubiquitylome analysis in Supplementary Figure 3F, proteins in the lowest 20% of the S25E-versus-EV His-Ub ranking were compared with the oppositely ranked 20% and with the complete quantified universe. Length dependence was controlled by permutation within eight protein-length strata using 20,000 permutations. For Supplementary Figure 3G, the top 10% of proteins showing the strongest protein-dominant response to CDK4 activity were compared with the remaining 90% using the same minimum-window ΔG_app framework and length-stratified permutation analysis.

### Matched transcriptome and proteome analysis of CDK4/6 activity

#### RNA sequencing and analysis

HCT-116 cells carrying endogenous 2×FLAG-mAID-AMBRA1 and OsTIR1 were treated as described above to generate CDK4-active conditions following AMBRA1 degradation and matched CDK4/6-inhibited conditions following palbociclib treatment, using the AMBRA1-degron system described previously^16^. Total RNA was submitted to the NYU Langone Genome Technology Center (GTC) for library preparation and sequencing. Stranded total-RNA libraries with ribosomal-RNA depletion were prepared using the Illumina TruSeq Stranded Total RNA workflow and sequenced on an Illumina NovaSeq 6000. Sequence processing followed the established NYU GTC/Applied Bioinformatics Laboratory workflow described previously^34^: FASTQ files were generated with bcl2fastq, reads were mapped to the human reference genome (GRCh38) using STAR, sample contamination was assessed with Fastq Screen, gene-level counts were generated with featureCounts, and differential expression between CDK4-active and palbociclib-treated samples was analyzed with DESeq2. Three biological replicates were analyzed per condition.

#### Proteome

The matched global-proteome dataset was generated and processed as described under *Quantitative TMT phosphoproteomics*, using the same CDK4-active and palbociclib-treated samples.

### Integrated transcriptome–proteome pathway analysis

Matched transcriptomic and global-proteomic datasets from HCT-116 mAID-AMBRA1 cells were analyzed under CDK4/6-active conditions following AMBRA1 degradation and after acute CDK4/6 inhibition with palbociclib. Differential responses were oriented such that positive values indicate increased RNA or protein abundance in the CDK4/6-active state. Preranked gene-set enrichment analysis (GSEA) was performed independently on the RNA and protein differential-response rankings using the MSigDB C5 Gene Ontology Biological Process (GO BP) and Cellular Component (GO CC) collections (human gene-symbol sets, v2026.1). All eligible GO BP and GO CC terms were tested and retained in the source data; pathways were not preselected on the basis of biological theme. For each GO term, the normalized enrichment score (NES) obtained from the RNA analysis was matched to the corresponding NES from the proteomic analysis and plotted in Figure 3D, with the diagonal representing equivalent responses in the two molecular layers.

To formally assess whether individual biological programs were preferentially regulated at the protein or transcript level, the analysis was restricted to genes quantified in both datasets. Gene-level responses were standardized separately within the RNA and protein datasets, and a differential-layer score was calculated as ΔZ = Z(protein response) − Z(RNA response). Genes were ranked by ΔZ and subjected to preranked GSEA using the same GO BP and GO CC collections. The differential-layer analysis used 1,000 permutations. Positive differential NES values indicate protein-dominant regulation, whereas negative values indicate transcript-dominant regulation. An FDR q < 0.10 was used as the discovery threshold for direct differential enrichment. Filled labeled terms in Figure 3D satisfy this criterion, whereas open labeled terms denote contextual or trending programs from the same unbiased analysis that did not meet the direct differential threshold. Labels represent nonredundant biological themes selected for readability and do not define the statistical test universe. The displayed “Pyrimidine nucleotide biosynthesis” label corresponds to GOBP_PYRIMIDINE_NUCLEOSIDE_MONOPHOSPHATE_BIOSYNTHETIC_PROCESS; broader pyrimidine-related terms were retained in the complete source-data table. Full ranked inputs and GO-level enrichment statistics are provided in the source data.

### Functional annotation and pathway enrichment

Functional organization of the WT- and S25E-enriched interactomes in Supplementary Figure 3B–C was based on Gene Ontology enrichment followed by grouping of representative proteins under the major enriched functional terms; complete memberships and adjusted enrichment statistics are provided in the source data. The targeted ubiquitylome enrichment in Supplementary Figure 3D was analyzed separately and was not an unrestricted GO-enrichment screen: six membrane-associated cellular-component terms were prespecified from the membrane systems highlighted by the independent S25E interactome and tested by one-sided Fisher exact test against the common 4,531-protein His-Ub universe, with Benjamini–Hochberg correction across those six hypotheses. RNA and protein pathway responses in Figure 3D were evaluated by gene-set enrichment analysis as described above.

#### Recombinant protein expression and purification

His₆-tagged DeSI1 variants were expressed from pET-21b in *Escherichia coli* BL21(DE3). Cultures were grown at 37 °C to OD₆₀₀ of approximately 0.6–0.8 and protein expression was induced with 0.5 mM IPTG overnight at 18 °C. Bacterial pellets were resuspended in 50 mM Tris-HCl (pH 8.0), 300 mM NaCl and 10 mM imidazole supplemented with protease inhibitors and lysed by sonication. Clarified lysates were applied to Ni-NTA agarose, washed, and bound proteins were eluted with 250 mM imidazole. Proteins were exchanged into 20 mM HEPES (pH 7.5), 150 mM NaCl and 1 mM DTT and further purified by size-exclusion chromatography. Purity was assessed by SDS-PAGE and Coomassie staining. Purified proteins were aliquoted, snap-frozen and stored at −80 °C.

### Analytical size-exclusion chromatography

Analytical size-exclusion chromatography was performed on an ÄKTA Pure system (Cytiva) using a Superdex 75 Increase 10/300 GL column equilibrated in 20 mM HEPES (pH 7.5), 150 mM NaCl and 1 mM DTT at a flow rate of 0.5 mL/min. Apparent molecular masses were assigned using a calibration curve generated with globular protein standards. Fractions were analysed by SDS-PAGE followed by Coomassie staining or immunoblotting, as indicated. For analysis of disulfide-crosslinking reactions, DTT was omitted from the running buffer and samples were analysed under reducing and non-reducing conditions.

### Split-GFP complementation assay

U2OS *DESI1*-knockout cells were co-transfected with GFP11-mRFP-DeSI1 (WT, S25A, S25E, C108S or S25E/C108S) and DeSI1-GFP1–10. GFP11 alone served as the fragment-only negative control. Following expression, cells were analysed by flow cytometry and DeSI1 self-association was quantified from GFP complementation as FITC fluorescence within the RFP-positive population. Three independent biological experiments were performed.

### Activity-based probe labelling and modifier specificity assays

Purified recombinant DeSI1 WT, S25A, S25E, C108S and S25E/C108S proteins were incubated with HA-ubiquitin-vinyl sulfone (HA-Ub-VS), HA-SUMO1-vinyl sulfone, biotin-SUMO2-vinyl methyl ester (VME), or FLAG-NEDD8-vinyl sulfone under reducing conditions. Reactions were terminated in Laemmli sample buffer and analysed by SDS-PAGE. Covalent probe–enzyme adducts were detected by immunoblotting with antibodies against the corresponding probe tag, and parallel gels were silver-stained to assess protein input. USP7, SENP1/SENP2, and SENP8 served as positive controls for ubiquitin-, SUMO-, and NEDD8-directed probe reactivity, respectively. For deSUMOylase assays, SUMO2 polychains were incubated with the indicated DeSI1 variants or SENP1/SENP2 and chain cleavage was assessed by SDS-PAGE and silver staining.

### Disulfide crosslinking with ubiquitin G76C

To restrict crosslinking to the catalytic cysteine C108, DeSI1 proteins used in this assay additionally carried the C58A substitution. Ubiquitin G76C was activated with a fivefold molar excess of 5,5′-dithiobis-(2-nitrobenzoic acid) (DTNB; Ellman’s reagent) in 50 mM HEPES (pH 7.5), 150 mM NaCl for 30 min at room temperature, followed by removal of excess DTNB by buffer exchange. Activated ubiquitin G76C was mixed with DeSI1(C58A) or DeSI1(S25E/C58A) at a 1.5:1 molar ratio in 50 mM HEPES (pH 7.5), 150 mM NaCl and incubated overnight at 4 °C. Reactions were quenched with 10 mM NEM and analysed by non-reducing SDS-PAGE; parallel aliquots were reduced in Laemmli sample buffer containing 5% β-mercaptoethanol. Crosslinking reactions were additionally resolved by analytical size-exclusion chromatography, and collected fractions were analysed under reducing and non-reducing conditions by SDS-PAGE and immunoblotting for DeSI1 and ubiquitin.

### Assembly of resin-anchored polyubiquitin chains

Assembly of GST-Ub1–75-anchored polyubiquitin chains. Polyubiquitin chains were assembled on GST-Ub1–75, which lacks the C-terminal Gly76 and therefore serves as a non-cleavable ubiquitin anchor. For K48-directed chain assembly, GST-Ub1–75 was incubated with UBA1, an AMFR(RING)–UBE2G2 fusion protein, and His-tagged ubiquitin K63R. For K63-directed chain assembly, GST-Ub1–75 was incubated with UBA1, UBE2N/UBE2V2, and ubiquitin K48R. Reactions contained 20 mM HEPES, ATP/MgCl2, 1 mM DTT, and 100 µM soluble ubiquitin and were incubated at 37 °C for 4 h. Following chain assembly, GST-Ub1– 75 conjugates were captured on affinity resin and washed extensively to remove soluble ubiquitination components. Residual E1/E2/E3 activity was quenched with 5 mM N-ethylmaleimide for 15 min at room temperature, after which the beads were washed and equilibrated in DUB reaction buffer containing 50 mM HEPES-KOH (pH 7.5), 100 mM NaCl, 1 mM DTT, and 5 mM MgCl2.

### Assembly and deubiquitylation of peptide-anchored ubiquitin chains

Biotinylated peptides were immobilized on streptavidin magnetic beads and washed to remove unbound peptide. Peptide-loaded beads were then subjected to an ATP- and Mg²⁺- dependent in vitro ubiquitination reaction containing E1, an AMFR RING–UBE2G2 fusion E2/E3 reagent described previously^35^, and ubiquitin at 100 µM for 1 h. Following chain assembly, beads were washed extensively to remove soluble ubiquitination components and incubated with 5 mM N-ethylmaleimide (NEM) for 15 min at room temperature to suppress residual E1/E2/E3 activity. Beads were subsequently washed and equilibrated in DUB reaction buffer containing 50 mM HEPES-KOH (pH 7.5), 100 mM NaCl, 2 mM DTT and 5 mM MgCl₂. Peptide-anchored ubiquitin conjugates were incubated with the indicated recombinant DUB at 200 nM for 1 h at 37 °C with shaking. Bead-bound and released fractions were separated and analyzed by SDS-PAGE and anti-ubiquitin immunoblotting. BSA and USP21 served as negative and positive controls, respectively, where indicated.

### Deubiquitylase assays on anchored chains

Resin-bound K48- or K63-directed polyubiquitin chains were incubated with recombinant DeSI1 WT, S25E, or C108S at 200 nM for 1 h at 37 °C with shaking at 900 rpm. BSA was used as a negative control. Where indicated, OTUB1 and AMSH were used as linkage-selective positive controls for K48- and K63-linked chains, respectively. After incubation, supernatant and bead-bound fractions were separated and analysed by SDS-PAGE and anti-ubiquitin immunoblotting. Release of ubiquitin-containing species into the supernatant, together with loss of bead-associated ubiquitin signal, was used as the measure of DeSI1 activity. Time-course reactions were sampled at 0, 30, 60 and 120 min where indicated.

### Peptide binding assays

Fluorescence-polarization peptide-binding assays. Fluorescently labeled peptides corresponding to the COX4 mitochondrial targeting sequence and FIS1 tail-anchored transmembrane domain with its basic C-terminal extension (FIS1 TMD-CTE), together with the indicated hydrophilic control peptide, were analyzed in 384-well black low-volume non-binding plates. Reactions contained 50 nM fluorescent peptide and a threefold serial dilution of recombinant DeSI1 from 15 µM to 0.254 nM, together with a no-protein tracer control, in 25 mM HEPES (pH 7.5), 150 mM NaCl, 0.5 mM TCEP, 0.01% Tween-20 and 0.1 mg/mL BSA. Fluorescence polarization was measured on a BioTek Synergy Neo2 using 485/20-nm excitation, 528/20-nm emission and a 510-nm dichroic. Where indicated, ionic strength was varied by adjusting NaCl to 150, 225 or 300 mM. Binding curves were fitted with a Hill model to obtain apparent K_d_ values and 95% confidence intervals. Technical triplicates were averaged within each experiment, and three independent experiments were performed.

### Peptide-anchored ubiquitin chain assembly and cleavage

Biotinylated peptides corresponding to the COX4 mitochondrial presequence, FIS1 TMD-CTE, or the hydrophilic COX4-derived control were immobilized on streptavidin resin. Polyubiquitin chains were assembled enzymatically on the immobilized peptides, and the resin was washed to remove soluble reaction components before deubiquitylation assays. Peptide-anchored polyubiquitin chains were incubated with BSA, recombinant DeSI1 WT, cyclin D1–CDK4-phosphorylated DeSI1, DeSI1(S25E), DeSI1(C108S), or USP21, as indicated. Following incubation, bead-bound and released material were separated and analysed by SDS-PAGE and immunoblotting with anti-ubiquitin antibodies. Release of ubiquitin-containing species into the supernatant, together with depletion of bead-associated polyubiquitin, was used as the measure of substrate cleavage. For kinetic experiments, COX4- and FIS1-anchored chains were incubated with DeSI1(S25E) and sampled at 0, 30, 60, and 120 min.

### GPS reporter assays

U2OS cells stably expressing Global Protein Stability (GPS) reporters were generated by retroviral transduction. In these bicistronic reporters, EGFP fused to the indicated test sequence was co-expressed with IRES-driven mCherry as an internal expression control. Reporter lines were analysed in five genetic backgrounds: parental cells; parental cells carrying doxycycline-inducible cyclin D1(T286A); *DESI1*-knockout cells; and *DESI1*-knockout cells reconstituted with pBabe HA-DeSI1 WT or S25E.

Cells were analysed under basal conditions, after cycloheximide treatment (100 µg/mL, 4 h), and 120 min after cycloheximide washout. For quiescence–G1 experiments, cells were serum-starved for 48 h and analysed after 6 h of serum re-addition, with or without cycloheximide. For organelle-specific stress experiments, ER reporters were treated with tunicamycin 5 µg/mL and mitochondrial reporters with FCCP 10 uM for 120 min and analyzed at the end of the treatment period and 120 min after washout.

Cells were harvested by trypsinization, washed in PBS and resuspended in FACS buffer containing PBS, 2% FBS and 2 mM EDTA. At least 10,000 live singlet events were acquired per sample. The GFP/mCherry ratio was calculated for each cell, and the mean ratio for each sample was normalized to that of parental cells within the same treatment condition (parental = 1). Three independent experiments were performed per condition.

### AHA metabolic labelling, click chemistry and pulse-chase

Cells were washed twice with methionine-free DMEM and pre-incubated in methionine-free medium for 30 min to deplete intracellular methionine, then pulse-labelled with 4 mM L-azidohomoalanine (AHA; Thermo Fisher Scientific, C10102) for 30 min at 37 °C. For the experiments shown in Figure 5C,D and Supplementary Figure 5E, the chase duration was 90 min. Cells were lysed in denaturing buffer containing 50 mM Tris-HCl pH 8.0, 150 mM NaCl, 1% SDS and 10 mM N-ethylmaleimide (NEM).

Copper(I)-catalysed azide–alkyne cycloaddition was performed with biotin-PEG4-alkyne (100 µM; Click Chemistry Tools, TA105) or AZDye647A-alkyne (Jena Bioscience, CLK-1301A) in the presence of 1 mM CuSO₄, 10 mM sodium ascorbate and 100 µM THPTA for 1 h at room temperature with constant mixing, using the Click-iT Protein Reaction Buffer Kit (Thermo Fisher Scientific, C10276) where indicated. Excess click reagents were removed before enrichment.

Biotinylated nascent proteins were captured on Dynabeads MyOne Streptavidin T1 (Thermo Fisher Scientific, 65601) pre-blocked with 3% BSA, washed stringently and eluted by boiling in 2× Laemmli sample buffer. Whole-cell extracts and streptavidin pull-downs from the same experiments were analysed by anti-biotin immunoblotting to monitor AHA labelling and capture efficiency. The captured fractions were immunoblotted for the indicated nascent ER and mitochondrial proteins and non-hydrophobic controls.

For quantification of nascent-protein retention in Figure 5D, band intensities for MFN2, MCU, OMP25, DHCR7, ANT2 and COX4 were measured in Fiji/ImageJ. For each client, retention was calculated within the same cell line as the chase signal divided by the pulse signal × 100. Statistical analyses were performed on experiment-level means across the six clients; clients were not treated as independent biological replicates. Three independent experiments were analysed

### Extracellular flux analysis

Oxygen-consumption rate (OCR) was measured using a Seahorse XFe24 analyser (Agilent). RPE-hTERT shDeSI1 cells carrying doxycycline-inducible DeSI1 WT or S25E were seeded at 4 × 10⁴ cells per well in XFe24 plates for measurement. Doxycycline treatment was maintained for 24 h for acute experiments or for 7 days for chronic experiments; where indicated, cytidine was present throughout the chronic induction period. On the day of assay, medium was replaced with Seahorse XF DMEM (pH 7.4) supplemented with 10 mM glucose, 1 mM sodium pyruvate, and 2 mM GlutaMAX, followed by 1 h at 37 °C equilibration in a non-CO₂ incubator.

Mito Stress Tests were performed by sequential injection of oligomycin (1 µM), FCCP (1 µM) and rotenone+ antimycin A(0.5 µM each). The FCCP concentration was selected by titration at 0.5, 1 and 2 µM and was fixed at 1 µM for the reported experiments.

Basal mitochondrial OCR was calculated as the final pre-oligomycin OCR minus the minimum OCR after rotenone+antimycin A. ATP-linked OCR was calculated as basal OCR minus post-oligomycin OCR; proton leak as post-oligomycin OCR minus non-mitochondrial OCR; maximal OCR as the maximum FCCP-stimulated OCR minus the minimum OCR after rotenone+antimycin A; spare respiratory capacity as maximal minus basal mitochondrial OCR; and coupling efficiency as ATP-linked OCR divided by basal mitochondrial OCR × 100. Data were exported from Agilent Wave software.

OCR was normalized in each well to the corresponding post-assay cell count and, for absolute traces, expressed as pmol O₂/min per 10,000 cells. Technical wells were averaged within each independent biological experiment and were not treated as independent replicates. Where indicated, endpoint values were expressed relative to the matched shDeSI1 condition from the same biological experiment (shDeSI1 = 100%). Three independent biological experiments were performed per condition.

### Cell cycle analysis

Cells were pulse-labelled with 10 µM EdU for 45 min, washed with PBS, fixed in 4% paraformaldehyde, and processed using the Click-iT Plus EdU Alexa Fluor 488 Flow Cytometry Assay Kit (Thermo Fisher Scientific, C10632) according to the manufacturer’s instructions. Total DNA was counterstained with FxCycle PI/RNase Staining Solution (Thermo Fisher Scientific, F10797). After exclusion of debris and doublets, G1, S and G2/M populations were assigned by bivariate EdU/DNA analysis in FlowJo v10. Three independent biological experiments were performed.

### Transmission electron microscopy

Transmission electron microscopy was performed at the NYU Langone Microscopy Laboratory (RRID:SCR_017934, partially funded by NYU Cancer Center Support Grant NCI P30CA016087). For acute experiments, U2OS parental cells, DESI1-knockout cells, and knockout cells reconstituted with DeSI1 WT or S25E were analysed after 24 h of induction. For chronic experiments, RPE-hTERT shDeSI1 cells expressing empty vector, DeSI1 WT or DeSI1 S25E were analysed after 7 days of induction, with cytidine present throughout the induction period where indicated.

Samples were processed essentially as described previously for electron microscopy performed at the same facility^36^. Cells were fixed overnight at 4 °C in 0.1 M sodium cacodylate buffer (pH 7.2) containing 2.5% glutaraldehyde, 2% paraformaldehyde and 2mM CaCl_2_, post-fixed for 1 h at 4 °C in 1% osmium tetroxide containing 1% potassium ferrocyanide (Sigma), and block-stained overnight at 4 °C with 0.25% aqueous uranyl acetate. Samples were then processed through standard dehydration and embedded in EMbed 812 resin. Ultrathin sections (70 nm) were mounted on 200-mesh copper grids, stained with uranyl acetate and lead citrate, and examined by JEOL 1400 Flash transmission electron microscopy (JEOL, Ltd. Japan) and photographed with a Gatan Rio16 camera (Gatan Inc. Pleasanton, CA). All other chemicals and EM grids are purchased from Electron Microscopy Sciences, Hatfield, PA.

For the acute experiment, mitochondrial ultrastructure was quantified for cristae area fraction, matrix density index, matrix lucency fraction and aspect ratio in four independent biological replicates per genotype. Measurements from individual mitochondrial profiles were summarized within each biological replicate, and the biological-replicate summary was used as the experimental unit.

For the chronic cytidine-rescue experiment, matrix density index, matrix lucency fraction and mitochondrial circularity were quantified in three independent biological replicates per condition. Images used for quantitative analysis were unadjusted 16-bit images; representative fields were displayed after identical linear brightness and contrast adjustment across conditions. Images were acquired at 8,000× (0.83 nm per pixel). Measurements from individual mitochondrial profiles were summarized at the biological-replicate level before statistical analysis.

### Polar metabolomics and lipidomics of cultured RPE cells

#### Experimental design and sample collection

RPE-hTERT shDeSI1 cells and matched shDeSI1 cells carrying doxycycline-inducible DeSI1 WT or S25E were analysed after 72 h of serum starvation or following 8 h of serum-stimulated G1 entry (n = 6 biological replicates per cell line and condition). Doxycycline was added for the final 24 h of the 72-h starvation period, up to serum release, such that these experiments examined the acute DeSI1 state. Replicate identifiers denoted matched experimental blocks across cell lines and conditions. For polar metabolomics, cells were washed twice with ice-cold saline and metabolites were extracted in 80% methanol/water at −80 °C. Insoluble material was removed by centrifugation, and metabolite-containing supernatants were dried and reconstituted for LC-MS analysis.

#### Polar LC-MS

Polar-metabolite profiling was performed at the Laura and Isaac Perlmutter Metabolomics Center, Rappaport Faculty of Medicine, Technion–Israel Institute of Technology. Metabolites were separated using the center’s established ZIC-pHILIC LC-MS workflow, using a ZIC-pHILIC analytical column (150 × 2.1 mm, 5 µm) with a corresponding guard column. The aqueous mobile phase consisted of 20 mM ammonium carbonate containing 0.1% ammonium hydroxide and the organic phase was acetonitrile. Metabolites were separated over a 15-min gradient from 80% organic to 80% aqueous phase at 0.2 mL/min and 45 °C and analysed by high-resolution Orbitrap mass spectrometry with electrospray ionization and polarity switching. Metabolites were identified from accurate mass and retention time using an in-house library established with authentic commercial standards, and peak areas were extracted using Thermo TraceFinder software. This chromatographic and identification workflow has been used consistently in recent studies from the same metabolomics center.

#### Lipidomics

Lipidomic profiling was performed at the same metabolomics center using its established global lipidomics workflow. Lipids were extracted using a methanol:butanol-based extraction and analysed by reversed-phase LC-MS/MS on a Vanquish Flex HPLC system coupled to an Orbitrap Exploris 240 mass spectrometer (Thermo Fisher Scientific). Lipids were separated on an Acclaim C30 column (100 × 2.1 mm, 3 µm) at 60 °C using acetonitrile/water and isopropanol/acetonitrile mobile phases containing 10 mM ammonium formate and 0.1% formic acid. Data were acquired in positive- and negative-ionization modes, and lipid species were identified and aligned using LipidSearch 5.0. Lipid identities and class assignments were taken from the facility-provided annotation table. Free sterols were not quantified in the dataset used for these analyses.

For downstream analysis, metabolite and lipid intensities were log₂-transformed and median-normalized within each sample. G1-versus-starvation responses were calculated separately for each biological replicate and compared across matched experimental blocks. For lipid-class analyses, class abundance was calculated within each sample as the sum of the median-normalized linear abundances of all detected species assigned to that class, followed by log₂ transformation; individual lipid-species rows were analysed using their respective species-level intensities. Lipid identities and class assignments were taken from the facility-provided annotation table. Free sterols were not quantified in the dataset used for these analyses.

### Metabolomics and lipidomics data processing and statistical analysis

#### Preprocessing and quality control

Polar-metabolite and lipidomic datasets were processed separately. Feature intensities were log₂-transformed, with missing or non-positive measurements retained as missing and not imputed. To correct for sample-level loading differences, data were median-normalized within each sample in log₂ space. Sample quality was assessed from the global normalized profiles by principal-component analysis and sample-distance analysis. One shDeSI1 G1 polar-metabolomics sample (replicate 2) was excluded following global-profile quality control. Because subsequent analyses used matched experimental blocks, the corresponding block was excluded from the S25E-versus-shDeSI1 comparison, leaving five matched blocks; all six blocks were retained for the S25E-versus-WT comparison. No lipidomics samples were excluded.

#### Polar-metabolite analysis

For each metabolite, the response to G1 entry was calculated within each cell line and matched biological replicate as the normalized log₂ abundance in G1 minus that in the serum-starved state. The effect of S25E was then calculated within each block as the difference between the S25E response and the corresponding shDeSI1 or WT response. Heatmaps display the mean of these matched difference-in-differences. Statistical significance was assessed using two-sided one-sample *t*-tests against zero, equivalent to paired tests of the genotype-specific G1 responses. Asterisks denote nominal *P* values. Benjamini–Hochberg-adjusted *q* values were additionally calculated within each displayed contrast and are provided in the source data as a multiple-testing sensitivity analysis.

#### Lipid-class and species analysis

Lipid species were not treated as independent biological replicates. For each sample, median-normalized linear abundances of all annotated species belonging to a given lipid class were summed to obtain a single class-level abundance, which was subsequently log₂-transformed. G1-minus-starved responses and S25E-versus-reference difference-in-differences were then calculated using the same matched-block framework as for polar metabolites. The nine prespecified class-level endpoints were phosphatidylglycerol (PG), cardiolipin (CL), phosphatidylinositol (PI), phosphatidylcholine (PC), phosphatidylethanolamine (PE), phosphatidylserine (PS), sphingomyelin (SM), free fatty acids (FA), and triacylglycerol (TG). Nominal two-sided one-sample *t*-test *P* values are indicated in the heatmap, and Benjamini–Hochberg correction was performed separately within each contrast across these nine class-level endpoints. Selected individual lipid species and subclasses, including cardiolipin species and TG unsaturation groups, were secondary analyses and were not included in the primary class-level multiple-testing family. In all analyses, the biological sample was the unit of statistical inference.

### Chemical competition profiling

U2OS cells stably marked with EF1α-GFP (parental cells or DeSI1-knockout cells reconstituted with DeSI1 WT) were mixed 1:1 with EF1α-mCherry-marked cells expressing cyclin D1(T286A) or DeSI1(S25E), respectively. A total of 100,000 cells (50,000 cells of each population) were seeded per well of a 6-well plate and treated for 72 h with the indicated compounds or vehicle (Sham). Relative representation of the two populations was determined by flow cytometry from the GFP:mCherry ratio and normalized to the corresponding Sham condition (Sham = 1.0). Three independent experiments were performed. In all cytidine-containing conditions, cytidine (100 µM) was co-administered with tetrahydrouridine (THU; 100 µM) to inhibit cytidine deamination; uridine was used at 100 µM where indicated. Compounds and final concentrations used in the screen are listed in Supplementary Table 4.

### Proliferation assays

RPE-hTERT shDeSI1 cells expressing doxycycline-inducible DeSI1 WT or S25E were seeded at 10,000 cells per well and induced with doxycycline for 8 days in the presence or absence of the indicated supplements. Cell numbers were determined every 2 days. Population expansion was calculated relative to the number of cells seeded on day 0. For the experiment shown in Figure 6I, cytidine (200 µM) was co-administered with THU (100 µM) throughout the 8-day assay to inhibit cytidine deamination. Three independent experiments were performed.

### Mouse lines

All animal procedures were approved by the NYU Langone Institutional Animal Care and Use Committee (protocol IA16-00005) and performed in accordance with institutional and federal guidelines. Knock-in and knock-out lines were generated on a C57BL/6J background.

#### Generation of *Desi1* mutant mice

The *Desi1* loss-of-function allele was generated by CRISPR–Cas9 editing of mouse embryonic stem (ES) cells. Correctly edited ES-cell clones were validated by sequencing, including TOPO cloning of PCR products, and used in subsequent attempts to derive mice. The loss-of-function allele was generated by editing exon 1 introducing a premature termination codon at amino acid 27 and predicting a 26-amino-acid truncated product. Mice carrying this allele were successfully derived from the edited ES cells and subsequently maintained through the NYU Langone Rodent Genetic Engineering Laboratory. The *Desi1* S25E mouse line was generated by introducing the S25E substitution (AGC→GAG) at the endogenous *Desi1* locus, whereas the S25A line was generated by introducing the S25A substitution (AGC→GCT). Edited embryos were transferred to recipient females, and founder animals carrying the intended substitutions were identified by genotyping and sequence validation and bred to establish the respective lines. In all three cases, heterozygous animals were intercrossed to generate experimental cohorts.

#### Genotyping and animal husbandry

Genotyping was performed on tail or ear biopsies by PCR using the primers 5′-TAAGCGTGCAAGCAAGCCACCA-3′ (forward) and 5′-TTGACCTAACCTGGAGCCGGCA-3′ (reverse), followed by Sanger sequencing of the amplified region to distinguish wild-type, heterozygous and homozygous animals. The Desi1 S25A line was used for Mendelian transmission analysis; because no homozygous Desi1^S25A/S25A^ animals were recovered, subsequent phenotypic and molecular analyses were performed using wild-type, Desi1^S25E/S25E^ and Desi1^−/−^ mice. Mice were maintained under specific pathogen-free conditions on a 12-h light/dark cycle with ad libitum access to standard chow and water. Both sexes were used, as indicated. Mendelian transmission from heterozygous intercrosses was assessed by χ² goodness-of-fit testing against the expected 1:2:1 genotype distribution.

### Behavioral phenotyping

Wild-type (n = 10), Desi1S25E/S25E (n = 11) and Desi1-null (n = 10) mice of both sexes were phenotyped longitudinally at 3, 4 and 5 months of age by an observer blinded to genotype.

#### Neurological phenotype scoring

Neurological function was assessed using the composite phenotype scoring system described by Guyenet et al^37^. Mice were evaluated for performance on the ledge test, hindlimb clasping, gait and kyphosis. Each measure was scored from 0 to 3, with increasing scores indicating greater phenotypic severity, and the four component scores were summed to generate a cerebellar composite score ranging from 0 to 12. The animal was the biological unit of analysis. At each age, composite scores were compared between genotypes using two-sided Mann–Whitney U tests. Longitudinal composite scores within S25E mice were analysed using a Friedman test followed by paired two-sided Wilcoxon signed-rank tests for the 3-versus-4-month and 4-versus-5-month comparisons.

#### Balance-beam testing

Motor coordination was assessed using a balance-beam assay. Mice were trained for 2–3 days before the test day on a 28-mm-round beam, with three training trials per day, until they reliably traversed the beam to the goal box. On the test day, mice were tested on a 10-mm-square beam. Traversal time from the starting position to the goal box was recorded in three trials, and the mean of the three trials was calculated for each animal and used as the biological replicate for statistical analysis. Balance-beam traversal times were compared between genotypes at each age using two-sided Welch’s t-tests. The mean of three trials was used as the value for each animal.

### Tissue collection and histology

At the indicated experimental endpoints, mice were euthanized in accordance with the approved institutional animal protocol. For biochemical and omics analyses, tissues were dissected on ice, snap-frozen in liquid nitrogen and stored at −80 °C until processing. For the tissue-expression panel in Figure 7E, heart, cerebellum, kidney, brain, lung, muscle, spleen, small intestine, stomach, liver, pancreas and large intestine were collected from adult wild-type C57BL/6 mice. Tissue lysates were prepared as described under Immunoblotting. For histological analysis, brains were fixed in 4% paraformaldehyde for 24 h and processed at the Experimental Pathology Research Laboratory, Division of Advanced Research Technologies, NYU Grossman School of Medicine (RRID:SCR_017928). Fixed tissues were dehydrated through graded ethanols, cleared in xylene and infiltrated with Surgipath Paraplast X-tra paraffin (Leica Biosystems, 39603002) using a Leica Peloris III automated tissue processor. Samples were embedded in the desired orientation using a Leica HistoCore Arcadia embedding station, and 5-µm sagittal and coronal sections were prepared on a Leica RM2255 microtome. For cerebellar analyses, midsagittal sections encompassing the vermis were selected consistently across animals.

### Bielschowsky silver staining

Bielschowsky silver staining and brightfield imaging. Five-micrometre paraffin sections were deparaffinized and rehydrated using a Leica HistoCore SPECTRA automated slide-staining workstation and stained with a Bielschowsky Silver Stain Kit (Abcam, ab245877) according to the manufacturer’s instructions. Sections were incubated in 20% silver nitrate solution for 15 min at 40 °C, rinsed in distilled water and incubated in ammoniacal silver solution for 10 min at 40 °C. Sections were then developed for approximately 20 s in a solution containing formalin, citric acid and nitric acid, incubated in ammonia water for 30 s, rinsed in distilled water and treated with 5% sodium thiosulfate for 2 min. Slides were subsequently dehydrated and coverslipped on the HistoCore SPECTRA using HistoCore SPECTRA CV X1 mounting medium (Leica Biosystems, 3801733). Brightfield slides were scanned at 40× using a Leica AT2 whole-slide scanner with Aperio Image Library v12.0.16 (Leica Biosystems), and image files were stored in the NYU Grossman School of Medicine OMERO Plus system.

### Multiplex immunofluorescence and image acquisition

Multiplex immunofluorescence was performed on formalin-fixed, paraffin-embedded brain sections using a Leica BondRx automated stainer according to the manufacturer’s instructions. Sections were deparaffinized and subjected to heat-mediated antigen retrieval with Leica ER1 (AR9961) or ER2 (AR9640) retrieval buffer at 100 °C for 20 min, followed by treatment with 3% H₂O₂ to suppress endogenous peroxidase activity. Sections were blocked with Primary Antibody Diluent (Leica, AR93520) and sequentially incubated with primary antibody and HRP-polymer detection reagents followed by HRP-mediated Opal tyramide signal amplification. After each staining cycle, primary and secondary antibodies were removed by heat-mediated retrieval before application of the subsequent antibody/fluorophore pair. Following completion of the three staining cycles, sections were counterstained with spectral DAPI (Akoya Biosciences, FP1490) and mounted with ProLong Gold Antifade reagent (Thermo Fisher Scientific, P36935).

The three-marker panel consisted of Iba1, calbindin-D28k and GFAP. Iba1 was detected with Wako 019-19471 (1:250; RRID:AB_839504), calbindin with Proteintech 14479-1-AP (1:5,000; RRID:AB_2228318), and GFAP with Abcam ab7260 (1:1,700; RRID:AB_305808). Rabbit-on-Rodent HRP polymer was used for detection, with Opal 620, 570 and 520 assigned to Iba1, calbindin and GFAP, respectively.

Whole-slide multispectral imaging was performed using a PhenoImager HT system (formerly Vectra Polaris; Akoya Biosciences). Slides were initially scanned at 20× magnification to establish exposure settings. A spectral-unmixing library was generated using PhenoImager HT software v2.0.0 and inForm v3.0, and this library was used to create an unmixed acquisition protocol. Slides were subsequently rescanned at 20× to generate spectrally unmixed whole-slide qptiff files, which were stored in the NYU Grossman School of Medicine OMERO Plus image-management system (Glencoe Software).

GFAP-positive area was quantified in QuPath using a pixel classifier on coded images and summarized to per-animal means using the same hierarchical approach.

### Quantitative cerebellar histology

Purkinje-cell density was quantified from calbindin-positive midsagittal cerebellar sections in QuPath as the number of calbindin-positive Purkinje-cell somata per millimetre of traced Purkinje-cell layer. Images were analysed using coded identifiers and decoded after scoring. Measurements from individual regions were first collapsed to section-level values and subsequently to per-animal means, with the animal used as the unit of statistical analysis. Molecular-layer thickness was measured from the same sections as the mean length of perpendicular measurements spanning the molecular layer and similarly summarized at the animal level.

### Brain proteomics

Whole brains from wild-type, DeSI1(S25E/S25E) and DeSI1-null mice (n = 6 per genotype, with balanced representation of males and females) were lysed in RIPA buffer supplemented with 5 mM EDTA, cOmplete EDTA-free protease inhibitor cocktail (Roche) and phosphatase inhibitors, and cleared by centrifugation. Proteomic analysis was performed by the NYU Langone Proteomics Laboratory (RRID:SCR_017926; project PRL-SK-1867).

Cleared lysates were reduced with DTT at 57 °C for 1 h, alkylated with iodoacetamide at room temperature in the dark, and processed by single-pot solid-phase-enhanced sample preparation (SP3). Proteins were bound to SP3 beads in ethanol, washed four times with 80% ethanol, and digested overnight with trypsin in ammonium bicarbonate. Peptide digests were acidified with trifluoroacetic acid and loaded onto Evotips together with indexed retention-time (iRT) standards. Peptides were separated using an Evosep Eno LC system and analysed in dia-PASEF mode on a Bruker timsTOF mass spectrometer.

Data were processed in Spectronaut v20.5.260227.92449 against the mouse UniProt database using dynamic mass tolerances. Carbamidomethylation of cysteine was specified as a fixed modification, and protein N-terminal acetylation, GlyGly modification of lysine, methionine oxidation and phosphorylation of serine, threonine and tyrosine were specified as variable modifications. Identifications were controlled at 1% false-discovery rate. Protein quantification was based on MS2 areas without additional normalization. Proteins supported by only a single unique peptide were excluded from quantitative analyses.

Differential brain-proteome analysis. Protein-level quantities from the Spectronaut Protein_Quant output were log2-transformed and median-normalized within each sample. For each pairwise genotype comparison, proteins quantified in at least two biological replicates in each group were retained. Log2 fold change was calculated as the difference between the mean normalized log2 protein abundance in the two groups. Statistical significance was assessed using two-sided Welch’s t-tests, and P values were adjusted separately within each genotype contrast using the Benjamini–Hochberg procedure. Proteins with q < 0.05 and |log2 fold change| ≥ 0.5 were considered significantly altered.

For the focused nucleotide-pathway analysis in Supplementary Figure 9F, protein abundances were compared between genotypes using two-sided Welch’s t-tests on log₂-transformed protein quantities without correction for multiple comparisons.

Whole-brain immunoblot signals were normalized to the indicated loading control and expressed relative to the wild-type mean. Where proteins were quantified across two membranes, measurements represented n = 6 independent mice per genotype. Pairwise comparisons of S25E with wild-type and DeSI1-null mice were performed using two-sided Welch’s t-tests.

### Brain lipidomics and polar metabolomics

Polar metabolite and lipid profiling was performed by the NYU Langone Metabolomics Core Resource Laboratory (RRID:SCR_017935; iLab service ID MBX-SK-1669) on one brain hemisphere per animal from the same 6-month cohort used for proteomics (initially n = 6 per genotype, with equal male and female representation). Polar metabolites were acquired using the laboratory’s hybrid polar LC-MS assay (HELM:0.92.12.12), and lipids using its global nonpolar lipidomics assay extended to the lower mass range for free fatty acids (HELM:0.92.10.22). Brain_SE_1 was not acquired in the lipidomics or precursor-ion runs.

#### Sample-level quality control

Global sample QC was performed before pathway-level statistical testing using principal-component analysis of log2-transformed profiles together with within-genotype sample-to-sample correlations on each platform. Brain_WT_3 was excluded from the primary brain metabolomic and lipidomic analyses because it was the only sample showing concordant outlier behavior in both the polar metabolome and lipidome. Samples flagged on only one platform were retained in the primary analysis. The resulting primary cohorts were WT n = 5, S25E n = 6 and KO n = 6 for polar metabolomics, and WT n = 5, S25E n = 5 and KO n = 6 for lipidomics. Analyses combining lipid and polar-metabolite measurements were restricted to the common S25E and WT cohort (n = 5 per genotype). Sensitivity analyses with Brain_WT_3 retained were performed for the principal pathway-balance endpoints and are provided in the source data.

#### Extraction

Frozen brain tissue was homogenized in 100% methanol to a tissue-equivalent concentration of 20 mg/mL using a two-layer extraction based on the Vorkas et al^38^. tissue-metabolomics workflow. After phase separation with water and chloroform, 100 µL of the upper aqueous-methanol phase was collected for polar metabolomics, dried in a speed vacuum and reconstituted in 20 µL LC-MS-grade water. For lipidomics, equal aliquots of the upper and lower phases were collected, dried and reconstituted in 40 µL isopropanol:acetonitrile:water (4:3:1, v/v/v). Isotopically labeled internal standards added during extraction were used to monitor analytical performance.

#### Polar LC-MS

Polar metabolites were separated on a ZIC-pHILIC column (2.1 × 150 mm, 5 µm; Millipore/SeQuant) coupled to a Dionex Ultimate 3000 system, with the column held at 25 °C. Mobile phase A was 10 mM ammonium carbonate in water (pH 9.0) and mobile phase B was acetonitrile, at 0.1 mL/min. The gradient was 80% to 20% B over 0-30 min, returned to 80% B from 30-31 min and held at 80% B to 42 min; injection volume was 2 µL. MS analysis used a Thermo Q Exactive HF mass spectrometer with heated electrospray ionization and polarity switching in a data-dependent Top 5 method. Spray voltage was 3.5 kV, capillary temperature 320 °C, sheath gas 35, auxiliary gas 10 and maximum spray current 100 µA. Full MS scans were acquired at 120,000 resolution (AGC target 3 × 10^6, maximum injection time 100 ms, m/z 67-1000). Tandem MS scans were acquired at 15,000 resolution (AGC target 1 × 10^5, maximum injection time 50 ms, isolation window 0.4 m/z, isolation offset 0.1 m/z, fixed first mass 50 m/z) using stepped normalized collision energies of 10, 35 and 80. The minimum AGC target was 1 × 10^4 with an intensity threshold of 2 × 10^5. Data were acquired in profile mode.

#### Lipid LC-MS

Lipid extracts were separated on an Acquity Premier CSH C18 column (2.1 × 150 mm, 1.7 µm, 130 Å; Waters, part no. 186005298). Mobile phase A was acetonitrile:water (60:40, v/v) containing 10 mM ammonium acetate and 0.1% acetic acid, and mobile phase B was isopropanol:acetonitrile (90:10, v/v) containing 10 mM ammonium acetate and 0.1% acetic acid. The gradient was 40% B from 0–1.25 min, 40–50% B from 1.25–2 min, 50–54% B from 2–11 min, 54–70% B from 11–12 min, 70–99% B from 12–18 min, held at 99% B from 18–23 min, and returned to 40% B from 23–24 min. The injection volume was 2 µL.

#### Lipid identification and feature curation

Data-dependent lipid MS/MS spectra were searched against the LipidBlast tandem mass spectral library using an in-house implementation of MSPepSearch_x64. Putative identifications were ranked by reverse-dot score; duplicate structural assignments were removed by retaining the top-scoring spectrum together with its neutral formula, detected m/z and ionization polarity. For downstream statistical analysis, unannotated features and isotopic-adduct entries were removed. When multiple annotated ions mapped to the same core-provided cluster, one representative ion was retained by selecting the most abundant non-isotopic annotated feature across the dataset.

#### Analytical QC and normalization

Instrument performance was monitored with the isotopically labeled internal-standard mixture and core QC injections. For the brain polar dataset (SQ2193), reported mass accuracy was 1.7 ppm, retention-time drift across the run was 0.31 min, and median internal-standard response variability was 16% across biological samples and 4% across internal-standard injections. For downstream analysis, positive polar-metabolite intensities and lipid intensities were log2 transformed. Each platform was median-centered within sample to remove multiplicative sample-loading differences while preserving between-feature relationships; lipid values were converted back to linear space after normalization when class totals were required. Features with fewer than three finite positive values in either group were not assigned a P value.

#### Polar-metabolite comparisons

For the supplementary polar-metabolite heatmap, S25E/WT and S25E/KO effects were calculated as differences in mean median-normalized log2 abundance, with the mouse as the unit of replication. Between-genotype comparisons used two-sided Welch t-tests. Heatmap asterisks denote nominal P values; Benjamini-Hochberg (BH) q values were calculated across all displayed metabolites and separately within the predefined pathway blocks and are reported in the source data.

#### Mouse-level lipid-class analysis

Lipid species were not treated as biological replicates. For each mouse, a single abundance value for each lipid class was calculated by summing the per-sample median-normalized linear abundances of all de-duplicated annotated species assigned to that class and then log2 transforming the class total. S25E/WT and S25E/KO differences in class abundance were tested by two-sided Welch t-tests across mice. The primary multiple-testing family comprised nine predefined lipid classes (PA, TG, PG, PI, PC, PE, PS, SM and CL), with BH correction performed separately for each genotype contrast. Cardiolipin was additionally summarized by total double-bond number (≤4, 5-8 and ≥9); these saturation bins and individual lipid-species analyses were treated as secondary analyses and were not included in the primary class-level BH family. Nominal P-value stars are displayed in the heatmap, with BH q values reported in the figure legend and source data.

#### Phospholipid pathway-balance analysis

The main brain lipidomics panel was designed as a within-mouse pathway-balance analysis so that each animal, rather than each lipid species, remained the independent observation. Log2 ratios were calculated within each mouse for PA/TG, PG/PA, PI/PA and CL/PG. Aggregate branch balances were defined on the log scale so that constituent classes contributed equally: the CDP-DAG branch was the geometric mean of PI, PG and CL, and the mitochondrial PG+CL branch was the geometric mean of PG and CL. These branch summaries were expressed relative to PA or TG as indicated in the figure. S25E/WT differences in within-mouse log2 balances were tested by two-sided Welch t-tests. Asterisks in the heatmap denote nominal P values. BH q values were calculated across all displayed rows and, separately, across the prespecified CDP-DAG mechanistic family (PG/PA, PI/PA, CDP-DAG branch/PA, [PG+CL] branch/TG and CDP-DAG branch/TG) and are provided in the source-data table.

#### Kennedy-pathway and cytidine:uridine specificity controls

The re-derived polar-metabolite dataset included CDP-choline and CDP-ethanolamine. Within-mouse log2 ratios of CDP-choline/phosphocholine, CDP-ethanolamine/phosphoethanolamine, CMP/UMP and cytidine/uridine were calculated directly from the corresponding positive intensities and tested between S25E and WT mice by two-sided Welch t-tests. To match the lipidomics cohort used in the pathway-balance panel, these cross-platform specificity controls were restricted to the common S25E and WT samples (n = 5 per genotype). These rows were included as mechanistic specificity controls rather than as evidence of equivalence when nonsignificant.

#### CDP-choline precursor-ion validation

A separately re-derived precursor-ion dataset was used as an orthogonal check of CDP-choline-related signals. The precursor at m/z 489.1148 yielding a product ion at approximately m/z 360.0616 was treated as the CDP-choline parent-like feature. Higher-m/z precursor features yielding the same approximately 360 product ion were reported by their measured precursor and product m/z and by delta mass relative to 489.1148; no structural assignments were made from delta mass alone. S25E/WT and S25E/KO effects were tested by two-sided Welch t-tests on log2 intensities, with BH correction across the precursor-ion scan within each contrast. Agreement between the targeted CDP-choline measurement and the 489.1148 precursor feature was evaluated by Pearson correlation on the common quantified samples.

### Pathway and enrichment analysis

Gene Ontology over-representation analysis of the DeSI1 interactome was performed using Enrichr^39^. Proteins quantified in the S25E-versus-WT interactome were ranked by log₂ fold change, and the 100 most directionally enriched proteins from each end of the distribution were analysed separately, with all proteins tied at the rank-100 boundary retained. This yielded 101 S25E-enriched and 135 WT-enriched proteins. Gene Ontology Biological Process and Cellular Component annotations were used to identify the dominant functional themes. Full input lists and enrichment statistics are provided in the source data. The protein groups displayed in Supplementary Figure 3B–C are representative members of the resulting functional themes and are not restricted to individual statistically significant GO terms.

For the targeted membrane-compartment analysis of the His-ubiquitin screen, the 4,531 proteins quantified in the common analysis universe were ranked by the S25E-versus-EV ubiquitylation statistic. The 907 proteins in the lowest 20% of this distribution, representing proteins with the strongest reduction in His-Ub recovery in S25E-expressing cells, were tested for enrichment of six prespecified membrane-associated Cellular Component categories: outer membrane, ER membrane, ER exit site, coated-vesicle membrane, vesicle membrane and Golgi membrane. Enrichment was calculated relative to the complete 4,531-protein quantified background using one-sided Fisher’s exact tests, with Benjamini– Hochberg correction across the six prespecified categories.

For the integrated transcriptome–proteome analysis in Figure 3D, gene-set enrichment analysis was performed independently on the RNA-seq and proteomic differential-response rankings using the MSigDB C5 Gene Ontology Biological Process and Cellular Component collections (human gene-symbol sets, v2026.1). Normalized enrichment scores (NES) were used to compare the magnitude and direction of pathway responses at the RNA and protein levels. Positive NES values indicate enrichment under CDK4/6-active conditions relative to palbociclib treatment. Figure 3D displays the RNA and protein NES for all tested terms and highlights selected protein-dominant, transcript-dominant and contextual pathways.

### Statistical analysis

Statistical analyses were performed using Python 3.10+ (including pandas, NumPy, SciPy, statsmodels and Matplotlib) and GraphPad Prism 10. The biological replicate was the unit of statistical inference unless otherwise stated. Technical replicates, including Seahorse wells and measurements from individual mitochondrial profiles or histological regions, were summarized within the corresponding biological replicate before statistical testing. Sample sizes and the statistical test used for each experiment are specified in the corresponding figure legend.

Unless otherwise indicated, comparisons between two independent groups were performed using two-sided Welch’s t-tests. Matched experimental designs were analysed using paired tests or repeated-measures models as indicated. Factorial experiments were analysed using two-factor models including the relevant interaction term, and genotype × treatment or genotype × condition interaction P values are reported where applicable. Non-parametric tests were used where specified in the corresponding figure legends.

Multiple-testing correction was applied to prespecified families where indicated. Phosphoproteomic datasets were controlled using Benjamini–Hochberg false-discovery rates within each screen. GPS reporter comparisons were corrected using the Holm procedure, and chemical-competition analyses used Holm–Šídák correction within the specified comparison families. For cellular and brain lipidomics, Benjamini–Hochberg correction was applied to the nine prespecified lipid-class endpoints within each contrast, whereas individual lipid species and secondary compositional analyses were analysed separately. Targeted membrane-compartment enrichment in the His-ubiquitin screen was corrected across the six prespecified categories. Proteome-wide brain DIA-MS significance was defined using Benjamini–Hochberg-adjusted Q values. Where both nominal P values and adjusted values are reported, the relevant convention is stated in the figure legend and source-data table.

Analyses reported only with uncorrected nominal P values were treated as exploratory or descriptive unless an omnibus or interaction test was specified as the primary inferential test; conclusions were not based solely on isolated nominally significant pairwise comparisons

Mendelian transmission was assessed by χ² goodness-of-fit testing against the expected 1:2:1 genotype distribution. Longitudinal within-animal behavioural comparisons were analysed using paired non-parametric tests as specified in the corresponding legend. Bar graphs show mean ± SEM unless otherwise indicated, with individual biological replicates displayed where possible. Where asterisks denote nominal P values, *P < 0.05, **P < 0.01, ***P < 0.001 and ****P < 0.0001; panels displaying adjusted q values are explicitly identified as such.

## DATA AND CODE AVAILABILITY

Data deposition is in progress. Raw and processed mass-spectrometry proteomics data generated in this study will be deposited with the ProteomeXchange Consortium through MassIVE, and RNA-sequencing data will be deposited in the NCBI Gene Expression Omnibus. Metabolomics and lipidomics datasets will be deposited in public data repositories, including Zenodo. Uncropped immunoblot images and analysis scripts used to generate the principal computational analyses and figures will also be made publicly available. Repository accession numbers, DOIs, and links will be provided upon completion of deposition and no later than publication. Source data underlying the figures, including quantitative values and statistical analyses, will accompany the final manuscript. Additional information required to reproduce or reanalyze the data reported in this study is available from the lead contact upon reasonable request.

## ACKNOWLEDGMENT

This work was supported in part by NIH/NIGMS grant R35-GM136250 to M.P. M.P. and N.Z. are Investigators of the Howard Hughes Medical Institute. S.K. is supported by an NIH/NIGMS K99 Career Development Award (1K99GM155613-01A1) and a Weizmann Institute Career Development Award and was previously supported by a Life Sciences Research Foundation fellowship and an EMBO Long-Term Postdoctoral Fellowship. We thank the members of the Experimental Pathology Research Laboratory (RRID:SCR_017928), Proteomics Laboratory (RRID:SCR_017926), Microscopy Laboratory (RRID:SCR_017934), Advanced Rodent Transgenics Laboratory (RRID:SCR_017692), and Metabolomics Laboratory (RRID:SCR_017935) at NYU Langone Health for technical and infrastructure support. The Advanced Rodent Transgenics Laboratory also assisted with generation of the *Desi1* mutant mouse lines. These cores are partially supported by Cancer Center Support Grant P30CA016087 to NYU Langone’s Laura and Isaac Perlmutter Cancer Center. The PhenoImager HT multispectral imaging system was initially purchased through Shared Instrumentation Grant S10 OD021747. We thank the Genome Technology Center at NYU Langone Health (RRID:SCR_017929) for RNA-sequencing library preparation and sequencing. The Genome Technology Center is partially supported by Cancer Center Support Grant P30CA016087 to NYU Langone’s Laura and Isaac Perlmutter Cancer Center.

## AUTHORS CONTRIBUTIONS

S.K. and M.P. conceived the study. S.K. designed and coordinated the experimental program, performed experiments and data analyses, and supervised the study. E.L., R.W., J.E., M.C., D.S., Y.-T.J., M.L., Q.Z., and Y.K. performed experiments and/or provided technical assistance. I.A. and H.W. performed and analyzed the RPE polar-metabolomic and lipidomic studies. T.R. and D.J. performed and analyzed the brain lipidomic studies. G.W. and C.A.L. performed histological and immunohistochemical processing and analyses. R.B. contributed to generation of the mouse strains, including tetraploid-complementation procedures. A.H. contributed to RNA-seq data generation. N.D.U. and S.A.C. generated and analyzed one of the HCT-116 mAID proteomic and phosphoproteomic screens (screen 1). J.Y. and F.-X.L. performed transmission electron microscopy. T.C. and B.U. generated and analyzed mass-spectrometry data for the DeSI1 interactome, His-ubiquitin proteomic screen, one of the HCT-116 mAID proteomic and phosphoproteomic (screen 2), and brain proteomics. N.Z. contributed to structural modeling and interpretation. S.K. and M.P. wrote the manuscript with input from all authors.

## Supplementary Figure Legends

**Supplementary Figure 1: Phosphoproteomic screen identifies DeSI1 as a major cyclin D1-CDK4 substrate (A)** Global proteome corresponding to phosphoproteomic Screen 1. Volcano plot shows protein-abundance changes for the same comparison used in Figure 1B. DeSI1 and representative RB-family proteins are indicated. Dashed line denotes FDR = 0.05.

**(B)** Global proteome corresponding to phosphoproteomic Screen 2. Volcano plot shows protein-abundance changes for the same comparison used in Figure 1C. Representative RB-family proteins are indicated. Dashed line denotes FDR = 0.05.

**(C)** Comparison of global-proteome and phosphoproteome significance for the top phosphopeptide hits in Screen 1 (left) and Screen 2 (right). Heatmaps show −log10(P) values in the corresponding global-proteome and phosphoproteome datasets; phosphopeptide sequences are shown at right with the phosphorylated residue highlighted. DeSI1 p-S25 ranks among the strongest phosphorylation changes despite little or no corresponding change in total protein abundance.

**(D)** Specificity controls. HEK293T cells were transfected with non-targeting (NT) or DeSI1 siRNA and the indicated expression vectors. Phos-tag analysis detects phosphorylation of exogenous HA-DeSI1; conventional SDS-PAGE shows endogenous and exogenous DeSI1 and the indicated controls.

**(E)** The p-DeSI1(S25) antibody recognizes the phosphomimetic S25E substitution. Upper, HEK293T DeSI1-knockout cells expressing Flag-DeSI1 WT, S25A (SA), or S25E (SE); whole-cell extracts (WCE) and anti-Flag immunoprecipitates were immunoblotted with the indicated antibodies. Asterisk indicates a non-specific band. Lower, recombinant Flag-DeSI1 WT, S25A (SA), C108S (CS), and S25E (SE) purified from *E. coli* were immunoblotted with the indicated antibodies.

**(F)** Cyclin–CDK specificity of DeSI1 S25 phosphorylation. HEK293T DeSI1-knockout cells expressing HA-DeSI1(WT) were co-transfected with the indicated FLAG-tagged CDKs and EGFP-tagged cyclins. DeSI1 S25 phosphorylation was detected with the phospho-specific p-DeSI1(S25) antibody. Expression and activity of the indicated cyclin–CDK complexes were monitored by FLAG, GFP, p-RB(S807/811), and p-H3(S10) immunoblotting; total RB and actin served as controls.

**(G)** Cyclin D1–CDK4 directly phosphorylates DeSI1 at S25 in vitro. Left, purified DeSI1 was incubated with recombinant cyclin D1–CDK4 at the indicated concentrations; where indicated, palbociclib was included. Reactions were immunoblotted for p-DeSI1(S25) and total DeSI1. Right, GST-RB was phosphorylated in parallel with the same kinase preparation as a positive-control substrate.

**Supplementary Figure 2. Dimer-interface determinants and modifier specificity of DeSI1. (A)** Detail of the three interface clusters shown in Figure 2D, with side chains labelled.

**(B)** Size-exclusion chromatography of recombinant interface mutants. LL20,24EE and L144R elute as monomers, F140S gives a heterogeneous profile, and Δ151–163 remains dimeric. Coomassie stain of fractions below.

**(C)** Activity-based probe labelling with HA-Ub-VS. Only S25E forms an adduct; USP7 is the positive control. Upper, probe blot; lower, silver stain.

**(D–G)** Activity-based probe labelling of the indicated DeSI1 variants with HA-SUMO1-VS (D) and biotin-SUMO2-VME (E), cleavage of SUMO2 polychains (F), and labelling with Flag-NEDD8-VS (G). SENP1, SENP2 and SENP8 are positive controls. Upper, probe blot; lower, silver stain.

**(H)** Analytical size-exclusion chromatography of the crosslinking reactions in Figure 2F. Upper panel, immunoblots of fractions under reducing (+BME) and non-reducing (−BME) conditions, with corresponding chromatograms below. Left, S25E monomer; right, wild-type dimer. Lower panel, densitometric quantification of the reducing (+BME) gels shown above. Band intensities for DeSI1 (∼20 kDa) and ubiquitin (∼10 kDa) were integrated per lane after local background subtraction and plotted against elution volume; each trace is scaled to its own maximum, so the two readouts are comparable in shape but not in absolute intensity. All preparations are FLAG-6His-tagged, which shifts every species to earlier elution volumes than the 6His-only proteins in Figure 2A. Dashed vertical lines mark reference elution volumes determined by size-exclusion chromatography of each component run alone: DeSI1 dimer (∼36 kDa) at 8.5 mL, DeSI1(S25E) monomer (∼18 kDa) at 11.0 mL, and free ubiquitin (∼10 kDa) at 12.5 mL. The DeSI1–Ub conjugate (∼30 kDa) position at 9.75 mL was taken from these data, as the volume at which the DeSI1 and ubiquitin signals are jointly maximal in the +BME profile.

**(I)** Time course of K48-linked chain disassembly by wild-type and S25E DeSI1 over 120 min.

**(J)** As in Figure 2G, with K63-linked chains and AMSH as a K63-specific positive control.

**Supplementary Figure 3. Orthogonal proteomic analyses converge on hydrophobic-protein regulation by phospho-DeSI1. (A)** Experimental design of the three orthogonal screens used in Figure 3. Left, phosphorylation-dependent interactome: HEK293T cells expressing FLAG–DeSI1 WT or S25E were subjected to anti-FLAG immunoprecipitation followed by LC–MS/MS. Middle, ubiquitylome: HEK293F cells expressing 6×His–ubiquitin together with the indicated DeSI1 constructs were subjected to Ni-NTA pulldown under denaturing conditions followed by LC– MS/MS. Right, CDK4-dependent proteome–transcriptome analysis: endogenous cyclin D-CDK4/6 activity was acutely increased by AMBRA1 degradation and compared with palbociclib-inhibited cells by quantitative proteomics and RNA-seq.

**(B-C)** Interactors of FLAG-DeSI1 identified by IP-MS in HEK293T cells (Figure 3A), preferentially associated with wild-type DeSI1 relative to S25E (B) or with the S25E phosphomimetic relative to wild type (C). For Gene Ontology over-representation analysis, proteins were ranked by log2 fold change using log₂ fold change. The 100 most enriched proteins in each direction were selected, with all proteins tied at the rank-100 cutoff retained (S25E-enriched, n=101; WT-enriched, n=135). The tables provide a compact visualization of representative, directionally concordant proteins from the full interactome assigned to the dominant functional themes identified by the enrichment analysis; the displayed protein lists are therefore not restricted to the GO input set, and the subheadings are organizational labels rather than a claim that every individual subheading independently meets BH-adjusted P<0.05. Full ranked inputs, GO statistics, and displayed-protein source values are provided in the source data. (B) WT-enriched interactors organize around ubiquitin-dependent protein catabolism/proteostasis, cellular stress responses, and vesicle/protein trafficking. (C) S25E-enriched interactors organize around mitochondrial, ER, vesicle, and other organelle-membrane compartments.

**(D)** Targeted membrane-component enrichment in the S25E deubiquitylation tail. Proteins in the lowest 20% of the common-universe S25E-versus-EV His-Ub ranking (907 of 4,531 proteins) were tested for enrichment of six prespecified membrane-associated cellular-component terms. Values in parentheses indicate the number of lowest-20% proteins/background proteins within each category. Fold enrichment and one-sided Fisher exact P values are shown; Benjamini–Hochberg correction was performed across the six targeted hypotheses. All six categories—outer membrane, ER membrane, ER exit site, coated-vesicle membrane, vesicle membrane, and Golgi membrane—were significantly enriched after correction. The union comprises 193 unique proteins.

**(E)** Proteins within the lowest 20% of the S25E-versus-EV His-Ub ranking were filtered for directional rescue by the catalytically inactive S25E/C108S mutant, defined permissively as t(S25E/C108S versus S25E) > 0. The resulting 165 unique proteins were grouped according to the S25E-interactome functional modules: mitochondrial matrix, mitochondrial membrane, endoplasmic reticulum, vesicle membrane, and bounding membrane of organelle. To avoid duplicate display of proteins assigned to multiple modules, each protein was shown once using the priority order: mitochondrial matrix > mitochondrial membrane > endoplasmic reticulum > vesicle membrane > bounding membrane of organelle.

**(F)** Ubiquitylome proteins preferentially depleted of His-Ub in S25E cells are biased toward membrane-insertion-prone sequences. Frequency distributions of minimum-window ΔG_app for the S25E-associated reduced-ubiquitylation tail and the oppositely ranked EV-enriched population. Lower ΔG_app values indicate greater membrane-insertion propensity. The population with reduced His-Ub recovery in DeSI1(S25E)-expressing cells is shifted toward lower ΔG_app, and the difference remains significant after matching for protein length.

**(G)** CDK4-dependent protein-dominant responses are enriched for membrane-insertion-prone proteins. Cumulative distributions of minimum-window ΔG_app for the top 10% of proteins showing the strongest protein-dominant response to CDK4 activity and the remaining 90% of quantified proteins. The top decile shows lower ΔG_app values (median 0.302 versus 0.380 kcal mol⁻¹; Mann–Whitney P = 4.9 × 10⁻⁵; length-matched P = 0.010), indicating greater membrane-insertion propensity.

**Supplementary Figure 4. Validation and specificity controls for DeSI1 substrate recognition and activity. (A)** Interactome partition by immunoblot. HEK293T cells expressing Flag-tagged EV, WT, phospho-deficient S25A, or phosphomimetic S25E DeSI1 were subjected to anti-Flag immunoprecipitation. WCE and immunoprecipitates were probed for UPS and chaperone components (VCP, ubiquilin 1/2, BAG6, and ubiquitin) and membrane-associated clients (DHCR7, SLC25A5, ATP1A1, TIM23, MFN1, and COX4). SKP1, negative control.

**(B)** Cartoon representation of the experimentally determined human DeSI1 structure (PDB 3EBQ), highlighting residues at and adjacent to the dimer interface. The hydrophobic dimer-interface residues L20, L24, F140, and L144 are shown in orange, the acidic residues E34 and E134 in red, and the regulatory S25 residue in magenta. S25 is unmodified in the experimental structure; phosphorylation is not modeled. The highlighted residues illustrate the juxtaposition of hydrophobic and acidic features surrounding the S25-containing region.

**(C)** Fluorescence-polarization analysis of DeSI1-client binding specificity and ionic-strength dependence. Left, direct binding of DeSI1 to the COX4 mitochondrial presequence measured by fluorescence polarization. Fluorescently labelled COX4 presequence peptide was titrated with recombinant DeSI1 WT, S25E or S25E/C108S in 150 mM NaCl. Points, mean ± s.d. of three biological replicates; curves, Hill fits. Fitted Kᴅ values (µM) are inset; peptide sequences, physicochemical parameters and 95% confidence intervals are given in Supplementary Table 3. Middle, the polar COX4-scaffold control peptide titrated with the same DeSI1 variants; ΔmP is shown tracer-referenced, with the dashed line marking the polarization change obtained with the cognate COX4 presequence for scale. No variant gives a saturable response. Right, binding of S25E/C108S to the COX4 presequence at 150, 225 and 300 mM NaCl; points, mean ± s.d. of three biological replicates; curves, Hill fits; fitted Kᴅ values are inset. Affinity decreases monotonically with increasing ionic strength, consistent with an electrostatic contribution to client binding.

**(D)** Deubiquitylation of COX4-peptide-anchored polyubiquitin chains. Polyubiquitin chains were assembled on streptavidin-immobilized COX4 presequence peptide and incubated with BSA, WT DeSI1, phosphorylated DeSI1 (pDeSI1), S25E, C108S or USP21. Bead-bound and supernatant fractions were resolved by SDS-PAGE and immunoblotted with anti-ubiquitin.

**(E)** Time course of peptide-anchored chain cleavage. COX4- or FIS1-anchored polyubiquitin chains were incubated with DeSI1(S25E) for 0, 30, 60 and 120 min. Bead-bound and supernatant fractions were immunoblotted with anti-ubiquitin.

**(F)** Sequence specificity of peptide-anchored chain cleavage. Polyubiquitin chains were assembled on a streptavidin-immobilized hydrophilic COX4-derived peptide and incubated with BSA, WT DeSI1, phosphorylated DeSI1, S25E, C108S or USP21. Bead-bound and supernatant fractions were immunoblotted with anti-ubiquitin. No DeSI1-dependent chain release is detected.

**(G)** DeSI1 phosphorylation is required for substrate accumulation during G1 entry. RPE-hTERT cells stably depleted of endogenous DeSI1 (shDeSI1) and reconstituted with doxycycline-inducible Flag-tagged S25A or S25E DeSI1 were serum-starved, induced with doxycycline 48 h before release, and released into G1 by serum addition for the indicated times. Whole-cell extracts were immunoblotted for the indicated proteins. Cyclin E1, cyclin A2 and RB phosphorylation monitor comparable cell-cycle progression between lines; Flag confirms transgene expression; actin serves as loading control.

**Supplementary Figure 5. Extended analysis of nascent hydrophobic protein stabilization by the DeSI1 switch. (A)** Schematic of the Global Protein Stability (GPS) reporter. A bicistronic construct expresses EGFP fused to a TMD-containing sequence (GOI–TMD) together with an IRES-driven mCherry internal control, enabling ratiometric measurement of reporter stability.

**(B)** GPS reporter stability across the quiescence–G1 transition. U2OS lines as in Figure 5A were serum-starved (48 h), released by serum re-addition (6 h), or released in the presence of cycloheximide. Heatmaps and statistics as in Figure 5A; n = 3 independent experiments.

**(C)** GPS reporter stability during and after organelle-specific stress. U2OS lines as in Figure 5A were treated for 2 h with tunicamycin (ER stress; DHCR7 and HMGCR, with CA2 and GPS controls; Left) or FCCP (mitochondrial stress; MFN2 and SLC25A5; Right), and reporter stability was analysed at the end of the incubation with these inhibitors, or following a 120-min washout period. The tunicamycin and FCCP datasets are shown as separate heatmaps because the reporter classes were challenged with distinct organelle-specific perturbations. Heatmaps show mean GFP/RFP ratios normalized to parental cells within each condition (parental = 1). Asterisks indicate significantly increased stability relative to parental within the same condition; pooled-residual two-sided *t*-tests with Holm correction. *n* = 3 independent experiments.

**(D)** Total nascent protein capture controls for Figure 5C. Biotin immunoblots of whole-cell extracts and streptavidin pull-downs from the same experiment, confirming comparable AHA labelling and capture efficiency across genotypes and chase points.

**(E)** RB inactivation does not substitute for CDK4-dependent DeSI1 phosphorylation. RPE-hTERT cells expressing doxycycline-inducible cyclin D1 T286A or HPV E7 were subjected to AHA pulse-chase during G1 as in Figure 5C. Left, biotin immunoblot of nascent protein; right, immunoblots of representative clients from distinct compartments (MFN2, MFN1, ANT2, COX4, MCU, ATP1A1, HMGCR, DHCR7, DIABLO) and controls (Rac1, tubulin). cyclin D1, pDeSI1 (S25), HA (E7) and pRB (S807/810) confirm that E7 inactivates RB without inducing S25 phosphorylation.

**Supplementary Figure 6. Extended respiratory, ultrastructural, metabolic and cell-cycle analyses downstream of DeSI1 activation. (A)** Maximal OCR and coupling efficiency were derived from the cell-count-normalized Seahorse Mito Stress Test in Figure 6A-B. Maximal OCR was calculated as the maximum FCCP-stimulated OCR minus the minimum OCR after rotenone+antimycin A and, for visualization, was expressed relative to the matched shDeSI1 value from the same experiment (shDeSI1 = 100%). Coupling efficiency was calculated as ATP-linked OCR divided by basal mitochondrial OCR × 100 and is shown as the actual percentage. Small open circles represent technical wells and larger filled circles represent biological-replicate means; bars show mean ± SEM across n = 3 independent biological experiments. Statistical analysis used a one-way ANOVA with independent experiment as the matched factor followed by two-sided Dunnett’s multiple-comparisons test versus shDeSI1. Technical wells were not used as independent statistical replicates. Only significant comparisons are indicated; *P < 0.05, **P < 0.01, ***P < 0.001.

**(B)** Representative transmission electron micrographs after acute (24 h) induction: parental U2OS, DeSI1 KO, KO reconstituted with DeSI1 WT, and KO reconstituted with DeSI1 S25E. Scale bars, 500 nm. Quantitative morphometry in Supplementary Figure 6C.

**(C)** Quantification of mitochondrial ultrastructure from TEM images of U2OS parental, DeSI1 KO, DeSI1 WT, and DeSI1(S25E) cells. Shown are cristae area fraction, matrix density index, matrix lucency fraction, and aspect ratio. Small shaded points represent individual mitochondrial profiles; open circles represent the summary value for each independent biological replicate (n = 4 biological replicates per genotype). Bars show mean ± SEM. For statistical analysis, measurements were first summarized at the biological replicate level, and replicate summaries were treated as the experimental unit. Three comparisons were prespecified: S25E vs WT, S25E vs KO, and Parental vs KO. Statistics shown on the graphs indicate nominal two-sided exact Mann–Whitney P values, with significant comparisons displayed (*P < 0.05). Holm-adjusted P values across the three prespecified comparisons within each morphometric endpoint are reported below. For cristae area fraction, nominally significant differences were detected for S25E vs WT, S25E vs KO, and Parental vs KO (each nominal P = 0.0286; each Holm-adjusted P = 0.0857). For matrix density index, S25E vs KO was nominally significant (P = 0.0286; Holm-adjusted P = 0.0857). For matrix lucency fraction, S25E vs KO was nominally significant (P = 0.0286; Holm-adjusted P = 0.0857). For aspect ratio, S25E vs WT was nominally significant (P = 0.0286; Holm-adjusted P = 0.0857).

**(D)** Genetic rescue of mitochondrial respiration. Basal mitochondrial OCR and proton leak were quantified in ShDeSI cells, ShDeSI cells treated with D1, parental cells treated with D1, and ShDeSI+D1 cells reconstituted with DeSI1 WT. OCR was normalized in each well to post-assay cell number, and technical wells were averaged within each independent experiment. Basal OCR is expressed relative to the matched ShDeSI condition within each experiment (ShDeSI = 100%); proton leak is shown in absolute cell-normalized units because the matched ShDeSI denominator was unstable for ratio normalization. Bars show mean ± SEM of n = 3 independent biological experiments; small shaded points indicate technical wells and open circles indicate biological-replicate means. Statistical analysis was performed by one-way repeated-measures ANOVA followed by two-sided Šidák correction across four prespecified comparisons. Only significant comparisons are shown; *P < 0.05, **P < 0.01.

**(E)** Experimental design for the metabolomic and lipidomic profiling in Figure 6C and Supplementary Figure 6F. RPE shDeSI1 cells, alone or with doxycycline-induced DeSI1 WT or S25E, were compared after 72 h starvation versus 8 h serum-stimulated G1 entry; polar metabolites and lipids were quantified by mass spectrometry (n = 6 per condition). Doxycycline was added after 48 h of starvation and maintained for 24 hours until serum release, so these samples report the acute state of the switch.

**(F)** Polar metabolomics. Heatmap shows the difference between S25E and the indicated reference line in the matched G1-versus-starvation response. Intensities were log2 transformed and median-normalized within each sample. G1 and serum-starved samples were paired by biological replicate, and values shown are mean differences across matched replicate responses. One ShDeSI G1 sample (replicate 2) was excluded from the metabolomics analysis based on the PCA/QC assessment; consequently, SE/ShDeSI comparisons contain five matched replicate blocks, whereas SE/WT comparisons contain six. Statistical significance was determined by two-sided one-sample t-test of the matched difference-in-differences against zero. *P<0.05, **P<0.01, ***P<0.001. CTP and UTP were not measured on this platform.

**(G)** Cell-cycle distribution across acute and chronic DeSI1 states. Top, stacked G1/S/G2-M distributions; middle, S-phase fraction; bottom, G1 fraction. Conditions 1–6 are as indicated. Mean ± SEM; n = 3 independent experiments; Welch two-sided t-test, uncorrected; *P < 0.05, **P < 0.01, ***P < 0.001; ns, not significant.

**Supplementary Figure 7. Chemical-genetic profiling and analysis of cytidine rescue during prolonged DeSI1 activation. (A)** Design of the chemical resistance assay. EF1α-GFP-marked U2OS parental or DeSI1 WT cells were mixed 1:1 with EF1α-mCherry-marked cyclin D1 T286A or DeSI1 S25E cells, seeded, incubated for 72 h with the indicated inhibitors, and the GFP:mCherry ratio determined by flow cytometry.

**(B)** Chemical resistance assay of DeSI1-dependent competitive fitness. Reporter signal for 26 compounds spanning six mechanistic classes, measured in two competition setups: D1 T286A expressing cells versus parental, and DeSI1(S25E) versus DeSI1 wild-type rescued cells. Values are normalized within each setup to the Sham control (= 1.0); n = 3 per condition. Colour encodes direction and magnitude (blue, reduced; maroon, increased; white, unchanged). Asterisks mark ordinary two-way ANOVA post-tests comparing each compound with Sham within each setup, separate families per setup, Holm–Šidák-adjusted (*q < 0.05, **q < 0.01, ***q < 0.001, ****q < 0.0001). See Supplementary Table 4 for compounds and final concentrations.

**(C)** Cell-normalized mitochondrial stress-test profiles following chronic DeSI1 activation and cytidine supplementation. RPE-hTERT shDeSI1 cells carrying doxycycline-inducible empty vector, DeSI1 WT, or DeSI1 S25E were induced for 7 days, with or without cytidine as indicated. All conditions received doxycycline. OCR was normalized in each well to the corresponding post-assay cell count and is shown as pmol O₂/min per 10,000 cells. Technical wells were averaged within each independent biological experiment before aggregation. Traces show mean ± SEM across n = 3 independent biological experiments at each measurement. Dashed vertical lines indicate sequential addition of oligomycin, FCCP, and rotenone+antimycin A. The trace is shown in absolute cell-normalized units and is not additionally normalized to shDeSI1.

**(D)** Secondary respiratory parameters during cytidine rescue of chronic S25E cells. Proton leak, coupling efficiency and spare respiratory capacity were derived from the same cell-count-normalized Seahorse XF Mito Stress Test experiments shown in Figure 6H. Proton leak was calculated as the minimum OCR after oligomycin minus the minimum OCR after rotenone+antimycin A; coupling efficiency as ATP-linked OCR divided by basal mitochondrial OCR × 100; and spare respiratory capacity as maximal OCR minus basal mitochondrial OCR. For visualization, proton leak is expressed relative to the matched shDeSI1 condition within each experiment (shDeSI1 = 100%), whereas coupling efficiency and spare respiratory capacity are shown in their native units. Bars show mean ± SEM and circles represent n = 3 independent biological experiments; technical wells were averaged within experiment before statistical analysis. WT and S25E cells ± cytidine were analyzed by two-factor repeated-measures ANOVA with genotype and cytidine as within-experiment factors and independent experiment as the matched factor. Interaction P values are shown as genotype × cytidine interaction P values. For spare respiratory capacity, the bracket denotes the single pre-specified S25E versus S25E + cytidine comparison, tested by paired two-tailed t-test on biological-replicate means.

**(E)** Representative transmission electron micrographs corresponding to Supplementary Figure 7F. Chronically induced (7 days) RPE-hTERT shDeSI1 cells carrying empty vector, DeSI1 WT or DeSI1 S25E, each ± cytidine. Fields were selected as those whose matrix lucency lay nearest the median of their condition. Images were acquired at 8,000× (0.83 nm per pixel); brightness and contrast were adjusted linearly and identically across all panels for display only, and all quantification in Supplementary Figure 7F was performed on unadjusted 16-bit images scaled by a single intensity window shared across the entire experiment. Scale bars, 100 nm.

**(F)** Quantification of mitochondrial ultrastructure from TEM images of RPE-hTERT shDeSI1 cells expressing EV, chronic DeSI1 WT or chronic DeSI1 S25E, with or without cytidine supplementation. Shown are matrix density index, matrix lucency fraction, and mitochondrial circularity. Small, shaded points represent individual mitochondrial profiles; open circles represent the summary value for each independent biological replicate (n = 3 biological replicates per condition). Bars show mean ± SEM. Cytidine-supplemented conditions are indicated by diagonal hatching. For statistical analysis, biological-replicate summaries were analyzed testing genotype, cytidine supplementation, and their interaction (metric ∼ genotype × cytidine). The shDeSI1+EV condition is shown as a reference but was not included in the factorial analysis. Two comparisons were prespecified: WT chronic versus S25E chronic and S25E chronic versus S25E + cytidine. Stars on the graphs denote nominal model-based P values (*P < 0.05, **P < 0.01, ***P < 0.001). Holm-adjusted P values across the two prespecified contrasts within each metric are reported below. For matrix density index, S25E differed from WT chronic (nominal P = 0.0013; Holm-adjusted P = 0.0026), and cytidine increased matrix density in S25E cells (nominal P = 0.0048; adjusted P = 0.0048). The genotype × cytidine interaction was significant (nominal P = 0.0013; Holm-adjusted across the three morphometric interaction tests P = 0.0040). For matrix lucency, WT chronic versus S25E chronic did not reach significance (P = 0.054), whereas cytidine significantly reduced matrix lucency in S25E cells (nominal P = 0.0041; adjusted P = 0.0083). The genotype × cytidine interaction was significant (nominal P = 0.0105; adjusted P = 0.0188). For circularity, WT chronic and S25E chronic did not differ significantly (P = 0.629), whereas cytidine significantly altered circularity in S25E cells (nominal P = 0.0054; adjusted P = 0.0109). The genotype × cytidine interaction was significant (nominal P = 0.0094; adjusted P = 0.0188).

**(G)** Chronic S25E growth deficit is cytidine-reversible. Cumulative cell number over 8 days of doxycycline induction of DeSI1 WT or S25E ± cytidine. Mean ± SEM; n = 3 independent experiments. Cells were seeded at 10,000 per well on day 0 and counted every 2 days. Lines are log2-linear fits drawn at the mean per-replicate slope *k*, in population doublings per day (per-replicate fit r² = 0.995–0.9996). Mean *k* ± SEM: WT 0.961 ± 0.012, S25E 0.798 ± 0.020, WT + cytidine 0.942 ± 0.019, S25E + cytidine 0.950 ± 0.011. S25E lowers the slope relative to WT (Δ*k* = −0.163 doublings day⁻¹, p = 0.0044) and cytidine reverses this (S25E + cytidine versus S25E, Δ*k* = +0.153, p = 0.0065), while cytidine leaves WT unchanged (Δ*k* = −0.019, p = 0.47). Genotype × cytidine interaction p = 0.0007 (two-way ANOVA on *k*). All pairwise comparisons are Welch’s t-tests on raw, uncorrected p values. **p < 0.01.

**Supplementary Figure 8. In vivo characterization of the DeSI1 S25E allele. (A)** Loss of DeSI1 reduces post-weaning body-weight gain. Body weight of DeSI1^+/+^ (WT), Desi1^S25E/S25E^ (S25E) and Desi1^−/−^ (KO) mice measured weekly from P21 to P84 (n = 12 per genotype of both sexes). Points show mean ± SEM. The overall genotype effect was estimated from a linear mixed-effects model (body weight ∼ genotype × age, with a random intercept per animal, age centred): KO versus WT, −2.10 g, p = 0.0005; S25E versus WT, −0.24 g, p = 0.62. Individual time points were compared by unpaired two-tailed Welch t-test without correction for multiple comparisons; asterisks denote KO versus WT (*p < 0.05, **p < 0.01, ***p < 0.001). S25E did not differ from WT at any time point (p = 0.16–0.82). Each animal is a biological replicate.

**(B)** Extended behavioural analysis. Top, within-animal longitudinal change in cerebellar composite score across the three timepoints in S25E mice only, stratified by sex; connected lines, individual animals; bold line, group mean; two-sided Wilcoxon signed-rank test. Bottom, balance-beam traversal time (s) at the same timepoints. Bars, mean ± SEM with individual animals shown; P values as indicated; n.s., not significant.

**(C)** Molecular-layer thickness (µm), measured as the mean length of perpendicular lines drawn across the molecular layer in the sections analysed in Figure 7C and collapsed to per-animal means as in Figure 7D. Bars, mean ± SEM with individual animals shown; Welch two-sided t-test, uncorrected; *P < 0.05, ***P < 0.001; ns, not significant.

**(D)** Cerebellar neurodegeneration. Representative silver-stained cerebellar sections from 6-month-old DeSI1 knock-out, wild-type and S25E mice. Inset, magnified view of argyrophilic degenerating profiles in S25E. Scale bar, 100 µm.

**(E)** Astrocyte labelling. Representative immunofluorescence of mid-sagittal cerebellar sections from wild-type, S25E and knock-out mice stained for GFAP, from the same multiplex panel as the calbindin-D28k images in Figure 7C. Scale bars, 50 µm.

**(F)** GFAP-positive area (%) quantified from (E) in QuPath by pixel classifier, scored on coded images and collapsed to per-animal means as in Figure 7D. Bars, mean ± SEM with individual animals shown; Welch two-sided t-test, uncorrected; *P < 0.05; ns, not significant. S25E exceeds knock-out but does not differ significantly from wild type, indicating that at this age the degeneration proceeds without a pronounced astrogliotic response.

**Supplementary Figure 9. Molecular characterization of sustained DeSI1 activation in vivo. (A)** Ponceau staining of the tissue panel in Figure 7E.

**(B)** Brain proteome, S25E versus knock-out. Volcano plot of whole-brain DIA-MS (n = 6 per genotype) for the S25E versus knock-out contrast, plotted and annotated as in Figure 7F. Comparing the phosphomimetic allele against the null rather than against wild type separates gain-of-function effects of S25E from any consequence of reduced wild-type DeSI1 activity.

**(C)** Ponceau staining of the brain panel in Supplementary Figure 9D.

**(D)** Brain immunoblots. Whole-brain lysates from 6-month-old knock-out, wild-type and S25E mice (M, male; F, female) immunoblotted for switch readouts (pDeSI1 S25, total DeSI1), stabilized substrates (ACC2, MTHFSD, SEC63, SEC61G, SFXN2, PEX14), the de novo pyrimidine enzyme DHODH, respiratory and inner-membrane proteins (ATP5PB, NDUFS4, UQCRC2, SDHD, SLC25A5). Tubulin and GAPDH, loading controls; Ponceau staining is shown in Supplementary Figure 9C. Densitometry of the substrate and respiratory panels is shown in Figure 7G.

**(E)** Brain lipid classes and cardiolipin composition in DeSI1(S25E) mice. Heatmap shows log₂ lipid-class abundance differences between S25E and WT brain (S25E n = 5, WT n = 5; WT3 excluded a priori based on cross-platform QC). For each mouse, lipid-class abundance was calculated by summing the per-sample median-normalized linear abundances of de-duplicated annotated species within each class, followed by log₂ transformation. Statistics were calculated using two-sided Welch t-tests with mouse as the unit of replication. Asterisks indicate nominal significance (*P < 0.05, **P < 0.01, ***P < 0.001); P values for the nine predefined primary lipid classes were additionally corrected within the S25E/WT comparison using the Benjamini–Hochberg procedure. Cardiolipin saturation bins are secondary compositional analyses and were not included in the primary class-level correction. Less-unsaturated cardiolipin species (≤4 double bonds) were preferentially reduced in S25E brain, whereas more highly unsaturated cardiolipin species were relatively preserved.

**(F)** Nucleotide-pathway enzymes in the brain proteome, grouped by position relative to the CTP-consuming step: the CTP-consuming transferases (demand), the de novo pyrimidine enzymes, and the salvage and interconversion enzymes (supply). Columns show the S25E/wild-type and S25E/knock-out contrasts; cells render protein log₂ fold change. n = 6 mice per genotype; Welch two-sided t-test on log₂ protein quant, raw uncorrected P; *P < 0.05, **P < 0.01, ***P < 0.001; n.s., not significant. The rate-limiting cytidylyltransferase PCYT1A is elevated against both controls, while CTPS1, the isoform induced when demand rises, is unchanged in every genotype, as are CAD and UMPS at the entry to the pathway and the salvage and interconversion enzymes. DHODH and CTPS2 are reduced in S25E against both controls.

## Supplementary Tables

**Supplementary Table 1. Quantitative DeSI1 interactome identified by IP-MS.** Complete protein-level quantitative data from the DeSI1 interactome analysis underlying Figure 3A and Supplementary Figure 3B–C. HEK293T cells expressing FLAG-tagged DeSI1 WT or the phosphomimetic DeSI1(S25E) mutant were subjected to anti-FLAG immunoprecipitation followed by LC–MS/MS (n = 3 independent biological replicates per condition). The table contains protein identifiers, replicate-level quantitative measurements and derived comparison statistics. Fold changes are expressed as S25E relative to WT; therefore, positive log₂ fold-change values indicate preferential association with DeSI1(S25E), whereas negative values indicate preferential association with wild-type DeSI1. The complete ranked dataset was used for the interactome volcano plot and for selection of the directionally enriched protein sets used in the Gene Ontology and membrane-insertion-propensity analyses.

**Supplementary Table 2. His-ubiquitin proteomic screen.** Complete protein-level quantitative data from the denaturing His-ubiquitin proteomic analysis underlying Figure 3C and Supplementary Figure 3D–F. HEK293F cells expressing 6×His-tagged ubiquitin together with empty vector (EV), DeSI1(S25E), or the catalytically inactive DeSI1(S25E/C108S) mutant were subjected to Ni-NTA enrichment under denaturing conditions followed by LC–MS/MS. The table contains protein identifiers, replicate-level quantitative measurements and derived statistics for the indicated pairwise comparisons. Because enrichment was performed under denaturing conditions, the measured signal represents covalently ubiquitin-conjugated protein recovered in the His-Ub fraction rather than non-covalent association with DeSI1.

For the principal S25E-versus-EV analysis, a common no-MG132 analysis universe was defined by requiring quantification in at least two of three biological replicates in each of the EV, S25E and S25E/C108S conditions, yielding 4,531 gene-resolved proteins. Proteins were ranked by the S25E-versus-EV *t* statistic; increasingly negative values indicate lower His-Ub recovery in S25E-expressing cells and thus define the candidate DeSI1-dependent reduction-in-ubiquitylation direction. The lowest 20% of this ranking (907 proteins) was used for the targeted membrane-compartment enrichment and membrane-insertion-propensity analyses. Comparison with DeSI1(S25E/C108S) was used to assess dependence on DeSI1 catalytic activity.

**Supplementary Table 3. Peptide binding of DeSI1 variants.** Sequences (N→C), length (n), net charge at pH 7.4 (z₇.₄), GRAVY hydropathy index and mean hydrophobic moment (µH) for the cognate COX4 mitochondrial presequence, the FIS1 tail-anchored transmembrane domain with its basic C-terminal extension, and a polar COX4-scaffold control peptide, with fluorescence-polarization Kᴅ values (µM; 95% confidence interval in parentheses) for DeSI1 WT, S25E and S25E/C108S at 150 mM NaCl. n.b., no saturable binding, with the ΔmP at the highest titration point given in parentheses. Three biological replicates throughout.

**Supplementary Table 4. Compounds and concentrations used in chemical competition profiling.**

