## Supplementary_Data for "Cyclin D-CDK4/6 couple proliferation to membrane and mitochondrial protein supply through a DeSI1 phospho-switch"

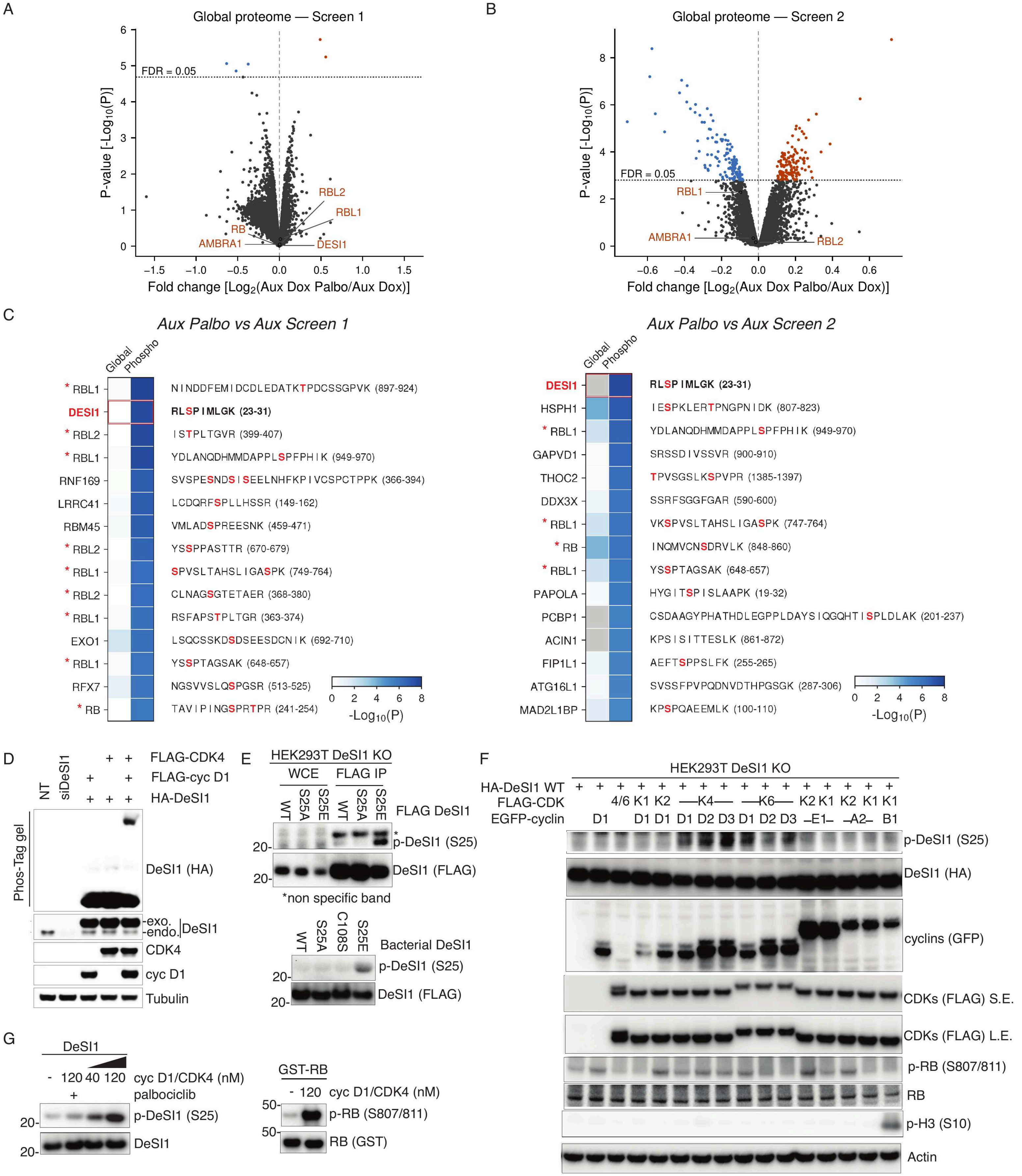


Kaisari et al., Supplementary Figure 1

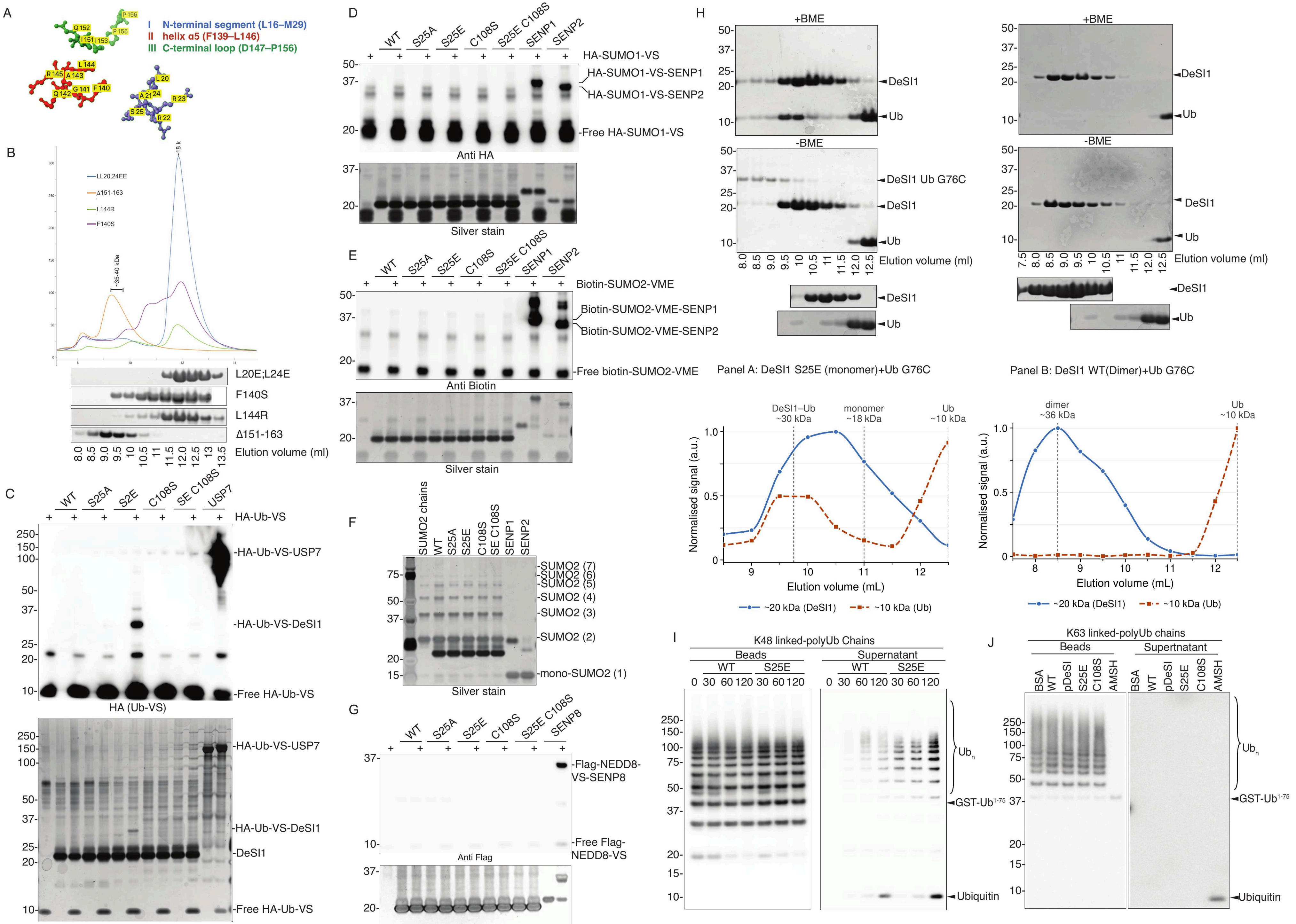


Kaisari et al., Supplementary Figure 2

A

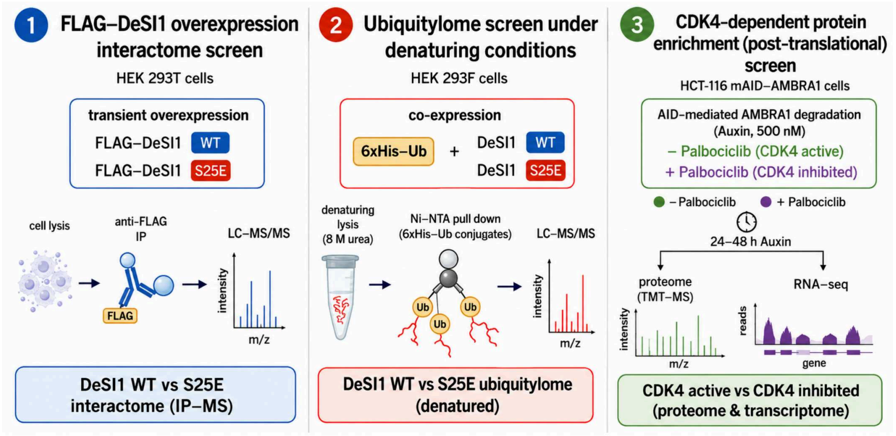


B

*WT vs. S25E DeSI1:*

| Functional Module | Proteins |
|---|---|
| **Ubiquitin-dependent Protein Catabolic Process** | |
| BAG6 Complex | ASNA1, BAG6, UBL4A |
| Ubiquilins | UBQLN1, UBQLN4, UBQLN2 |
| VCP Complex | VCP, NPL4, UFD1 |
| Proteasome Subunits | PSMA2, PSMB4, PSMB3<br>PSMD7, PSMA3, PSMC3<br>PSMA1, PSMB2 |
| Ubiquitin | UBA52, UBC |
| **Cellular Response to Stress** | |
| Response to Heat | HSPA2, HSPA6, HSPA1L<br>HSPA8, HSPA1A, HSPA4L |
| Response to Unfolded Protein | DNAJB2, DNAJC10 |
| Hydrogen Peroxide Metabolic Process | PRDX1, PRDX4 |
| **Vesicle Traffic** | |
| Rab GTPases | RAB5C, RAB7A, RAB11A |
| Trafficking Regulators | SDCBP, WLS, SQSTM1<br>DEGS1, PI4K2A, BCAP31<br>LRRC59 |

C

*S25E vs. WT DeSI1:*

| Functional Module | Proteins |
|---|---|
| **Mitochondrial Matrix** | |
| | FASTKD5, PYCR1, HIBCH<br>UQCRC2, OAT, NDUFS1<br>NDUFS2, ALDH1L2, ECHS1 |
| **Mitochondrial Membrane** | |
| | MIRO1, SLC25A20, SLC25A5<br>SLC25A6, SLC25A4, SLC25A22<br>TIM44, SFXN1, AGK<br>AIFM1 |
| **Endoplasmic Reticulum Membrane** | |
| | SEC63, SRPRA, ESYT2<br>SEC61A1, MOGS, ALG6<br>MGAT2, COMT, SPTLC1<br>HSD17B12, HSD17B4, ACSL3<br>NAA50, STT3A, RPN1<br>HACD3, SMPD4 |
| **Coated Vesicle Membrane** | |
| | AP1G1, AP2B1, USO1<br>VPS26A, RHOG, GAPVD1<br>MEMO1, PKN2, PDLIM5<br>KLC1, ALG2, SEC22B<br>COPA, COPG1, COPG2<br>NSF |
| **Bounding Membrane of Organelle** | |
| | PEX3, PXMP2, ACAA1<br>ZADH2, ADGRL2, TMEM43<br>TMEM165, CLIC4, CNNM3<br>TRABD, CDIPT, PTGES2<br>PDXDC1, ATP1A1, ATP2A2<br>ATP2B1, TM9SF3, SLC16A1 |

D

*Membrane Enrichment of S25E vs EV Ubiquitylome:*

| Functional module | Fold | P | BH q |
|---|---|---|---|
| Outer Membrane (33/100) | 1.65× | 0.0014 | 0.00294 |
| Endoplasmic Reticulum Membrane (74/307) | 1.20× | 0.0397 | 0.0432 |
| Endoplasmic Reticulum Exit Site (9/20) | 2.25× | 0.00989 | 0.0148 |
| Coated Vesicle Membrane (23/81) | 1.42× | 0.0432 | 0.0432 |
| Vesicle Membrane (98/351) | 1.39× | 0.00013 | 0.000781 |
| Golgi Membrane (46/152) | 1.51× | 0.00147 | 0.00294 |

E

*Catalytic validation within the ubiquitylome hits*

| Functional Module | Proteins |
|---|---|
| **Mitochondrial Matrix (31)** | |
| | ALDH1B1, CCNB1, CLPX, EXD2, FDX1<br>FECH, GCDH, HARS2, IBA57, KARS1<br>LARS2, MRPL16, MRPL27, MRPL47, MRPS11<br>MRPS15, MRPS2, MRPS28, NDUFB8, NDUFS8<br>NME4, PARG, PC, PIN4, PTCD1<br>PTPN1, RIDA, RPUSD4, SUCLG1, TST<br>UQCC2 |
| **Mitochondrial Membrane (30)** | |
| | ABCB7, ACSL1, ARMCX3, ATAD1, ATP5MK<br>BID, CISD1, CYB5A, ETFDH, HK1<br>HK2, LGALS3, MAVS, MFN2, MTOR<br>MTX2, NDUFAF4, NDUFC2, NME3, PLA2G4A<br>PPOX, PTRH2, QTRT2, SAMM50, SLC25A10<br>SPIRE1, TIGAR, TIMM23, UQCC1, VDAC3 |
| **Endoplasmic Reticulum (40)** | |
| | ALDH3A2, COPB2, COPG2, DHCR24, DHCR7<br>DHRS4, DNAJB2, EMC2, GALNT2, MAGT1<br>NRAS, PIGK, RAB14, RHEB, SAR1A<br>SBF1, SCFD1, SEC24A, SEC24C, SEC24D<br>SEC62, SEC63, SORT1, SPAST, SPCS3<br>SPTLC2, STX17, TBL2, TECR, TEX264<br>TMED2, TMED4, TMED5, TOR1AIP2, UBA5<br>UBQLN4, VAPA, VTI1B, WASL, YIPF5 |
| **Vesicle Membrane (55)** | |
| | AP2A2, AP3M1, AP3S1, ARPC2, ATP6V0A1<br>ATP6V1A, CCDC93, CHMP1A, CHMP3, CHMP4A<br>COMMD5, COMMD9, DMXL2, DNM2, EHD3<br>EPN1, GGA2, GNPNAT1, HIP1R, HTT<br>INPP5F, IQGAP2, LAMTOR3, MYO1B, MYO1C<br>NCK1, NECAP1, NUMB, PDLIM4, RAB12<br>RAB23, RABGEF1, RAP1B, SLC12A2, SNF8<br>SNX1, SNX9, STK10, STX12, STX4<br>STXBP2, TBC1D24, TOM1, TOM1L1, TRAF6<br>TSG101, VAC14, VPS11, VPS37B, VPS45<br>VPS52, VTA1, WASH3P, WASHC3, YKT6 |
| **Bounding Membrane of Organelle (9)** | |
| | ACBD3, ARFGEF1, CHP1, COG6, GOLGA5<br>MGAT4C, PRKCI, QPCTL, TRAPPC4 |

*Ubiquitylome membrane insertion propensity:*

F

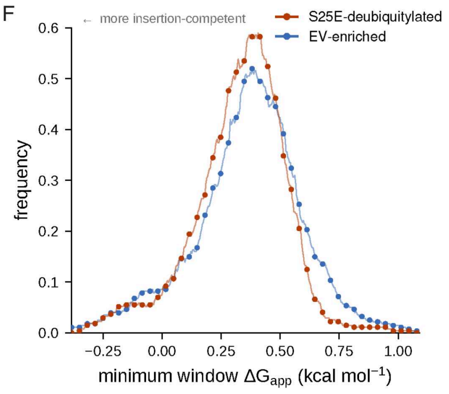


*CDK4-related proteome dominant membrane insertion propensity:*

G

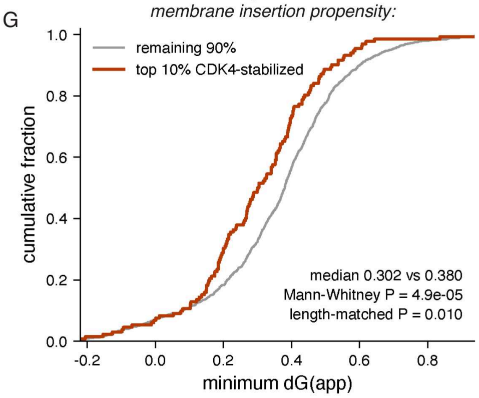


Kaisari et al., Supplementary Figure 3

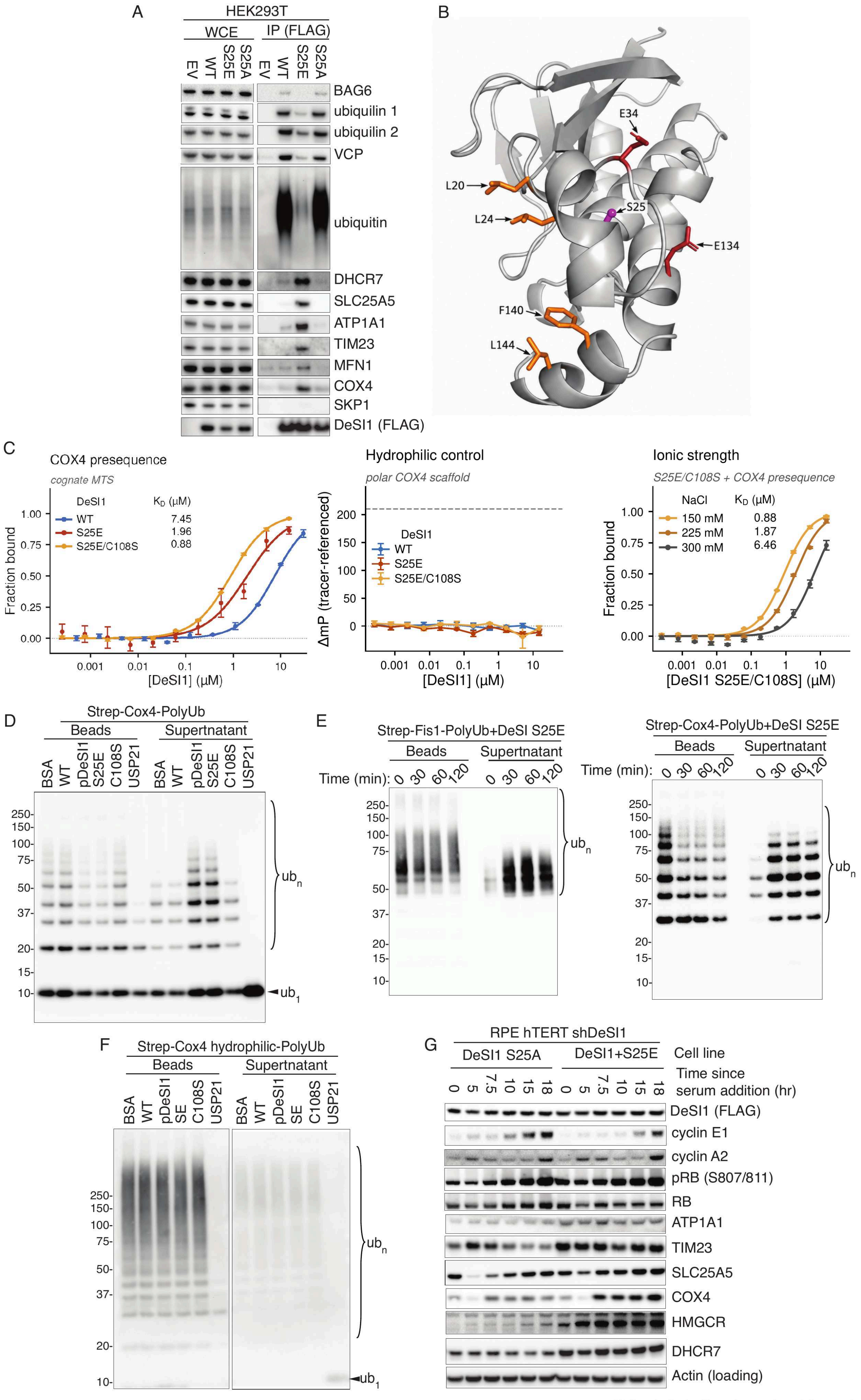


Kaisari et al., Supplementary Figure 4

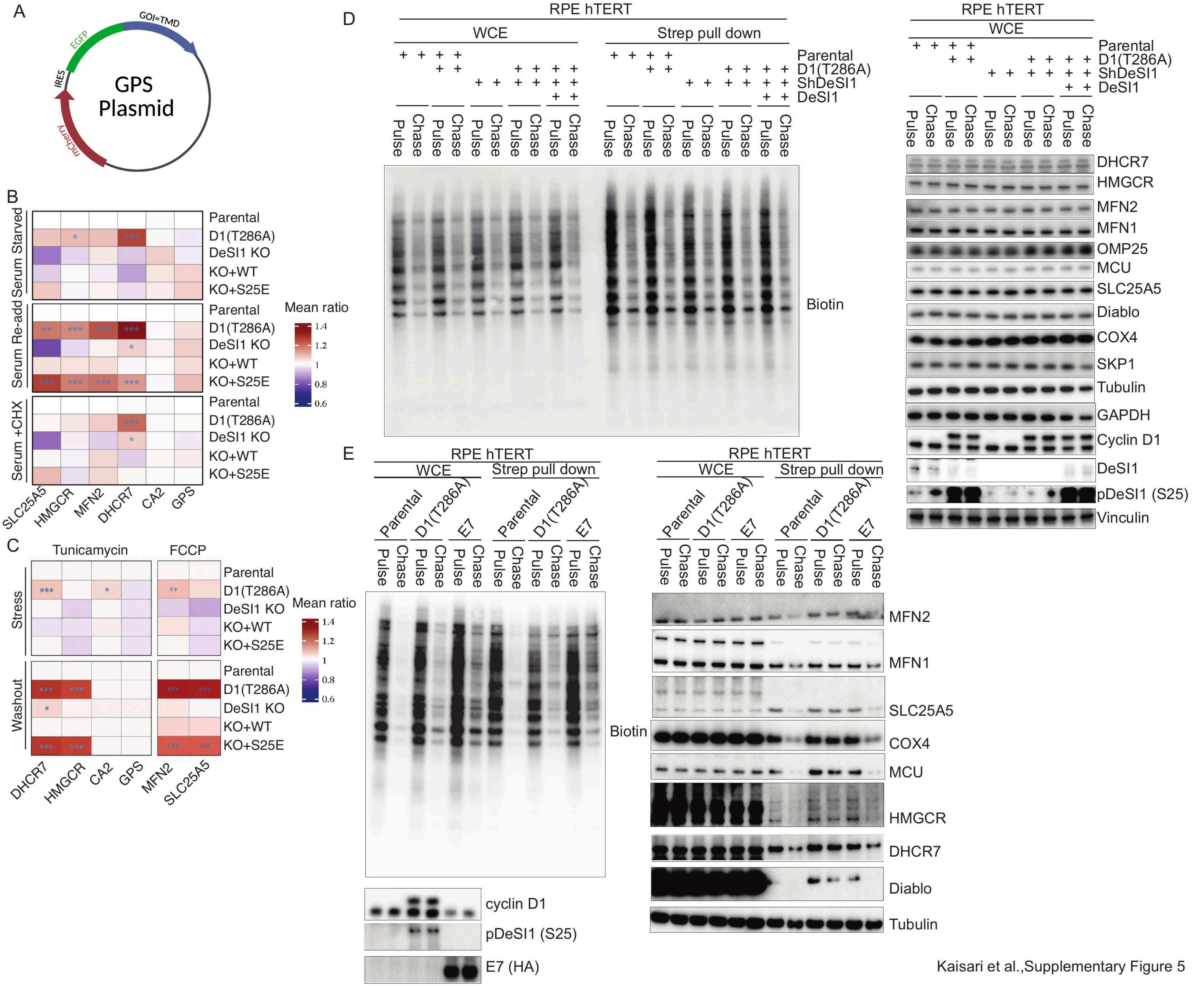


Kaisari et al.,Supplementary Figure 5

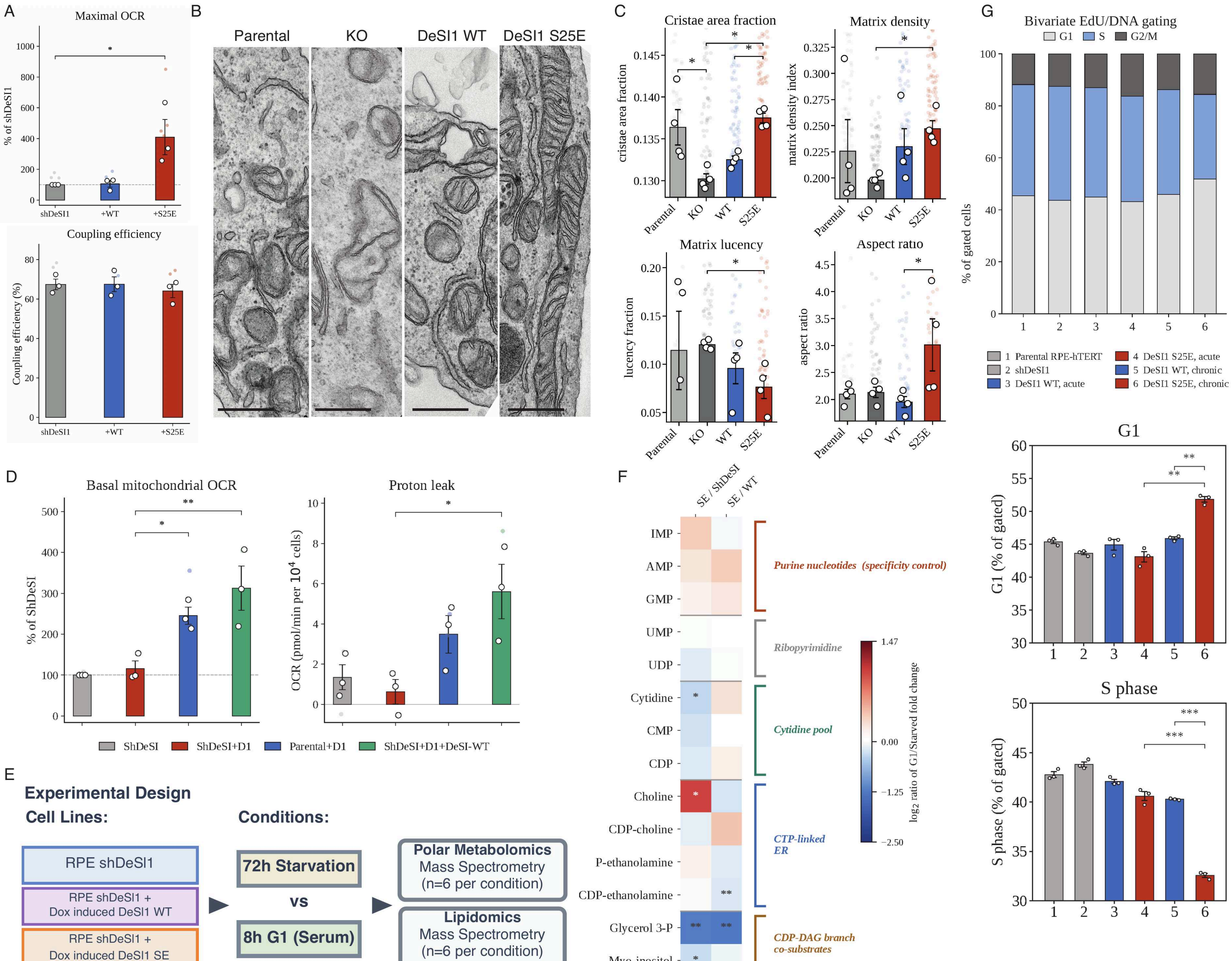


Kaisari et al.,Supplementary Figure 6

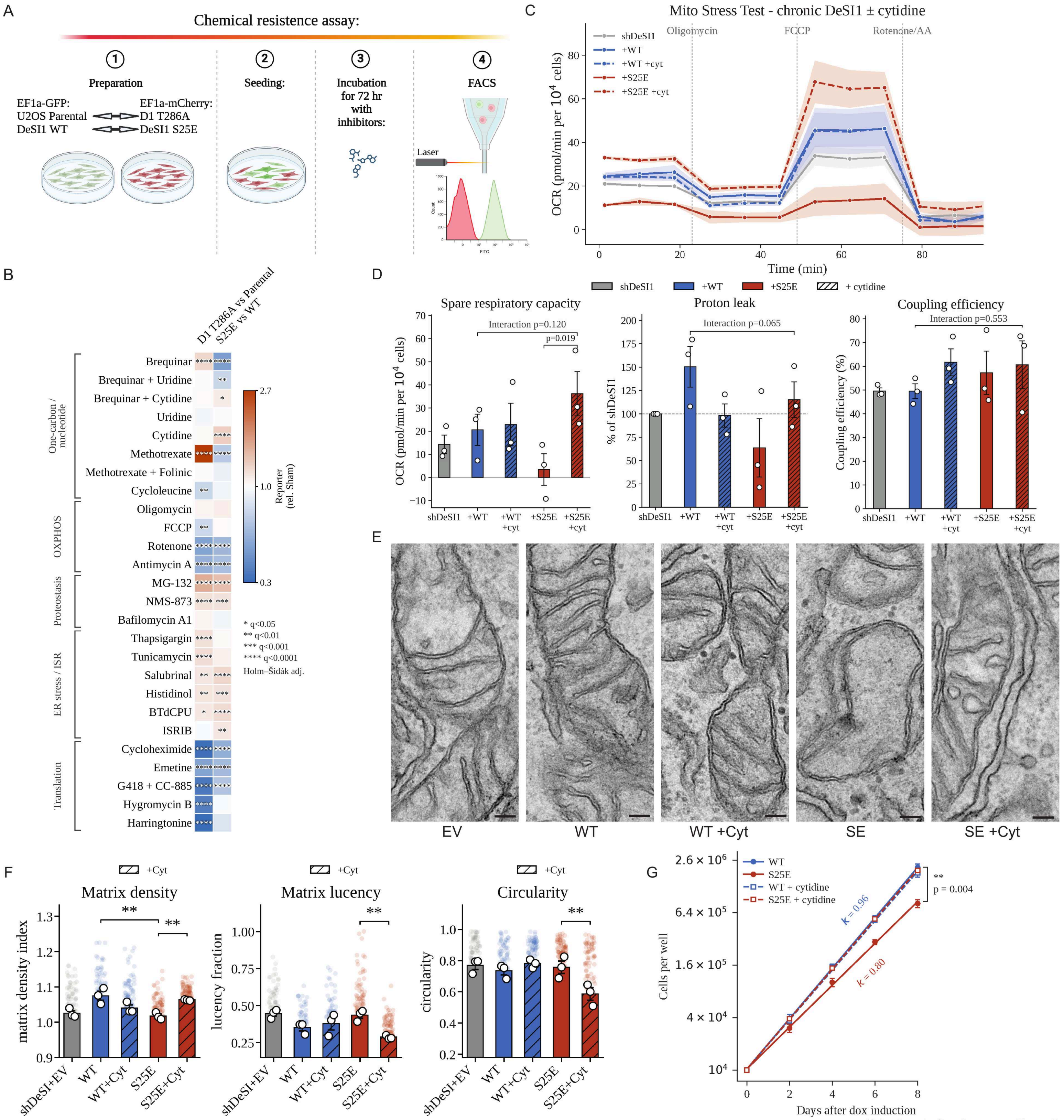


Kaisari et al., Supplementary Figure 7

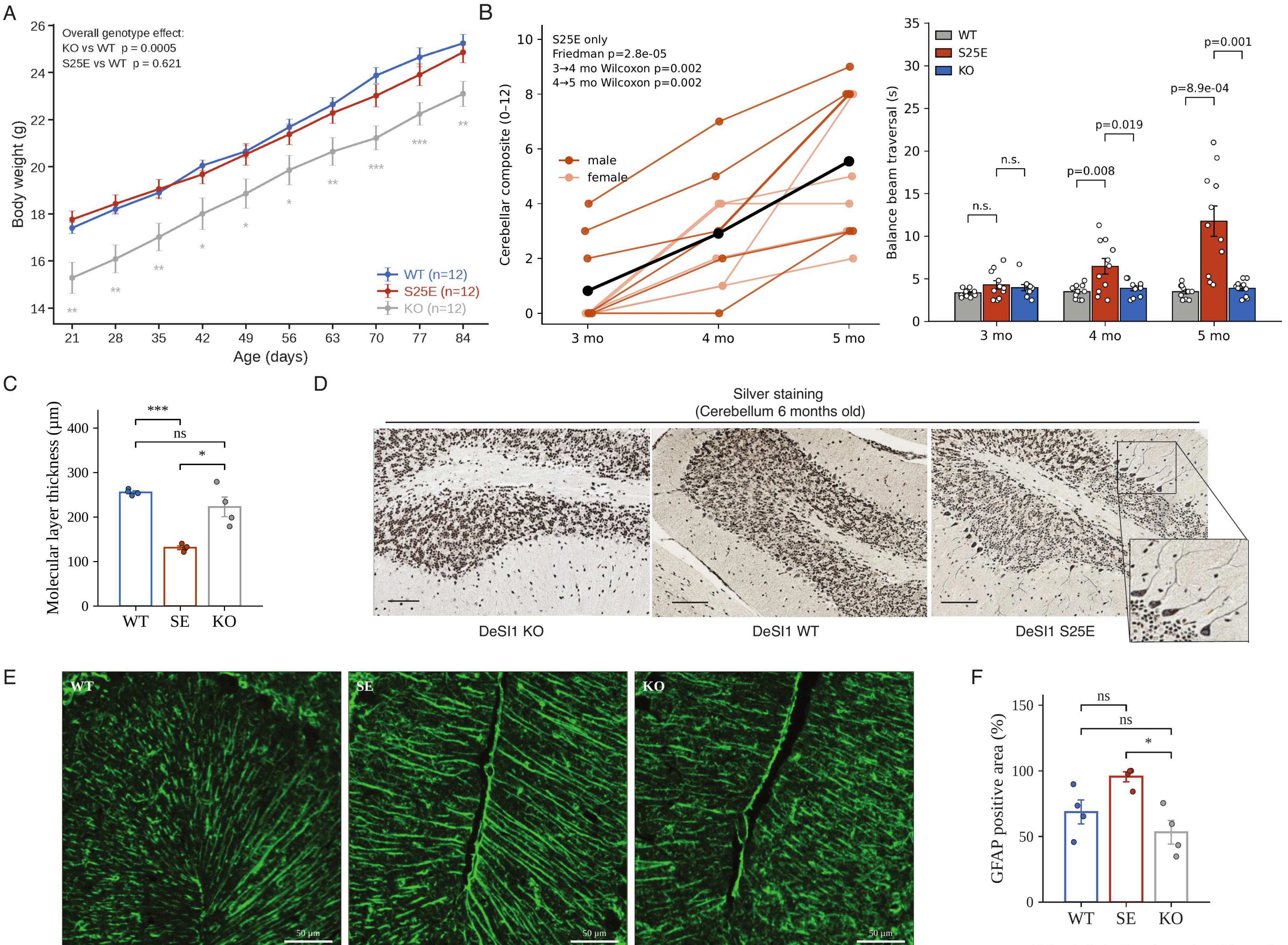


Kaisari et al., Supplementary Figure 8

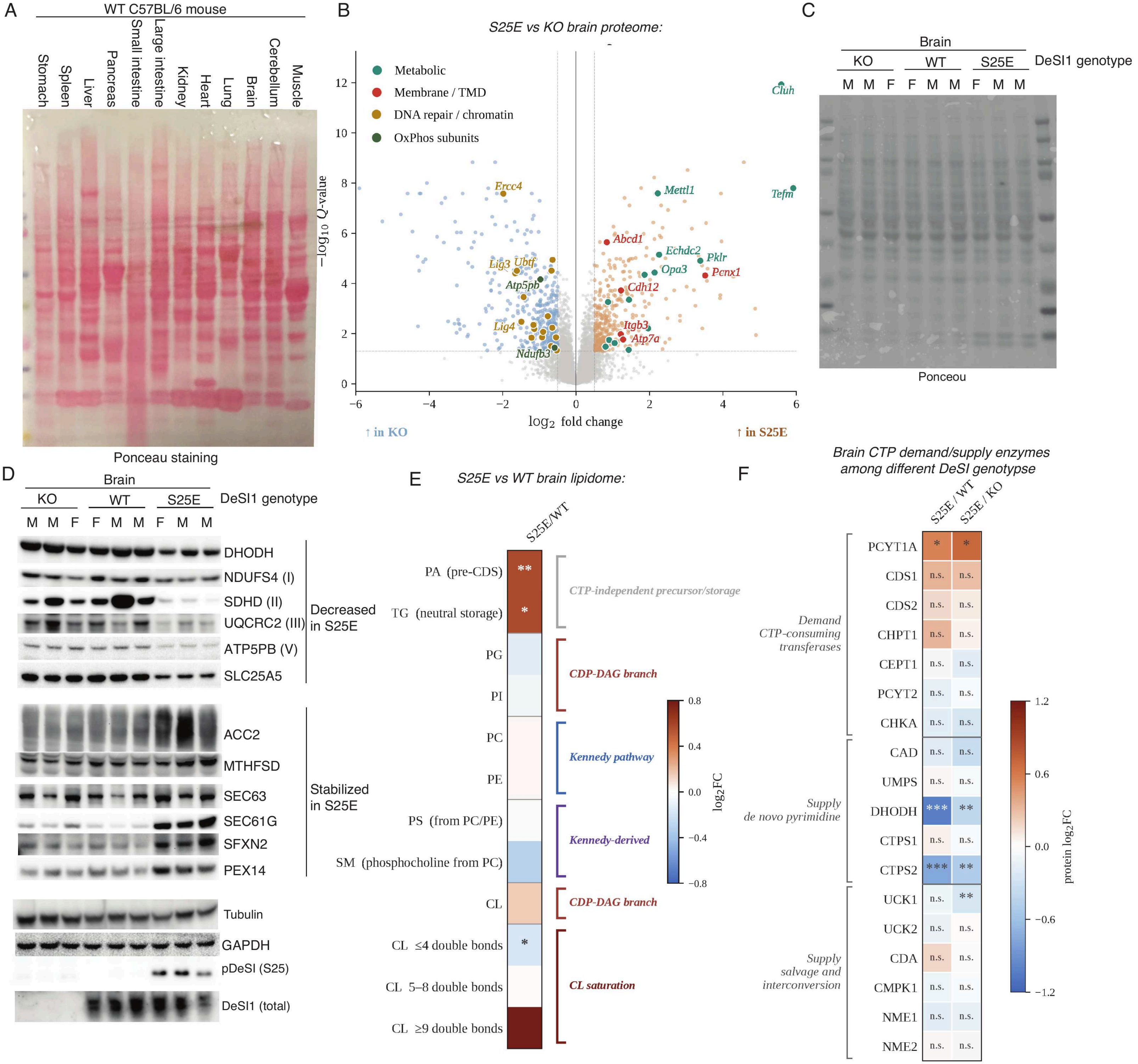


Kaisari et al., Supplementary Figure 9

| Peptide | Sequence (N→C) | $n$ | $z_{7.4}$ | GRAVY | $\mu_H$ | $K_D$ (μM) WT | $K_D$ (μM) S25E | $K_D$ (μM) S25E/C108S |
|---|---|---|---|---|---|---|---|---|
| COX4 presequence<br>*cognate MTS* | MLSLRQSIRFFKPATRTLSSSRYLL | 25 | +4 | -0.036 | 0.398 | 7.45<br>(5.74–9.67) | 1.96<br>(0.81–4.73) | 0.88<br>(0.83–0.95) |
| FIS1 TMD-CTE<br>*tail-anchored TMD + basic extension* | LGAVAGAAILGGAILVGGMRRKKF | 24 | +3 | +0.971 | 0.053 | 9.10<br>(1.43–57.83) | 1.59<br>(0.50–5.01) | 1.08<br>(0.69–1.68) |
| Hydrophilic control<br>*polar COX4 scaffold* | MSSNTQSNSFFKPATNTSNSSNSNS | 25 | +0 | -1.200 | 0.158 | n.b.<br>(ΔmP +3.3) | n.b.<br>(ΔmP -10.7) | n.b.<br>(ΔmP -17.0) |

Kaisari et al., Supplementary Table 3
